# SpatialAgent: An autonomous AI agent for spatial biology

**DOI:** 10.1101/2025.04.03.646459

**Authors:** Hanchen Wang, Yichun He, Paula P. Coelho, Matthew Bucci, Abbas Nazir, Bob Chen, Linh T. Trinh, Serena Zhang, Ziyu Lu, Kexin Huang, Vineethkrishna Chandrasekar, Douglas C. Chung, Minsheng Hao, Ana Carolina Leote, Yongju Lee, Bo Li, Tianyu Liu, Jin Liu, Romain Lopez, Lucas Tawaun, Mingyu Ma, Nikita Makarov, Lisa McGinnis, Linna Peng, Stephen Ra, Gabriele Scalia, Avtar Singh, Liming Tao, Masatoshi Uehara, Chenyu Wang, Runmin Wei, Ryan Copping, Orit Rozenblatt-Rosen, Jure Leskovec, Aviv Regev

## Abstract

Advances in AI are transforming scientific discovery, yet spatial biology, a field that deciphers the molecular organization within tissues, remains constrained by labor-intensive workflows. Here, we present SpatialAgent, an autonomous AI agent for spatial biology research. SpatialAgent couples large language models with a Plan-Act-Conclude architecture, dynamic tool and skill retrieval, multimodal interpretation, and verification modules that audit generated claims. It supports the full discovery loop, from gene-panel design and multimodal annotation to trajectory inference, cell-cell communication analysis, imputation, and hypothesis generation. Across human and mouse brain, heart, tonsil, colon, and prostate datasets, SpatialAgent outperformed established computational baselines and matched or surpassed expert scientists in key tasks. In open-ended case studies, it recovered known tissue organization and generated spatially grounded hypotheses. In a prospective mouse prostate cancer Xenium study, it designed a compact 100-gene add-on panel that profiled 4.2 million cells across 21 samples, improved cell-type and malignant-state prediction, and captured spatially structured tumor and microenvironment programs. SpatialAgent establishes a framework for autonomous and collaborative discovery in spatial biology.

## Introduction

Modern AI is becoming a key tool for scientific discovery (1). Recently, Large language models (LLMs) (2) have shown promises in accelerating scientific progress by enabling reasoning, planning, and tool integration, through autonomous LLM-equipped agents (3). Autonomous agents are AI systems that iteratively perceive, plan, and act, allowing them to dynamically adapt to tasks with minimal human intervention. This makes them particularly valuable in more open-ended scientific research, where they integrate pre-existing knowledge (*e.g.*, via LLMs and retrieval-augmented generation) with analytical actions (in connected tools), and iterative reasoning. In this they are distinct from both prior automated pipelines, which had to be hard-coded and hence could not adapt to the dynamic needs of new data analysis, and human-driven analysis, where a seasoned data scientist must iteratively engage with analysis tools, results, and knowledge, including through LLM prompting.

Spatial genomics, a rapidly growing field, offers a particularly compelling case in point. Spatial genomics aims to decipher how biomolecules and cells are spatially organized within tissues in homeostasis and disease (4). As an emerging area, driven by novel lab technologies that generate complex high dimensional data, spatial genomics relies on computational methods that are quite fragmented, require extensive human intervention and do not yet generalize well across diverse datasets and biological contexts. Moreover, users have to reason over multiple analysis approaches and complex concepts spanning molecules, cells and multicellular interactions, that require deep biological knowledge across many cell types, gene programs and interaction mechanisms to interpret. Thus, in practice, most studies demand substantial commitment from expert analysts and time-consuming work. We reasoned that LLM-driven autonomous agents could help drive innovative work in spatial genomics, both accelerating routines and helping make new discoveries. Indeed, recent studies have demonstrated the promise of agentic AI systems in scientific workflows, from iterative hypothesis refinement to tool-using automation across molecular dynamics and biology (5–14).

Here, we introduce SpatialAgent, an LLM agent that autonomously navigates the spatial biology gamut, from experimental design to multimodal analysis and data-driven hypothesis generation. Unlike established computational approaches in spatial genomics, which rely on predefined workflows and rigid models, SpatialAgent employs adaptive reasoning and dynamic tool integration, allowing it to adapt to new datasets, tissue types, and biological questions. It processes multimodal inputs, incorporates external databases, and supports human-in-the-loop interactions, enabling both fully automated and collaborative discovery. Unlike general-purpose biomedical agents such as Biomni (9) that target broad workflows, SpatialAgent is specialized for spatial omics, integrating spatial analysis tools, image-aware interpretation, and tasks such as niche annotation, deconvolution, and spatial cell-cell communication. While some recent systems emphasize scientist-in-the-loop interaction (10), our main quantitative results are reported under fully autonomous runs, with co-pilot analyses presented separately. We evaluate SpatialAgent across multiple datasets spanning diverse human and mouse tissues (brain, heart, tonsil, colon and prostate) across both healthy and disease states (*e.g.*, colitis and cancer) and across a broad range of spatial genomics tasks, including experimental design (gene panel design), spatial analysis (cell and tissue annotation), data integration, and interpretation (cell-cell communication, trajectory inference, imputation) for generating novel biological hypotheses. SpatialAgent outperformed existing computational baselines, matched or surpassed human experts in key biological tasks, and accelerated scientific workflows. Beyond efficiency, SpatialAgent enhanced the interpretability of its AI-driven discoveries, bridging the gap between automation and human intuition and legacy knowledge. SpatialAgent sets a new standard for autonomous and collaborative research in spatial biology, with broader implications for biomedical research and precision medicine.

## Results

### SpatialAgent uses a Plan-Act-Conclude architecture with dynamic tool use and verification

We developed SpatialAgent, an autonomous AI agent for spatial biology that supports the full discovery loop across experiment, observation, and hypothesis stages (**Fig.** 1a). SpatialAgent supports end-to-end analyses including gene panel design, multimodal annotation of cell types and tissue niches, and hypothesis generation with explicit verification, for example by integrating imputation, cell-cell communication analysis, trajectory inference, and biological interpretation. The system is intended to generalize across organs, tissues, and cross-species settings, while supporting both fully autonomous analysis and iterative scientist follow-up.

**Figure 1:**
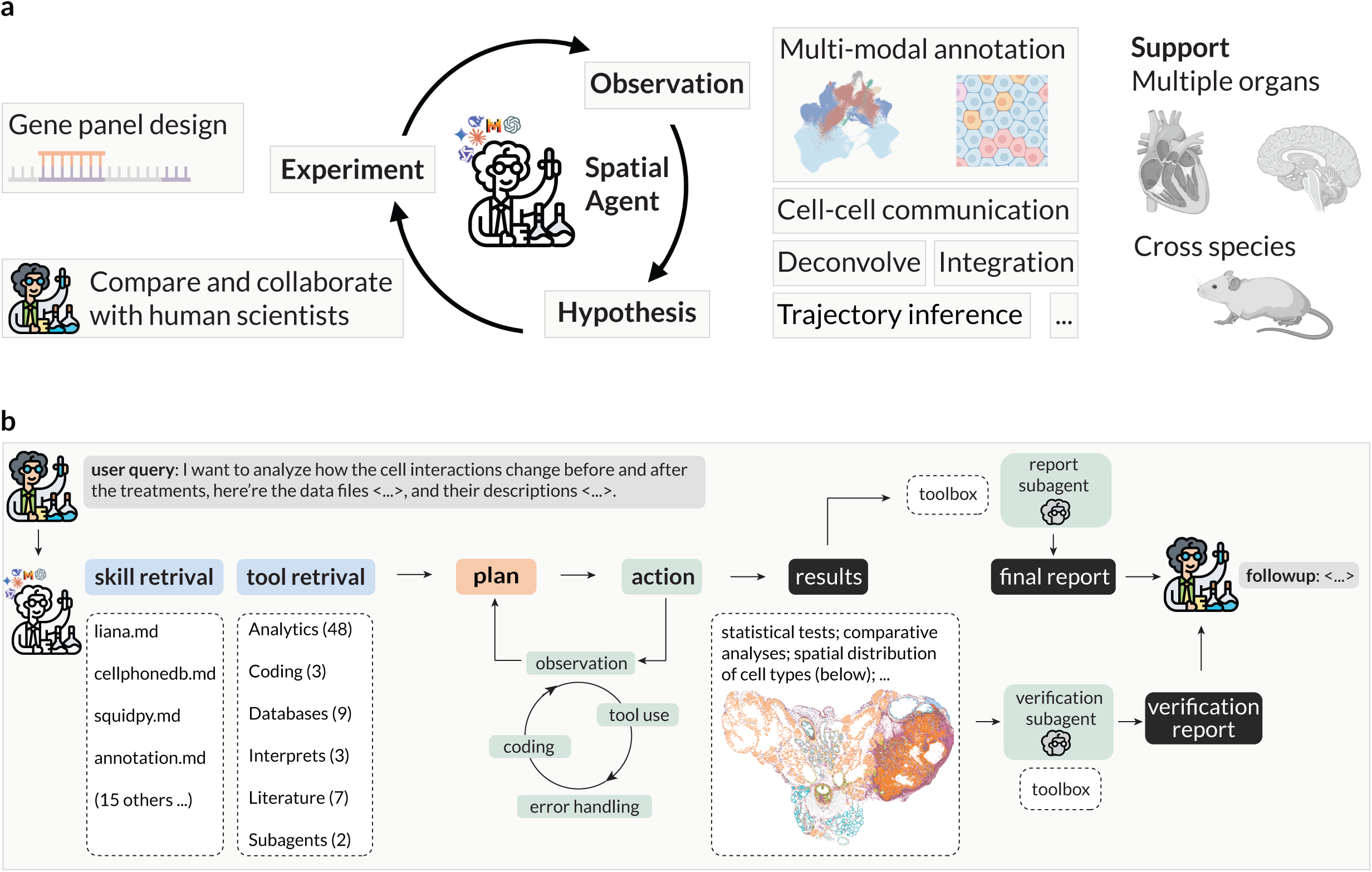
Overview and modular design of SpatialAgent. **(a)** Overview of SpatialAgent. **(b)** Plan-Act-Conclude architecture. In Plan, SpatialAgent retrieves relevant skills and tools, decomposes the query into concrete steps, and updates the plan through iterative observations. In Act, it executes code, database queries, and specialized tools to generate outputs and artifacts. Persistent state stores intermediate results, decisions, failures, and artifacts across steps and turns. In Conclude, SpatialAgent synthesizes outputs into a final response or report and can invoke reporting and verification subagents before returning the final answer.

At the system level, SpatialAgent links LLM reasoning to reliable execution through three coordinated stages: **Plan**, **Act**, and **Conclude** (**Fig.**1b, **Methods**). In **Plan**, the agent decomposes the user query into executable steps, retrieves task-relevant tools and skills, and updates the analysis strategy as new observations arrive. In **Act**, it grounds reasoning in external resources by running literature and database searches, invoking specialized tools, and writing, executing, and debugging analysis code. In **Conclude**, SpatialAgent synthesizes outputs into a final response or report, and can invoke reporting and verification modules to check key numerical and visual claims before returning the final answer.

To preserve flexibility without overloading context, SpatialAgent retrieves a task-relevant subset of tools for each query, typically 5 to 20 tools from a broader toolbox, while keeping a small core set always available for code execution, tool inspection, and literature or web search. It further relies on a library of reusable skills that encode common multi-step procedures in spatial biology, such as panel design, spatial domain detection, condition-aware cell-cell communication analysis, and trajectory inference. These skills act as structured guidance rather than fixed scripts, allowing the same skill to instantiate different action chains depending on the user request and intermediate observations. SpatialAgent also maintains persistence and auditability across analyses through restored context, saved artifacts, and structured execution records (**Methods**).

To improve reliability in open-ended scientific settings, SpatialAgent augments primary analysis with downstream reporting and verification modules that summarize workflows, reference generated artifacts, and audit numerical and visual claims. SpatialAgent uses a configurable LLM backend that supports both API-based and locally served models. We benchmarked multiple backends and decoding settings to quantify trade-offs in accuracy, latency, and cost, and separately ablated the contribution of the skill library (Supplementary Information 1.4).

Across these ablations, SpatialAgent’s performance was shaped by three interacting factors: model capability, decoding strategy, and domain guidance (Extended Data Figs. 1–4; Supplementary 1.4). Backend comparisons showed that stronger frontier models generally improved scientific reasoning and task completion, whereas locally served open-weight models provided a practical deployment path with greater local control but lower reasoning performance (Extended Data Figs. 1 and 3). Decoding analyses showed that temperature and reasoning effort affected both final metrics and execution behavior, including tool use, constraint following, and completion of multi-step analyses, with no single setting optimal across all models and tasks (Extended Data Fig. 2). Skill ablations showed that the skill library provides an additional reliability layer: its effects were modest in simpler panel-design settings, became more pronounced for larger or more constrained panels, and were strongest in cell-type annotation, where skills improved balanced accuracy, macro-F1, coverage, and ARI (Extended Data Fig. 4). These gains suggest that skills reduce common agent failure modes, including incomplete workflows, label collapse, and poor recovery of rare classes. Based on these ablations, unless otherwise noted, we used Claude Sonnet 4.5 with temperature 0.5 for the main experiments. We next evaluated SpatialAgent across benchmark tasks, open-ended case studies, and wet-lab validation (below).

### SpatialAgent outperformed prior methods and strengthened human-designed gene panels

To test whether SpatialAgent can design practical gene panels end to end, we benchmarked it on Visium spatial transcriptomics data from human dorsolateral prefrontal cortex (DLPFC), spanning 12 tissue sections from three adult donors (15). For each panel size (50, 100, or 150 genes), SpatialAgent either designed from scratch or refined an existing panel provided by a scientist. In both settings, SpatialAgent retrieved the same skill but it instantiated different action chains depending on the user’s request and the intermediate evidence it observes. For a panel designed from scratch, it typically expanded the search over reference datasets, markers, curated gene sets, and relevant literature before scoring candidates; for a refinement request, it started by diagnosing gaps in the input panel and then selectively added, removed, or re-weighted genes. In both, it iteratively scored candidates, checked expression in the target tissue, and returned a ranked panel with per-gene scores and provenance (**Fig.** 2a).

**Figure 2:**
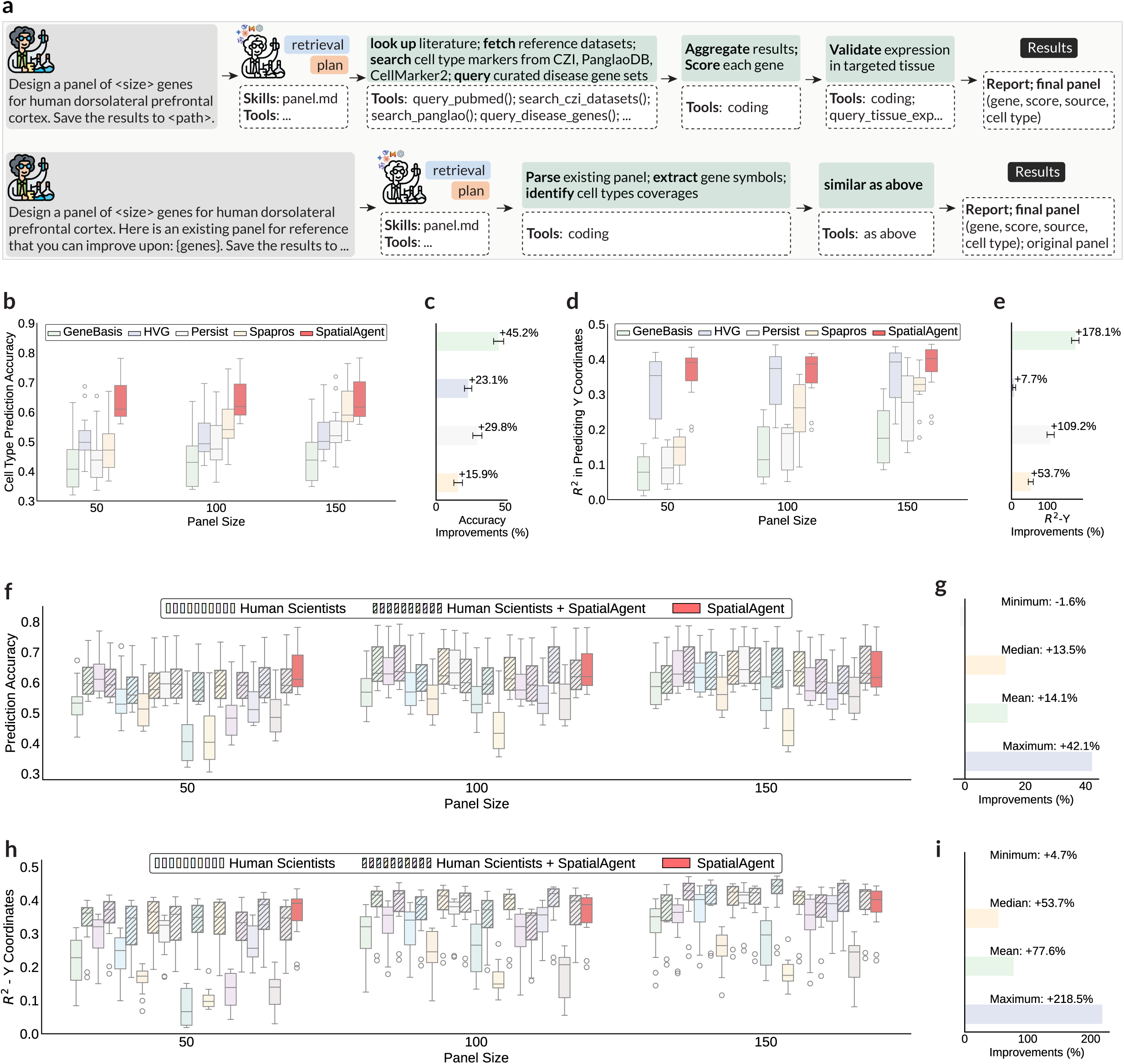
Autonomous gene panel design. **(a)** Illustrative step-wise autonomous workflow for designing a gene panel for human dorsolateral prefrontal cortex (DLPFC). Given a query to design a panel from scratch (top) or refine an existing one (bottom), SpatialAgent retrieves the same skill, then flexibly composes tailored action chains to meet the user’s request by gathering evidence (literature, reference datasets, markers, curated gene sets), scoring candidates, checking tissue expression, and returning a ranked panel with per-gene scores and provenance. **(b–e)** SpatialAgent outperforms established computational baselines for cell-type and spatial coordinate prediction. Absolute performance (b,d, y axis) and relative improvement (c,e, x axis, average across samples and panel sizes) for cell type (b,c) and spatial coordinate (d,e) predictions when designing 50, 100, or 150-gene panels based on 12 DLPFC samples, using SpatialAgent or four other computational pipelines. Box plots across 10 runs with medians, interquartile ranges, and 1.5× interquartile ranges; circles: outliers. **(f–i)** Comparison to human scientists and agent–scientist collaboration. Absolute performance (f,h, y axis) and relative improvement (g,i, x axis, average across samples and panel sizes) for cell type (f,g) and spatial coordinate (h,i) predictions for 50, 100, or 150-gene panels designed based on 12 DLPFC samples, using SpatialAgent (red), human expert (clear bars), agent-refined human designs (hashed bars). Box plots as in (b-e).

Across panel sizes from 50 to 150 genes, SpatialAgent’s panels consistently outperformed those from established computational pipelines (16–19) in terms of cell-type prediction accuracy (**Fig.** 2b,c, 15.9-45.2% improvement) and spatial coordinate prediction (**Fig.** 2d,e), with *R*^2^ gains of up to 178.1%. Thus, SpatialAgent autonomously designs gene sets that capture biological variance and spatial organization better than standard approaches.

Human-designed panels varied widely because scientists adopted distinct workflows (reference-only, literature-only, multi-dataset integration, etc.) and in some cases mismatched previous designs (**Extended Data Table** 1). Across 30 matched comparisons (10 human scientists on 3 panel sizes), SpatialAgent outperformed scientist-designed panels in 26/30 (86.7%) comparisons for cell-type accuracy and 29/30 (96.7%) for y-coordinate prediction (R-squared), with only 1/30 (3.3%) cases where one scientist exceeded SpatialAgent on both metrics (**Fig.** 2f–i). When SpatialAgent was used to refine each scientist’s panel, the refined panels almost always improved over the originals, improving 27/30 (90.0%) comparisons for accuracy and 29/30 (96.7%) for y-coordinate prediction. Refinement also helps improve the agent’s original designs: agent-refined scientist panels outperformed the agent-only design in 10/30 cases (33.3%) for accuracy and 16/30 cases (53.3%) for y-coordinate prediction, consistent with complementary strengths in which scientist priors provide a useful starting point and the agent’s evidence aggregation and scoring enhance the final panel.

### SpatialAgent selects spatially informative genes with low cost and latency

We next examined how SpatialAgent prioritizes genes, how well its panels preserve biological spatial organization (cortical-layer structure), and how its runtime and cost compare with computational baselines and scientists. In the “assign importance scores” step, SpatialAgent explicitly grounds each gene’s importance in a combination of evidence sources rather than a single heuristic. For example, it justifies a gene through canonical marker status and function (SLC17A7), cross-database agreement (PVALB), layer or cell-type specificity (SATB2), agreement with an existing scientist panel (MEIS2), tissue-expression support (GAD1), or literature and disease relevance (INPP5D) (**Fig.** 3a). This produces gene-level rationales that make the selection criteria transparent to humans and easier to audit, as well as helps the agent produce robust results even when any single resource is incomplete or inconsistent.

We then asked whether the selected genes preserve the laminar organization that defines the DLPFC. Using the 500-gene panel, SpatialAgent maintains a clear layer-associated structure in the spot embedding, visually resembling the embedding obtained from the full transcriptome more closely than a representative baseline (**Fig.** 3b). Across multiple quantitative measures of layer separation and coherence, SpatialAgent’s 500-gene panels are consistently among the best-performing approaches (**Fig.** 3c), only exceeded by the full set of genes, thus indicating that the panel captures both cell-identity variation and spatially structured signals that reflect cortical architecture. Moreover, the top-ranked genes selected by SpatialAgent include well-established neuronal markers (SLC17A7, PVALB, and GAD2), each showing spatially structured expression across the tissue section (**Fig.** 3d). In contrast, the top selected genes from Spapros (PCDH15, ATP1A2, and NKAIN3) show weaker spatial enrichment (**Fig.** 3d). Together, these observations suggest that SpatialAgent’s scoring favors genes that are simultaneously informative for cell identity and spatial organization, while remaining interpretable through explicit provenance and rationale.

Importantly, SpatialAgent is efficient, completing in about 12.2 minutes, compared to 3.8 hours for Spapros and 4.0 hours median scientist time (**Fig.** 3f). Estimated costs show a similar pattern: SpatialAgent is about $1.8 per run, compared to $123.2 for Spapros and $212.8 median scientist labor cost under our assumptions (**Fig.** 3g, **Methods**). These efficiency gains make it feasible to run multiple design–evaluate–refine cycles and to incorporate scientist feedback without turning panel design into a days-long bottleneck.

### SpatialAgent enhanced cell type and tissue niche annotation

Accurate annotation of spatially localized cells and tissue niches remains a major bottleneck in spatial biology, particularly when it requires integrating molecular measurements with spatial organization and anatomical context. SpatialAgent addresses this through a structured flow (**Fig.** 4a): given spatial molecular data and an anatomical reference image, it preprocesses the dataset, transfers annotations from an scRNA-seq reference, annotates cell types, and then segments the tissue into niches and annotates those niches with anatomical guidance and iterative refinement. We evaluated this capability on a high-resolution atlas of the developing human heart that combines scRNA-seq and MERFISH measurements across six samples collected at 9 to 16 post-conception weeks (142,946 scRNA-seq profiles and more than 1.5 million MERFISH-detected cells) (20). For cell-type annotation, we compared SpatialAgent with five automated baselines (21–24), the general purpose agent Biomni (9), and eight human experts (**Extended Data Table** 2), using the author labels as the reference (**Fig.** 4b,d; **Methods**). For tissue-niche annotation, we compared SpatialAgent with the same eight experts using the author annotations as the ground truth, as there are no automated tools for this task to our best knowledge (**Fig.** 4c,e; **Methods**).

Overall, SpatialAgent produced consistent, high-quality cell-type annotations that closely aligned with the author labels and achieved the strongest aggregate performance across accuracy, precision, F1, and adjusted Rand index (**Fig.** 4b-d), with an overall accuracy of 0.853 (confusion-matrix: **Supplementary Figs.** 17-19, raw annotations: **Supplementary Figs.** 11-12). SpatialAgent outperformed the five automated baselines (overall accuracy: PopV 0.262; scTab 0.157; Biomni 0.394; GPT-CellType 0.811; SpatialAgent 0.853), mostly because several baselines exhibit strong label-collapse biases. For example, PopV assigned many non-VSMC classes to VSMCs, scTab over-assigned immune labels, and Biomni was skewed toward endothelial cell annotations. While GPTCellType had relatively high overall accuracy, it mis-assigned 98.1% of epicardial, 85.5% of immune, and 96.9% of neuronal cells as fibroblasts. In contrast, SpatialAgent’s errors are more structured: it cleanly recovers cardiomyocytes (99.1% correct), fibroblasts (94.8%), immune (92.1%), and neuronal (95.3%) cells, while most remaining inaccuracies were between cells from closely-related stromal and vascular subsets (pericytes 94.8% predicted as endothelial; VSMCs 32.9% predicted as endothelial; endothelial cells 29.2% predicted as fibroblasts). A notable residual challenge was in epicardial cell annotation, where epicardial cells were frequently assigned fibroblast labels (82.2%) rather than epicardial (14.5%), consistent with difficulty in resolving tissue-specific epicardial states under limited MERFISH gene panels. Human experts showed a wide performance range (accuracy 0.459–0.834), reflecting differences in expertise and annotation strategy (**Extended Data Table** 2), and their errors concentrated in similar boundary cases, especially among vascular and stromal lineages (endothelial, pericyte, and VSMC).

For tissue-niche annotation, SpatialAgent produced anatomically coherent niche maps that were outperformed most human experts and were comparable to the top human performers (aggregate metrics: **Fig.** 44c,e, confusion-matrix: Supplementary Fig. 19-20, raw annotations: Supplementary Fig. 13-16)). There was substantial variability in nice annotations between human experts (accuracy range 0.108–0.873), whereas SpatialAgent achieved 0.661 accuracy. SpatialAgent was particularly reliable for the major cardiac chambers, correctly identifying the right atrium (96.0%), right ventricle (94.1%), and left atrium (87.7%), while the left ventricle (65.8%) was more often confused with adjacent structures such as flow tracts (31.4%). The dominant residual errors concentrated in thin or anatomically complex regions, especially the conduction system, flow tracts, and valves: the conduction system was frequently assigned as flow tracts (42.8%), flow tracts were split between conduction system (42.8%) and valves (50.9%), and valves were often assigned to right ventricle (38.5%). Notably, these regions were also challenging for human experts. For example, even the best-performing expert correctly identified conduction system in only 37.8% of cases. This suggests that ambiguity at boundaries and limited marker support drive most remaining disagreement, rather than gross anatomical misinterpretation by the agent or the human experts.

### SpatialAgent executes analyses with integrated verification and reporting for human tonsil

Beyond standardized tasks, we evaluated SpatialAgent in an open-ended, hypothesis-generation setting on a spatial transcriptomics CosMx dataset generated in-house from healthy human tonsil (25) (**Fig.** 5), a secondary lymphoid organ with stereotyped B-cell follicles and interfollicular T-cell zones. This well-characterized architecture provides a stringent test: the agent should recover established organization while surfacing additional, testable spatial patterns. We tasked SpatialAgent with three complementary analyses requiring distinct computational strategies: spatial neighborhood detection (**Fig.** 5a), inference of B-cell maturation trajectories (**Fig.** 5b), and reference-guided gene imputation to expand transcript coverage (**Fig.** 5c). For each task, the main agent retrieved the relevant skill module, constructed and executed a multi-step plan, and produced intermediate artifacts, such as spatial maps, graphs, and marker trends. These were synthesized by a report subagent into an accessible narrative, while an independent verification subagent audited the claims, highlighted ambiguities, and recommended robustness checks.

**Figure 3:**
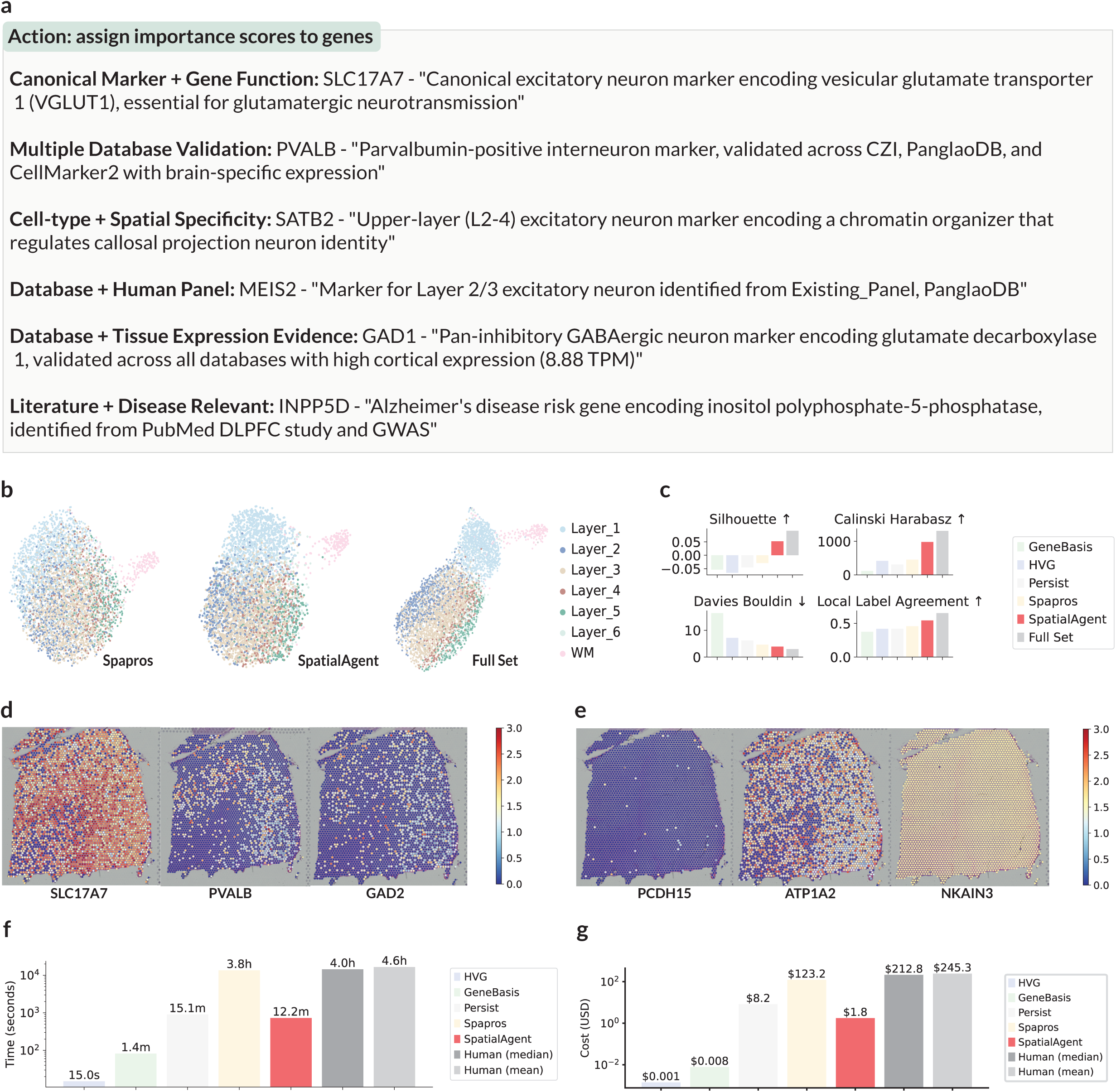
Reasoning, performance, and efficiency of SpatialAgent in gene panel design. **(a)** Evidence used by SpatialAgent to assign gene importance scores in the “assign importance scores to genes” step. **(b-e)** SpatialAgent selected genes have strong spatial patterns. (**b**) UMAP representation of DLPFC spatial spots (dots) colored by cortical layer based on 500-gene panels selected by Spapros (left), SpatialAgent (middle), or the full gene set (∼21,000 genes) (right).. **(c)** Cortical-layer separation metrics (label on top) for 500-gene panels from each method or the full gene set (x axis, color legend). Arrows: preferred direction. **(d,e)** Spatial expression patterns of the top three genes selected by SpatialAgent (d) and Spapros (e). **(f,g)** SpatialAgent efficiency. Runtime (f; y axis, log scale) and estimated cost (g; y axis, log scale) for each method (x axis). Human time is self-reported (median/mean) and the labor cost assumes an annual salary of $100,000 USD (8 working hours per weekday) (**Methods**).

**Figure 4:**
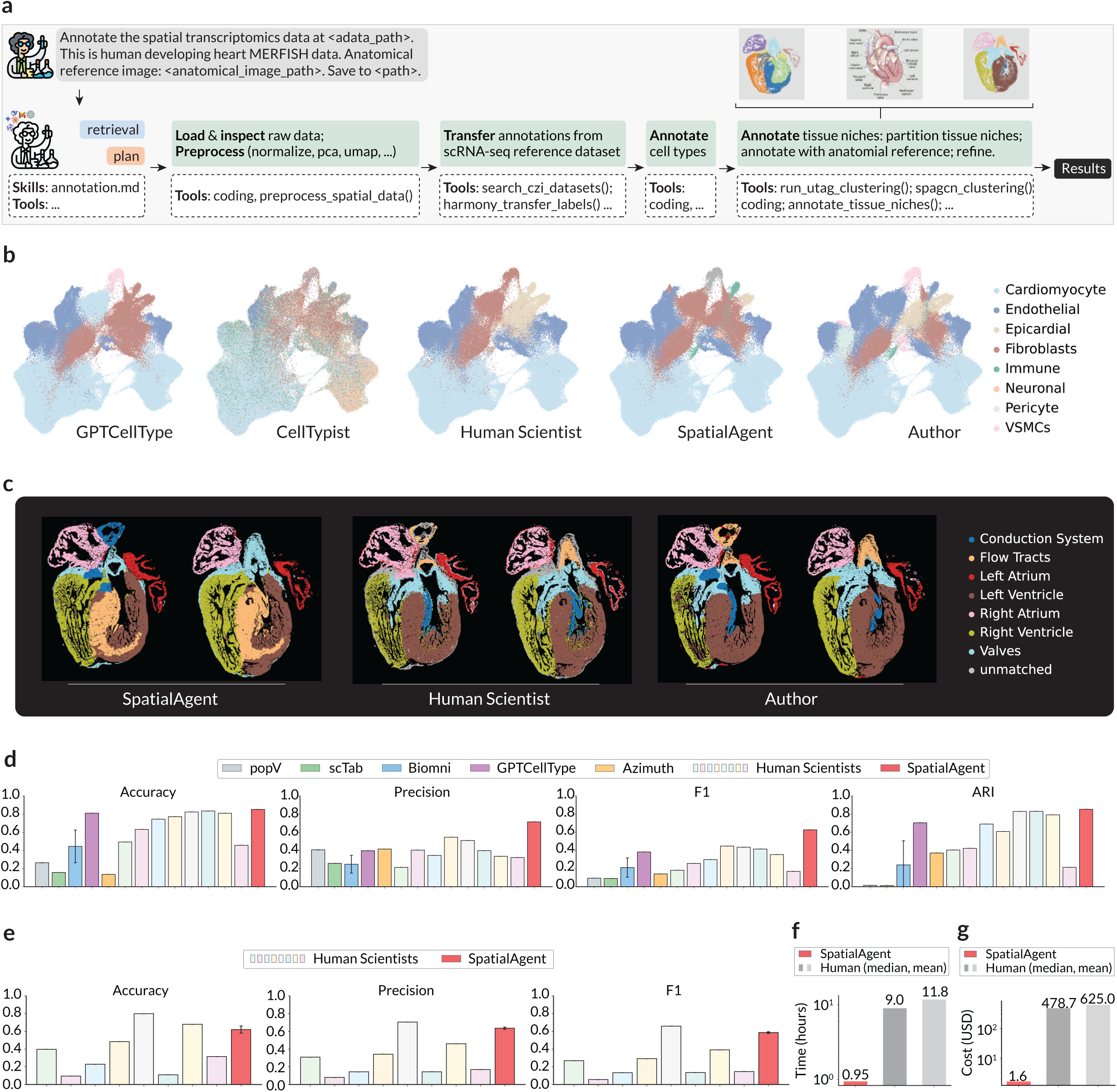
Cell type and tissue niche annotation by SpatialAgent. **(a)** Illustrative workflow. Given spatial data and an anatomical reference image, SpatialAgent retrieves the relevant skills/tools, inspects and preprocesses the data, transfers labels from scRNA-seq reference datasets, annotates cell types, then partitions and annotates tissue niches with anatomical guidance and refinement. **(b-e)** Cell type and niche annotation performance. (**b**) UMAP representation of cell profiles (dots) colored by eight major cell-type annotations from each computational pipeline, a representative human scientist, SpatialAgent, and original study. **(c)** Tissue section colored by tissue-niche annotation of seven major niches produced by SpatialAgent, a human scientist, and the original study. Unmatched: regions with no correspondence to author annotations. **(d,e)** Cell-type (d) and tissue niche (e) annotation quality metrics (y axis) with each tested approach (x axis). **(f, g)** SpatialAgent efficiency. Estimated time (left, y axis) and cost (right, y axis) for SpatialAgent and human scientists.

**Figure 5:**
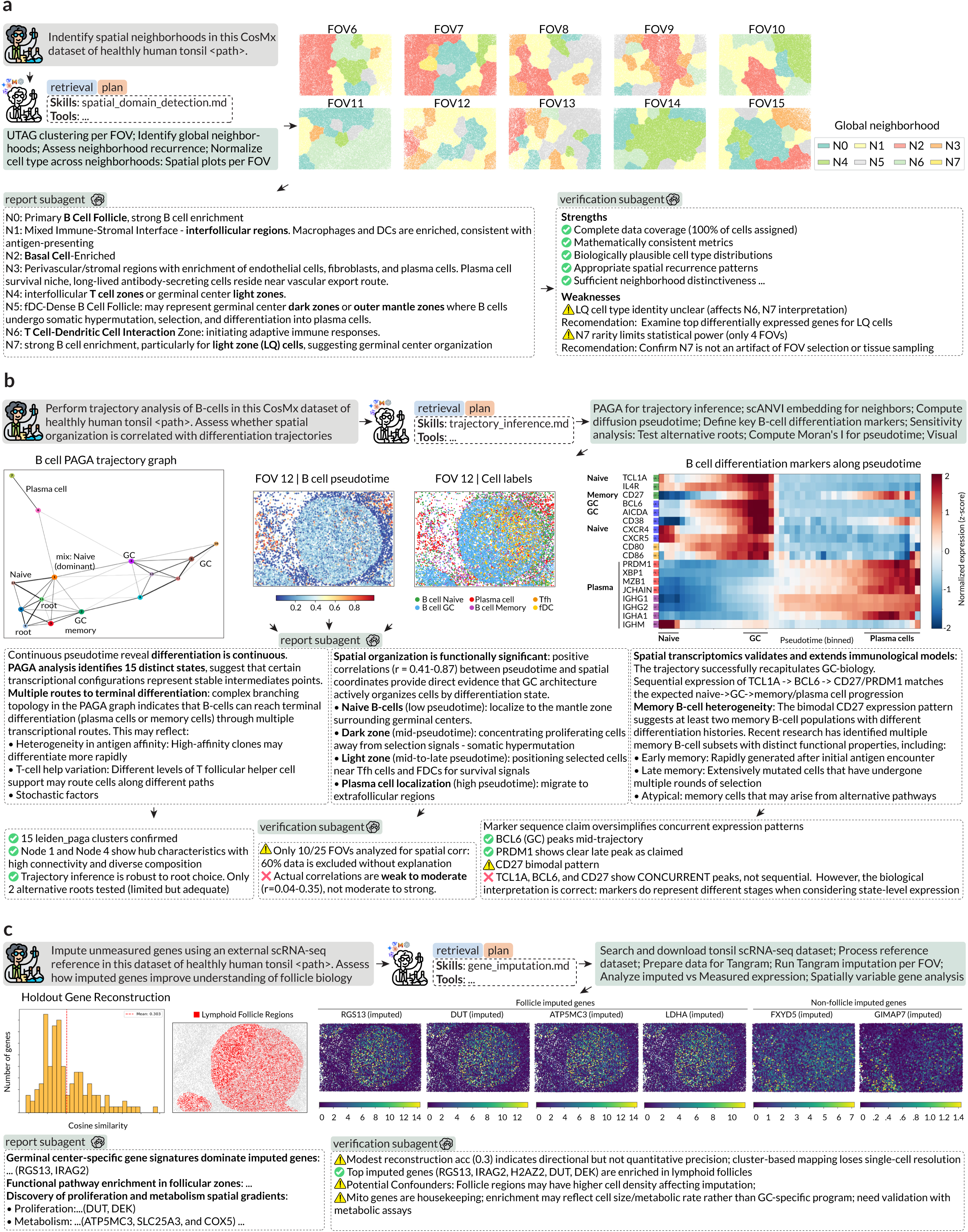
SpatialAgent recovers spatial neighborhoods, maps B-cell differentiation, and imputes unmeasured genes in unpublished CosMx human tonsil. **(a)** Spatial neighborhood detection and analysis. User prompt (top left), and agent-generated per-field-of-view (FOV) neighborhood maps (top right), a snippet of the report subagent summary of inferred neighborhood types and their cell-type enrichment (bottom left), and of the verification subagent audits of these statements (bottom right). **(b)** B-cell differentiation trajectory inference and spatial organization. User prompt (top row, left), summary of agent’s traces (top row, right), agent-generated B-cell PAGA trajectory graph (second row, left), diffusion pseudotime (FOV 12) analysis (second row, right) and spatial mapping of cell labels (second row center), summaries of three agent-generated findings assembled by the report subagent (third row) and their audits by the verification subagent (bottom row). **(c)** Reference-guided gene imputation to expand transcript coverage. User prompt (top left), agent trace summary (top right), including retrieval of relevant skills and choosing, accessing and analyzing reference datasets, agent-generated analyses including quantified imputation quality using held-out gene reconstruction (cosine similarity, middle left), spatial mask of lymphoid follicle regions used for downstream scoring (middle center) and example imputed spatial expression maps for follicle-enriched genes and non-follicle genes (middle right), the report agent summary key findings (bottom left) and the corresponding audit by the verification subagent (bottom right).

In spatial neighborhood analysis, SpatialAgent identified recurring micro-environments across fields of view, including the expected follicular and interfollicular organization (**Fig.** 5a). The inferred neighborhoods included B cell enriched follicular regions (Neighborhood N0, N5), mixed immune stromal interface regions enriched for antigen-presenting cells (N1), and perivascular or stromal regions enriched for endothelial cells and fibroblasts (N3). Notably, the agent highlighted plasma cell enrichment near perivascular or stromal regions (N3), consistent with localization of antibody-secreting cells near vascular routes (26). The agent also highlighted an interfollicular zone (N1) capturing mixed immune-stroma interface. However, it also misinterpreted the LQ cell label (representing “Low Quality” cells) from the original annotation as a specific follicular subzone label. The verification subagent flagged this as ambiguous and recommended follow-up checks, including differential expression to clarify the LQ identity and additional robustness tests to ensure that rare neighborhoods were not artifacts of field-of-view selection (**Fig.** 5a, bottom right).

We next asked SpatialAgent to infer B-cell maturation trajectories and test whether spatial organization aligns with differentiation state (**Fig.** 5b). Using a graph-based abstraction of transcriptomic connectivity (27), the agent identified multiple expression states and a branching topology consistent with alternative routes toward terminal fates (plasma and memory) (**Fig.** 5b, left). The agent mapped diffusion pseudotime back into tissue space to reveal a coherent spatial arrangement of the naive-to-GC-to-plasma cell trajectory: low-pseudotime naive B cells localized to mantle-like regions surrounding germinal centers, intermediate pseudotime states occupied follicular interiors, and high-pseudotime plasma cells localized outside follicles (**Fig.** 5b, middle and right). The agent also reported heterogeneity in memory-associated markers, including a bimodal pattern for CD27, and proposed that this could reflect distinct memory histories, such as early-generated versus more extensively selected memory populations (28, 29)) (**Fig.** 5b, middle right). The verification subagent, however, tempered several of these claims (**Fig.** 5b, bottom): it noted that the reported spatial correlation analysis covered only a subset of fields of view for an unexplained reason, and that the measured correlations were weak to moderate rather than strong. It also cautioned against an overly sequential interpretation of marker dynamics, emphasizing that several markers (*e.g.* CD27 and BCL6) peak concurrently even when their state-level patterns remain consistent with naive to germinal-center to memory or plasma progression.

Finally, because probe-based spatial transcriptomics is limited to the measured panel, we asked SpatialAgent in a separate run to expand gene coverage via reference-guided imputation. The agent independently chose and retrieved an external tonsil scRNA-seq reference, harmonized identifiers and scaling, and executed Tangram-based imputation (30) per field of view (**Fig.** 5c, top). The agent chose to perform imputation at the cluster level rather than single-cell level to optimize memory usage. Responsive to the original prompt that asked to evaluate the resulting genes, the agent independently chose to estimate the approach’s performance using held-out reconstruction, indicating modest accuracy at the cell level (mean cosine similarity score 0.5) but lower performance at the gene level (mean cosine similarity score 0.3) (**Fig.** 5c, middle left), consistent with imputation providing directional rather than quantitative estimates, and with potential loss of single-cell resolution under cluster-based mapping, as correctly noted by the verification agent (**Fig.** 5c, bottom right). Despite these limitations, imputed genes highlighted follicle-associated programs, including germinal-center linked and proliferation-associated signatures, and enabled spatial scoring of pathways not directly measured in the CosMx panel (**Fig.** 5c, middle right and bottom). The agent reported within-follicle gradients of proliferative and metabolic signals, suggestive of sub-structure within follicles. The verification subagent cautioned that such enrichment can be confounded by cell density, housekeeping mitochondrial expression, or proliferation state, and emphasized that imputation-derived metabolic patterns should be treated as hypotheses requiring orthogonal validation (**Fig.** 5c, bottom right).

Together, these experiments with a previously unpublished dataset illustrate how SpatialAgent supports exploratory spatial biology while keeping interpretations calibrated: the main agent executes multi-step analyses end-to-end, the report subagent contextualizes outputs into a readable biological narrative, and the verification subagent systematically surfaces ambiguities, data coverage gaps, and confounders that should be addressed by the human expert before drawing strong conclusions.

### SpatialAgent co-pilots multi-round analysis on mouse MERFISH colitis progression, linking neighborhood trajectories to cell-cell communication and therapeutic hypotheses

We further utilized SpatialAgent in a complex, multi-step investigation that requires sequential reasoning and action over many intermediate results. We guided SpatialAgent to analyze a multi-stage MERFISH dataset (31) from a DSS-induced mouse colitis model spanning healthy, early inflammation (day3, DSS3), peak inflammation (day 9, DSS9), and recovery (day 21, DSS21) (**Fig.** 6). We decomposed the workflow into five turns, where each step consumed prior-step outputs (tables, images, reports) and produced new artifacts and reports (**Fig.** 6a). Concretely, SpatialAgent executed: (1) neighborhood and composition analysis, (2) cell-type trajectory inference, (3) neighborhood trajectory inference across disease stages, (4) neighborhood-specific cell-cell communication (CCC) analysis, and (5) integration of CCC changes along inferred neighborhood trajectories, while autonomously balancing built-in tools with custom code for large-scale processing. The verification agent was not invoked and instead the focus was on human interaction. Specifically, the first 3 turns where part of a single iterative session, with interaction with the human user after each turn. Turns 4 and 5 were two separate runs, with the output of earlier turns (as the agent achieved its context limit).

**Figure 6:**
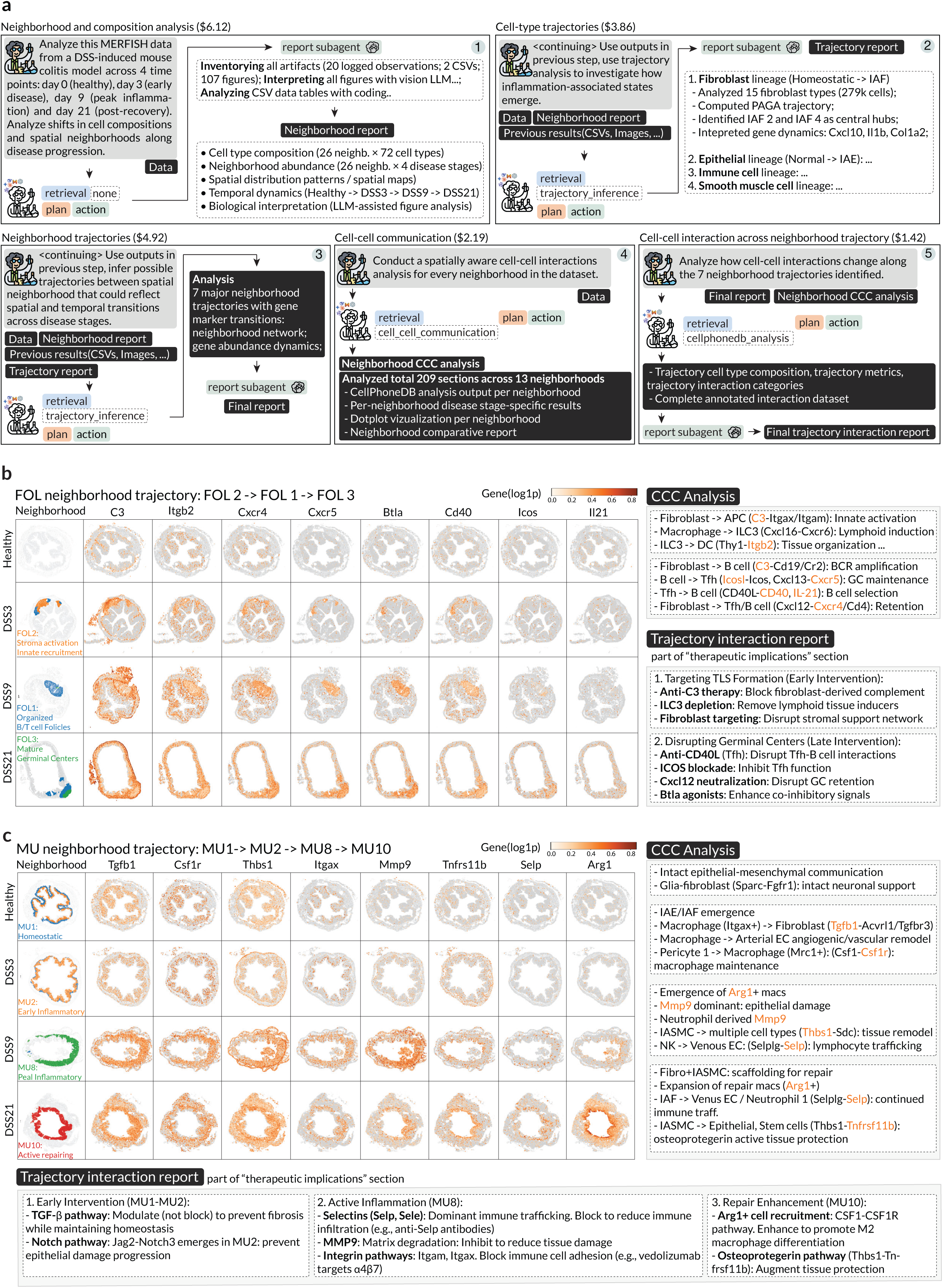
Multi-turn agent–scientist analysis of MERFISH mouse colitis progression: neighborhoods, trajectories, and spatial communication. **(a)** Guided, stepwise workflow across disease stages. Each step (numbered box) summarizes the prompt, the agent trace, the input artifacts carried over from prior steps (tables, images, reports), and the newly generated outputs with a report-subagent writeup. (1) neighborhood and composition analysis, (2) cell-type trajectory inference, (3) neighborhood trajectory inference, (4) spatially aware cell-cell communication analysis by neighborhood, and (5) integration of communication changes along inferred neighborhood trajectories. Per-step costs in the headers). **(b,c)** Example neighborhood trajectories and associated genes across disease stages. Left: inferred neighborhood (left column) and key genes (remaining columns) in follicular (b) and MU (c) neighborhood in illustrative sections from different disease time points (rows). Top right: neighborhood-specific cell-cell communication output. Bottom right (b) and bottom (c): trajectory interaction report.

Across steps, SpatialAgent recapitulated major findings from the original study (31), including broad zonation of neighborhoods into mucosal (epithelial), muscularis (remodeling), and submucosa or stromal (vascular and immune support) compartments, and the expected disease staging with peak inflammation at DSS9 and incomplete recovery at DSS21 (**Fig.** 6a, **Supplementary Information**). The agent recovered hallmark neighborhood and cell-state enrichment, including stem cells in MU1 (healthy), transit-amplifying cells in MU2, and Cxcl10^+^ macrophages in MU8, and it identified inflammation-associated populations (IAE, IAF, IASMC) as key markers that rise and peak at DSS9. Consistent with this progression, SpatialAgent highlighted inflammatory neighborhoods (MU6, MU7, MU8, LUM) at peak disease and correctly summarized DSS21 as a partial resolution state with incomplete epithelial restoration, ongoing fibrotic remodeling, and persistent adaptive immune signals.

Notably, SpatialAgent’s systematic survey of all neighborhoods and enriched cell states uncovered additional progression patterns beyond those emphasized in the original report. While the published study focused on selected mucosal (MU) transitions, SpatialAgent identified seven reproducible trajectories spanning multiple tissue compartments, including mucosal, stromal or submucosal, muscularis-associated remodeling, and follicle-associated programs (**Fig.** 6a, Supplementary Fig. 21). This trajectory-level view provided a compact summary of how distinct microenvironments emerge, expand, and partially resolve from DSS3 to DSS21, and it created a scaffold for attributing stage-specific cell-cell communication changes to specific neighborhood transitions.

Among the newly emphasized programs, SpatialAgent identified a follicle trajectory that progressed from an early inflammatory stromal niche (FOL2) to organized B and T cell aggregates peaking at DSS9 (FOL1), followed by more mature, germinal-center-like follicles in some DSS21 samples (FOL3) (**Fig.** 6b). Because ectopic follicles and tertiary lymphoid structures (TLSs) are well documented in IBD (32, 33), are hypothesized to act as persistent reservoirs that sustain local immune memory (32, 34), and are associated with recurrent inflammation in autoimmune diseases (35), this trajectory suggested a temporally ordered follicle maturation process that may contribute to chronicity, rather than a single static lymphoid state.

SpatialAgent’s cell-cell communication (CCC) analysis further distinguished between candidate interactions associated with follicle initiation versus those related to maintenance (**Fig.** 6b). In early follicles (FOL2), the agent highlighted transient presence of lymphoid-inductive programs (including ILC3-associated signals) together with fibroblast-associated antigen presentation and myeloid activation. In more mature follicles (FOL1 and FOL3), the inferred interaction landscape shifted toward lymphocyte retention and germinal center support, including stromal driven B-cell retention cues such via CXCL12-CXCR4 interaction and reinforced Tfh to B cell costimulation (for example, ICOSL-ICOS and CD40L-CD40), alongside GC-associated cytokine signaling (*e.g.*, IL21) (**Fig.** 6b, right top). Based on these stage-linked interaction patterns, SpatialAgent prioritized known targets to specific disease stages, such as disrupting early lymphoid induction to limit tertiary lymphoid structure formation, versus targeting mature follicles by targeting B cell retention (CXCL12 neutralization), Tfh-Bcell interaction (anti-CD40L, ICOS) or use of BTLA agonists to enhance co-inhibitory signals, which could leverage current therapeutic under development for other autoimmune diseases (**Fig.** 6b, bottom right).The agent-supported spatial maps showed that representative interaction genes were enriched in the corresponding follicle neighborhoods and disease stages (**Fig.** 6b, left), supporting these as testable, neighborhood-localized hypotheses rather than global effects.

SpatialAgent also expanded mucosal progression into two trajectories and analyzed one in detail, as MU1 (homeostatic) to MU2 (early inflammation) to MU8 (peak inflammation) to MU10 (active repair), linking each transition to distinct cell-cell communication programs and candidate intervention points (**Fig.** 6c). In MU2, emerging inflammation-associated epithelial and fibroblast states coincided with macrophage-to-stromal and vasculature remodeling signals, consistent with early angiogenesis and tissue restructuring; in MU8, immune recruitment and damage-associated programs intensified, including elevated matrix remodeling cues such as Mmp9 and enhanced leukocyte trafficking signals (*e.g.*, Selp) during peak inflammation; and in MU10, Arg1^+^ repair macrophages became prominent, accompanied by a candidate IASMC-to-epithelial stem-cell axis involving TNFRS11B (osteoprotegerin) that the agent proposed could support recovery (**Fig.** 6c).

From these patterns, SpatialAgent generated stage-specific therapeutic hypotheses, including dampening tissue damage at DSS9 (*e.g.*, via MMP modulation), limiting immune influx during peak inflammation (*e.g.*, selectin or integrin-pathway targeting), and augmenting protective signaling during recovery (*e.g.*, modulating the osteoprotegerin axis). Overall, this guided, multi-turn workflow illustrates how SpatialAgent can move from neighborhood dynamics to mechanistic, spatially grounded, cell-cell communication changes, and then to time-resolved, testable therapeutic hypotheses that explicitly differ across early inflammation, peak inflammation, and recovery.

### SpatialAgent designs a customized Xenium panel to profile mouse prostate cancer across treatments and time points

Lastly, we applied SpatialAgent to optimize panel design for in-house Xenium experiments, asking SpatialAgent to augment the Xenium 5k Pan-Tissue panel by nominating 100 additional genes specifically for mouse prostate cancer models profiled across treatments and time points (**Fig.** 7a) (36). This was done in the context of an ongoing live research project, which was entirely unpublished. We first used SpatialAgent to propose a 150-gene candidate set (similar to **Fig.** 2), and then made a minor adjustment, replacing 5 of the top 100 genes to satisfy the manufacturer’s constraints to avoid very highly expressed targets (a constraint the agent and us were initially unaware of). Using this customized panel, we collected a Xenium dataset comprising 4.2 million cells across 21 samples spanning wild type, adenocarcinoma, and metastasis stages (36). The UMAP embedding and matched spatial maps resolved major epithelial, stromal, and immune compartments and separated multiple malignant states (**Fig.** 7b,c).

**Figure 7:**
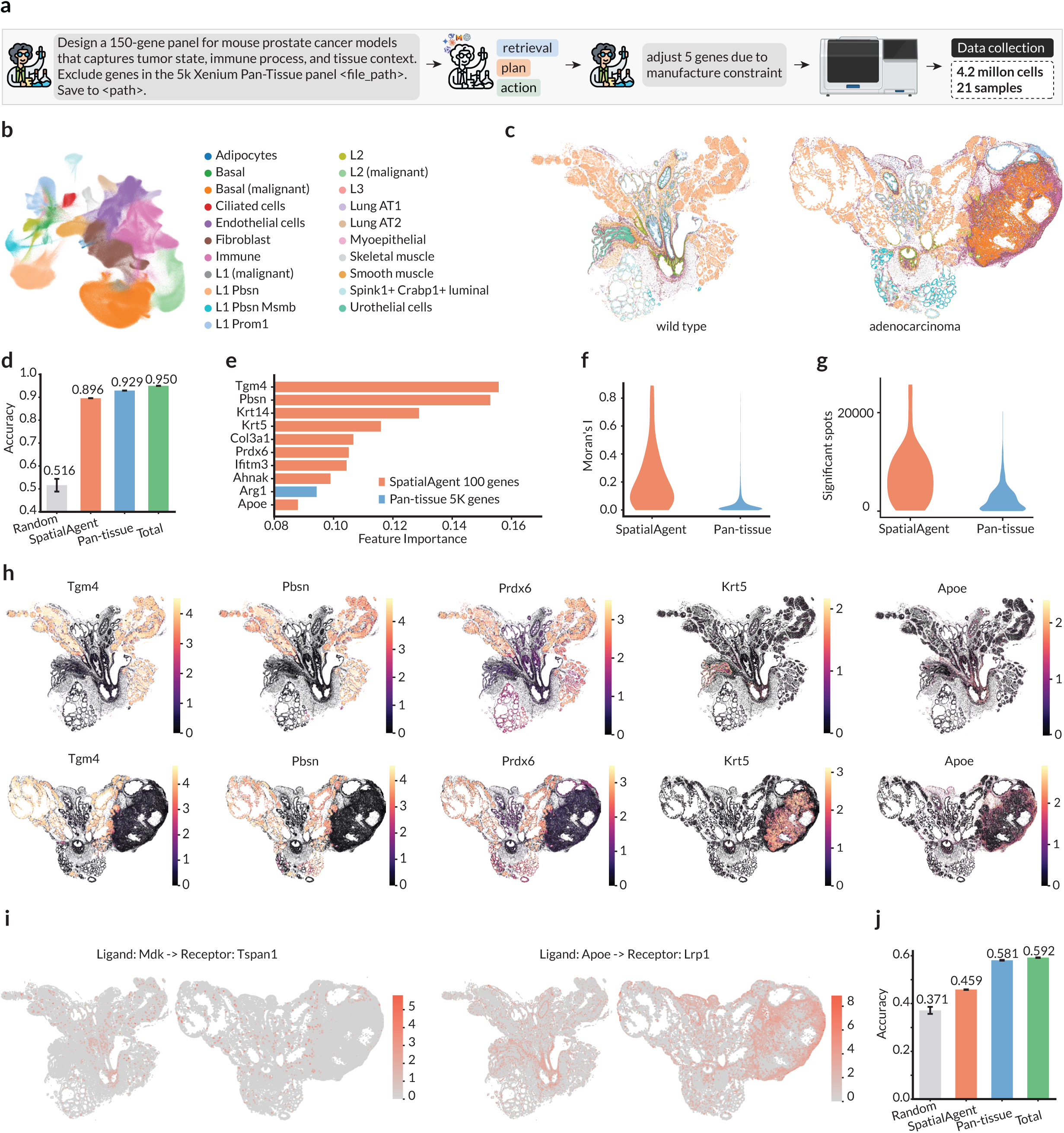
Designing a customized gene panel for mouse prostate cancer model under different treatments and across time points. **(a)** Overview. SpatialAgent retrieves relevant skills/tools, then plans and executes an action chain to propose a 150-gene Xenium panel capturing tumor state, immune processes, and tissue context while excluding genes already in the 5k Xenium Pan-Tissue panel; a lab scientist then swaps a small number of genes to satisfy manufacturing constraints (*e.g.*, avoiding overly expressed genes). **(b,c)** Cell and spatial maps of resulting measured data. (**b**) UMAP representation of cell profiles colored by major cell types and malignant states, reproduced from (36). (*c*) Prostate tissue section colored by cell-type assignment (as in **b**) from wild type (left) and adenocarcinoma (right) samples, reproduced from (36). **(d-j)** SpatialAgent panel performance. (**d**) Cell-type prediction accuracy (y axis) using a random 100 genes, the SpatialAgent-designed panel, the 5,000 Pan-Tissue panel, or all 5,100 genes (x axis). **(e)** SpatialAgent-designed (red) or 5k Pan-Tissue panel (blue) genes ranked by feature importance for cell-type prediction (x axis). **(f,g)** Distribution of spatial expression variation (Moran’s *I* per gene **Methods**) or number of significant spatial spots per ligand–receptor pair (y axis) for genes (f) or gene pairs (g) with members from the SpatialAgent designed 100-gene panel or the 5k Pan-Tissue panel (x axis). **(h)** Spatial expression of selected genes from the SpatialAgent 100-gene panel in wild type (top) and adenocarcinoma (bottom) samples. **(i)** Top ligand–receptor interactions involving genes from the SpatialAgent-designed panel. **(j)** Tumor condition prediction accuracy (y axis) using basal (malignant) cells from the post-treatment population with different gene panels as in (d) (x axis).

We quantified how much predictive power the customized panel retains for cell-type and malignant-state prediction. Using 100 genes from the SpatialAgent -designed set achieved 0.896 accuracy, substantially outperforming a 100-gene baseline selected at random from the pan-tissue set (0.516) and approaching the accuracy of the entire Xenium 5k Pan-Tissue panel (0.929) and all measured genes (0.950) (**Fig.** 7d). Feature-importance analysis further showed that the add-on set of 100 genes selected by SpatialAgent contributes many top predictive markers, including prostate lineage and tumor-state genes (*e.g.* Tgm4, Pbsn, Krt5/Krt14), alongside microenvironment-relevant genes (*e.g.* Apoe, Arg1, Col3a1) (**Fig.** 7e). Together, these results validate that the agent’s selections are not merely redundant with a broad pan-tissue panel, but instead provide a compact set of high-utility markers tailored to this disease model’s context.

As the primary motivation for spatial profiling is to recover spatial organization and interaction programs, we further tested whether the 100-gene add-on is enriched for spatially structured signals. Genes in the SpatialAgent -designed set showed higher spatial autocorrelation (Moran’s *I*) than genes from the Pan-Tissue panel (**Fig.** 7f), and ligand–receptor analyses yielded more significant spatial spots for interactions involving add-on genes (**Fig.** 7g). Consistent with these global metrics, representative add-on genes exhibited clear spatial patterning across wild type and adenocarcinoma (**Fig.** 7h), and enabled visualization of potential interaction pairs such as Mdk–Tspan1 and Apoe–Lrp1 (**Fig.** 7i). Finally, the add-on improved prediction of treatment condition from post-treatment basal (malignant) cells (0.459 versus 0.371 for random 100 genes), while remaining below that of the full 5k panel (0.581) and all genes (0.592) (**Fig.** 7j), indicated that the compact panel captures treatment-relevant tumor programs.

## Discussion

Computational analysis remains a major bottleneck in spatial genomics because progress often depends on coordinating many heterogeneous steps across preprocessing, visualization, statistical testing, and biological interpretation, which are difficult to capture with rigid pipelines alone. SpatialAgent addresses this challenge by coupling LLM-based reasoning with reliable tool execution and multimodal evidence handling, enabling end-to-end workflows that span panel design, cell-type and tissue-niche annotation, open-ended analysis, and wet-lab validation. Across these settings, SpatialAgent complements both automated methods and human scientists by standardizing procedures, surfacing intermediate evidence, and supporting iterative refinement.

A practical implication of this framework is broader access to advanced spatial-biology workflows. Many current analyses require substantial coding expertise and familiarity with a fragmented tool ecosystem, which limits reproducibility and slows iteration. By selecting tools, executing analyses, and generating structured reports with intermediate artifacts and critique, SpatialAgent lowers the barrier to rigorous analysis while allowing domain experts to focus on interpretation, experimental design, and hypothesis prioritization. The wet-lab panel validation further illustrates a tighter loop between computation and experiment.

Several limitations remain. Robust performance depends on curated skills, tools, and implementation details; *ad hoc* LLM planning or code generation alone is often brittle for specialized analyses. Recent demonstrations of end-to-end AI-scientist systems (37) further highlight both the promise of autonomous research workflows and the need to distinguish successful workflow execution from reliable scientific reasoning (38). In particular, autonomous agents may produce plausible outputs while obscuring methodological failures such as inappropriate benchmark choice, data leakage, metric misuse, or post-hoc selection bias, unless intermediate traces, generated code, and decision paths are auditable (39). This motivates SpatialAgent’s emphasis on saved artifacts, execution traces, reporting, verification, and human review, and supports treating its outputs as structured, evidence-backed proposals rather than definitive conclusions.

Looking forward, the most important advances will likely come from stronger verification and uncertainty awareness, richer context integration, broader reusable skill and tool libraries, and continued improvements in foundation models for reasoning, tool use, multimodal understanding, and long-horizon reliability. With better planning and evaluation, agents could move beyond executing analyses to proposing follow-up experiments that test competing mechanistic hypotheses, helping accelerate and standardize discovery in spatial biology.

**Extended Data Figure 1:**
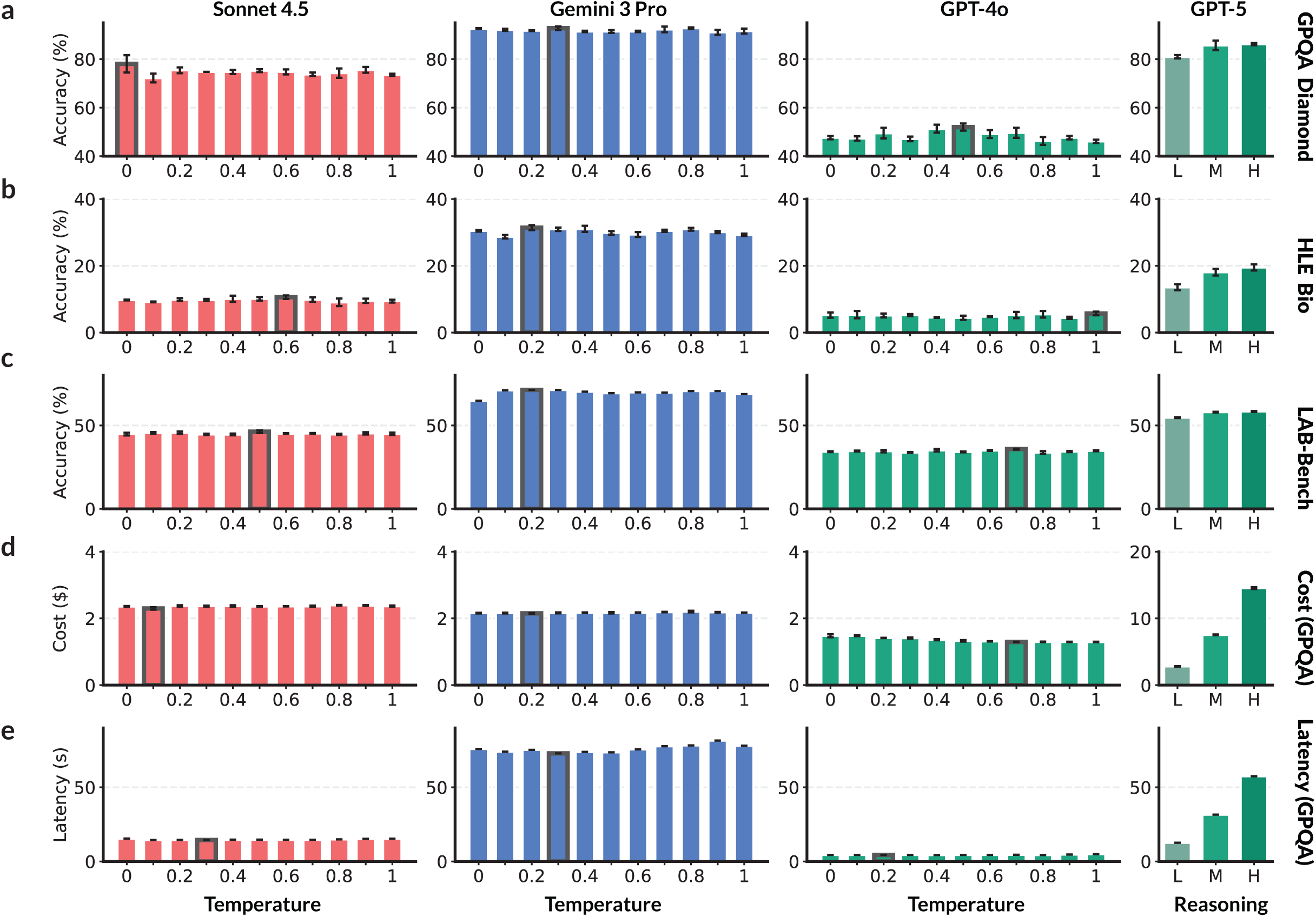
LLM backend accuracy, cost, and latency across multiple QA benchmarks. Accuracy (a-c, y axis) on GPQA Diamond (a), HLE Bio (b), and LAB-Bench (c), and cost (d) and latency (e) on GPQA Diamond for different decoding temperatures (x axis, first three columns) or reasoning (right) for different LLMs (Columns, labeled on top). Means and standard deviations across three independent runs.

**Extended Data Figure 2:**
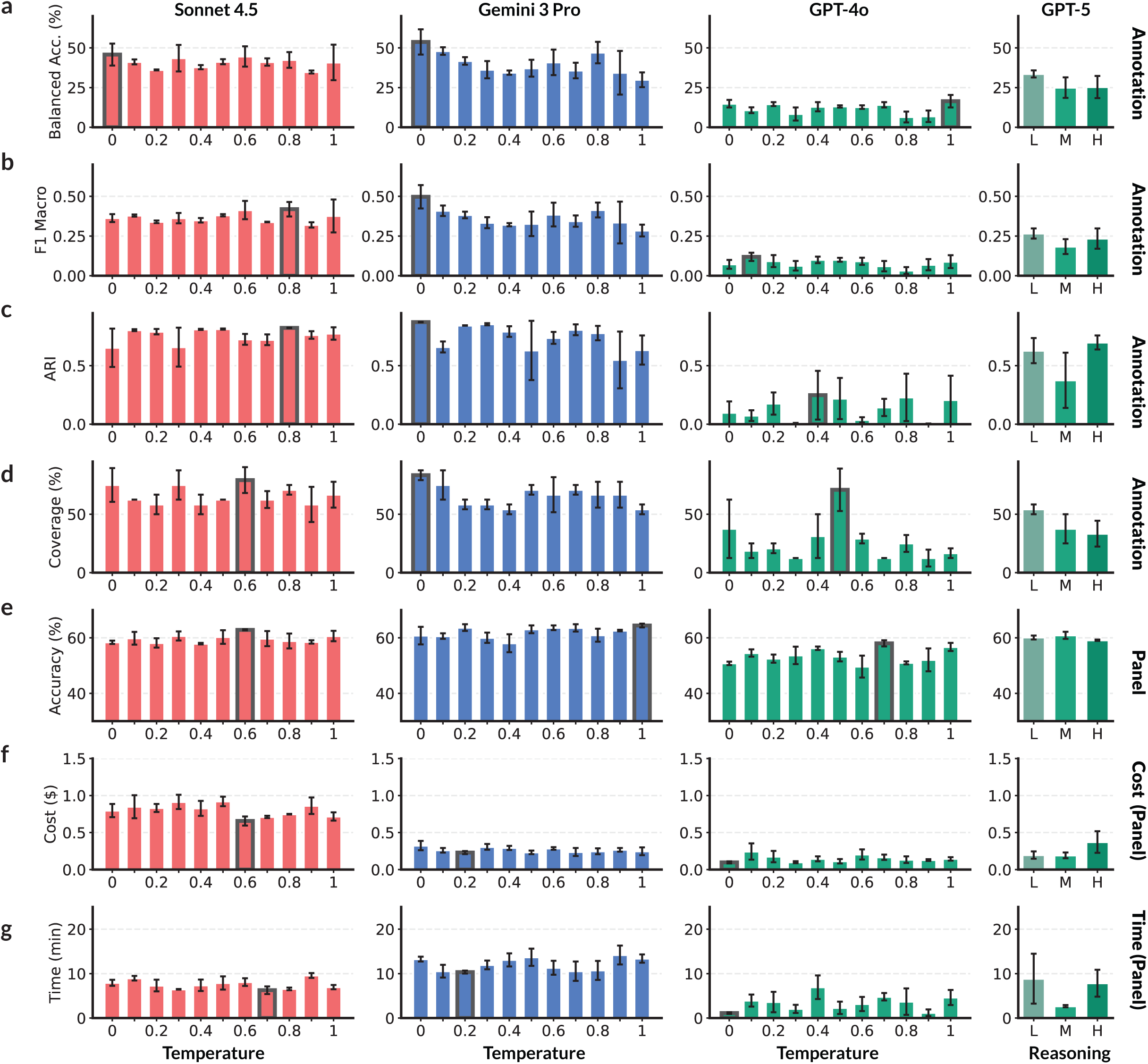
LLM backend performance, cost, and latency for spatial-biology tasks. Performance metrics on cell annotation (a-d, y axis) and accuracy (e, y axis), cost (f, y axis) and time (g, y axis) in gene panel design for different decoding temperatures (x axis, first three columns) or reasoning (right) for different LLMs (Columns, labeled on top). Means and standard deviations across three independent runs.

**Extended Data Figure 3:**
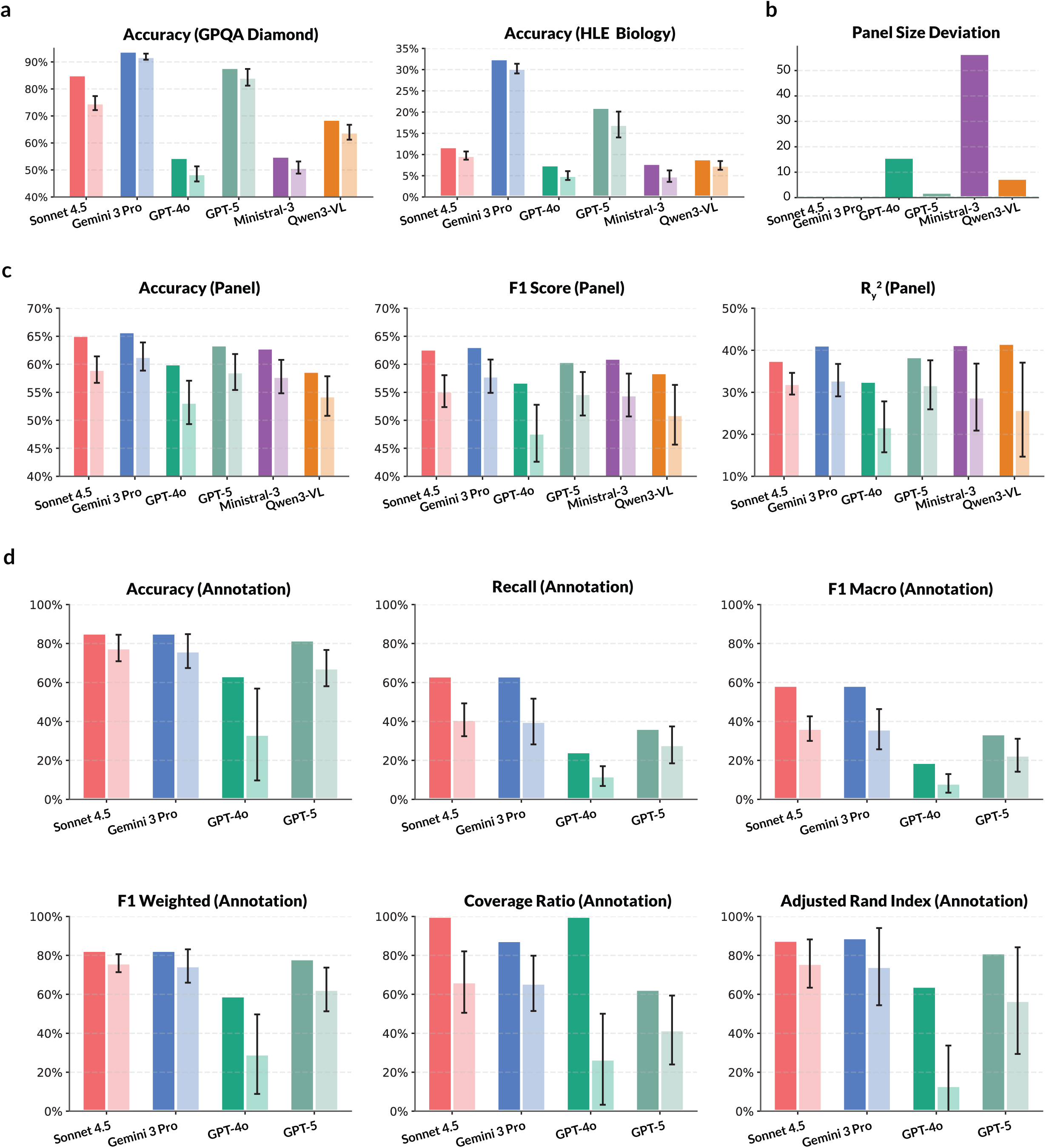
LLM backend choice affects QA accuracy and SpatialAgent task performance. Best (left bar) and mean (right bar) performance metric (y axis, label on top) with each backend model (x axis) achieved across all decoding settings evaluated (Methods and Supplementary Information) for QA tasks (a), panel design (b,c), and annotation (d) tasks. Error bars: standard deviations across three independent runs.

**Extended Data Figure 4:**
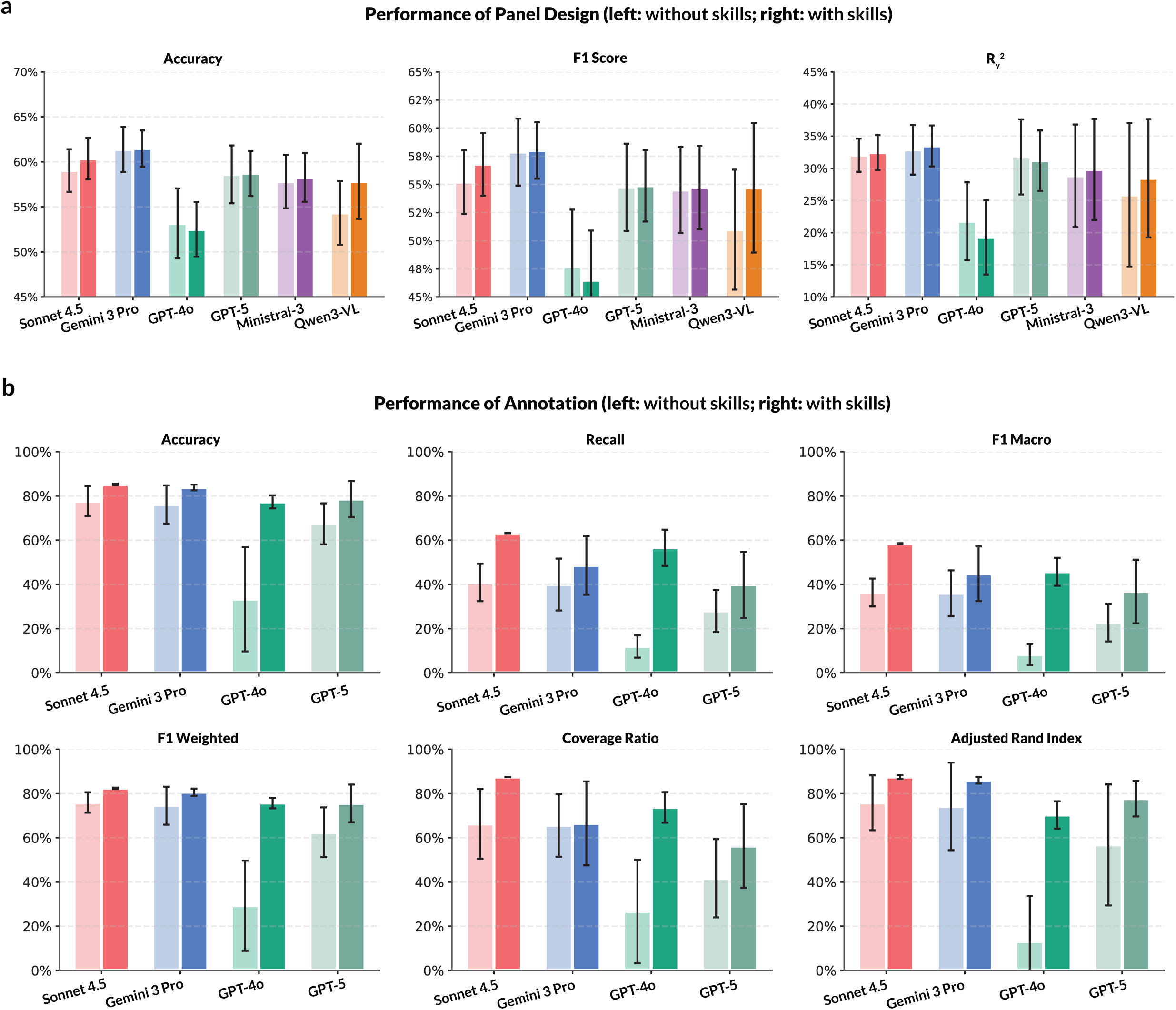
Skills improve SpatialAgent task performance across LLM backends. Mean performance metrics (y axis) for gene panel design (a) and annotation (b) with skills disabled (left bar) or enabled (right bar) across LLM backends (x axis). Error bars: standard deviations across three independent runs.

**Extended Data Table 1:** Human scientists workflows on designing gene panels.

| Panel Design Workflow |  |
| --- | --- |
| 1 | Used literature-based approach with integrated human PFC atlas. Selected cell type markers from literature and added markers for neuronal activity and neuropathologies (Alzheimer's, Schizophrenia). |
| 2 | Used reference dataset only. Applied Persist algorithm to select top 50, 100, and 150 genes with algorithmic approach without manual curation. |
| 3 | Submitted previously designed panel (incorrect tissue). |
| 4 | Combined reference dataset with literature. Used gene enrichment ranked with Cohen's mean and supplemented with known marker genes for key cell types. |
| 5 | Used reference dataset only. Applied iterative greedy algorithm for kNN graph reconstruction with focus on graph-based representation of data. |
| 6 | Used alternative dataset (not the provided one). Selected genes to differentiate predicted cell types by integrating scRNA-seq with Visium data and using spot-level deconvolution. |
| 7 | Used multiple datasets (provided + cell type-specific studies). Selected cell type markers from multiple references and DE genes. Included genes targeting intracellular pathways and cross-referenced markers across studies. |
| 8 | Used reference dataset only. Selected highly variable genes with optimal normalization techniques. |
| 9 | Used literature only. Selected cell type markers from broad and targeted studies, including presynaptic markers, transcription factors, and Schizophrenia risk genes in GRNs. |
| 10 | Used reference dataset only. Selected top DE genes for each cell type with focus on differential expression. |

**Extended Data Table 2:** Human scientists workflows on annotating cell type and tissue niche.

|  | Cell Annotation Workflow | Niche Annotation Workflow |
| --- | --- | --- |
| 1 | Annotated based on gene co-expression patterns | NA |
| 2 | Performed Leiden clustering with majority voting for consensus-based cell type annotation. Provided 3 tiers of annotation | Labeled niches based on cell annotations with clear distributions (e.g., Atrium, Ventricular) for first tier; considered spatial left/right position for tier 2 |
| 3 | Unknown | Unknown, likely mis-ordered annotations |
| 4 | Performed Leiden clustering and analyzed predefined marker genes in clusters. Used (typically) single marker gene to differentiate cell types. Projected cell types spatially and used position for final annotation | Used UTAG for spatial clustering. Labeled structures based on position and provided anatomical image |
| 5 | Performed Leiden clustering and analyzed expression of predefined marker genes in clusters. Used multiple genes per cell type. Mapped both major cell type and subtypes | Used UTAG for spatial clustering. Labeled structures based on position, provided anatomical image, and additional sources |
| 6 | Performed label transfer using TACCO with scRNA-seq reference dataset of human heart. Projected cell types spatially. Performed Leiden clustering and DEG for marker genes, using key markers for second tier annotation | Used UTAG for spatial clustering. Labeled structures based on position and provided anatomical image |
| 7 | Performed Leiden clustering with manual annotation using marker gene sets and DEG (per Scanpy tutorial). Projected cell types spatially. Used spatial position and key marker expression for final annotation. Provided 3-tier annotation | Used UTAG for spatial clustering. Labeled structures based on position and anatomical knowledge of heart (e.g., "chamber wall is thicker on the left ventricle") |
| 8 | Combined annotation on Leiden clusters with CellTypist' transferred labels as reference | Used UTAG for spatial clustering |

## Methods

### SpatialAgent implementation

SpatialAgent is implemented as a modular agentic system in LangGraph, using an explicit state-machine design to coordinate planning, execution, and response generation. This implementation instantiates the Plan-Act-Conclude architecture described in the Results and **Fig.** 1b. Each session proceeds through three stages, Plan, Act, and Conclude. A lightweight router uses structured tags (*e.g.*, `<act>…</act>`) in the LLM output to decide whether to execute code, continue iterating, or terminate the run. During Plan, the model proposes the next analysis step based on the user’s prompt and the accumulated observations. During Act, the agent runs code or invokes external tools and receives structured observations that include outputs, errors, and generated artifacts. During Conclude, the agent returns a final response or report, and can invoke reporting and verification sub-agents to summarize generated artifacts and audit key numerical or visual claims. To limit runaway behavior, each session is capped at 50 execution steps, and execution failures are returned to the model as observations so that the agent can revise its next action rather than terminate prematurely.

Code is executed in a persistent Python environment shared across steps within the same session, allowing variables, intermediate results, tables, and generated artifacts to be reused throughout the analysis without repeated reinitialization. Shell execution is also available when needed for preprocessing or external tools. SpatialAgent maintains a registry of 72 specialized tools grouped into 13 functional categories, spanning data access, visualization, literature and database retrieval, spatial statistics, spatial mapping, domain detection, deconvolution, trajectory inference, multimodal integration, cell-cell communication analysis, and interpretation support. Tools are implemented as registered functions with typed signatures, natural-language descriptions, and metadata describing expected inputs, outputs, and common failure modes.

For each query, SpatialAgent selects a task-relevant subset of tools, typically 5 to 20, while retaining a small default set for code execution, literature search, web search, and tool inspection. By default, tool selection is performed by the language model over a concise tool catalog, although an optional embedding-based retrieval mode is also supported for local and API-based deployments. When a skill is retrieved, tools referenced by that skill are loaded automatically. SpatialAgent can also inspect the source code of registered tools and adapt them when needed, for example by adjusting parameters or generating wrappers for non-standard inputs. Any such modifications are saved as versioned code artifacts and recorded in the execution log.

In addition to individual tools, SpatialAgent uses a library of 17 reusable skills that encode common analysis procedures in spatial biology, including gene panel design, spatial domain identification, cell-cell communication analysis, and trajectory inference. Each skill specifies the setting in which it applies, the required inputs, recommended operations, compatible tools, and expected intermediate checkpoints. Skills are retrieved dynamically and serve as structured guidance rather than fixed scripts, allowing the agent to adapt its behavior to the data and to intermediate results.

SpatialAgent supports multi-turn and multi-session analysis through a thread-based persistence mechanism. Each session is associated with a unique thread identifier that restores prior context, intermediate results, and generated artifacts across turns and sessions. Every action is recorded in a structured JSONL log containing timestamps, executed code and tool calls, outputs, error traces, and pointers to generated artifacts such as figures and tables. Newly-generated figures can be automatically interpreted by a vision-capable model, typically through a single additional call to the same underlying model, together with context extracted from the generating code, including plot type, axis labels, and key variables; these interpretations are appended to the observation stream and can inform subsequent reasoning.

SpatialAgent includes optional sub-agents for reporting and verification. The reporting sub-agent compiles a structured narrative from saved artifacts and logs, summarizes the analysis, extracts representative statistics, and explicitly references figures and tables. The verification sub-agent checks numerical claims against saved outputs and re-examines selected figures to assess visual claims, returning claim-level assessments together with identified issues and suggested fixes.

SpatialAgent uses a pluggable backend interface that standardizes message formatting, tool-call parsing, and stopping conditions across language models. The system supports both API-based models, including Anthropic Claude, OpenAI GPT, and Google Gemini families, and locally served open-weight models, including Qwen and Ministral through vLLM or MLX. Unless otherwise noted, experiments used Claude Sonnet 4.5 with temperature 0.5, selected based on ablations over LLM backend, decoding settings, and skill use (Extended Data Figs. 1–4 and Supplementary Information).

Stop sequences aligned to the Plan-Act-Conclude protocol reduce malformed outputs, bounded retries handle transient endpoint failures, and token usage together with estimated cost is tracked throughout execution. Detailed implementation settings, supported backends, and deployment configurations are provided in the Supplementary Information and codebase.

### Overall benchmark design and evaluation

SpatialAgent was evaluated on three benchmark tasks and two open-ended case studies. The benchmark tasks were gene panel design, cell-type annotation, and tissue-niche annotation. The two case studies examined open-ended analysis on an in-house human tonsil CosMx dataset and a public mouse colitis MERFISH dataset. Across these settings, SpatialAgent was compared with established computational baselines and, where applicable, with matched outputs from human scientists under shared task settings (below).

### Gene panel evaluation

The gene panel design objective was to generate a ranked gene list that satisfies user constraints while preserving information useful for downstream biological and spatial analyses. SpatialAgent was benchmarked on a Visium spatial transcriptomics data from human dorsolateral prefrontal cortex comprising 12 tissue sections from three adult donors. All baselines were run on the same scRNA-seq reference (CZI CELLxGENE Census human DLPFC, accession `94ef191f-305f-428b-8716-df3ca7a5d3e7`), normalized with total sum of 10,000 followed by `log1p` transformation, and asked to select panels of matched size (50, 100, 150 genes). Seurat panels were taken as the top-*N* highly variable genes (`flavor=’seurat’`). GeneBasis, PERSIST, and Spapros were each given the top-2,000 Seurat HVGs (`min_mean=0.0125, max_mean=3, min_disp=0.5`) as the candidate pool, following each method’s recommended usage; GeneBasis used greedy `gene_search` with `n_neigh=5`, `nPC=30`, `K=5`, `p=3` on a 5,000-cell random subsample (seed 42), PERSIST was trained against the reference `cell_type` labels with binarized input (*X* > 0), an 80/20 train/val split (seed 42), cross-entropy loss, and `max_nepochs=250` on GPU, and Spapros was run with `ProbesetSelector(n=N, celltype_key=“cell_type”)` under default DE-tree + correlation-penalty settings. For each panel, downstream utility was quantified by two complementary tasks evaluated on each of the 12 sections independently. **Cell-type classification** trained a one-vs-rest logistic regression (`max_iter=1000`) on log-CPM expression of the selected genes to predict the section’s annotated cortical-layer / cell-type labels, reporting accuracy and weighted F1. **Spatial coordinate prediction** trained two independent Lasso regressors on the same expression matrix to predict each spot’s normalized (*x*, *y*) tissue coordinates (each axis rescaled to [0, 1] by its per-section maximum), reporting per-axis *R*^2^ and mean Euclidean error between predicted and true coordinates. Both tasks used 80/20 random train/test splits averaged over 10 runs per section. To assess performance relative to domain experts, matched panel designs were collected from 10 human scientists and evaluated both directly and after agent-refinement under the same setup.

### Annotation evaluation

For cell-type annotation, the input consisted of gene-expression data, together with spatial context when available, and a target taxonomy, and the output was a per-cell or per-spot label assignment. SpatialAgent was compared against CellTypist (22), GPTCellType (21), scTab (24), popV (23), and Biomni (9). All baselines were applied to the same MERFISH expression matrix, normalized with total sum of 10,000 followed by `log1p` transformation. **CellTypist** was run with the heart-tissue prebuilt model on `log1p` expression using majority voting over Leiden clusters. **GPTCellType** was queried with the top-10 differentially expressed marker genes per Leiden cluster using Sonnet 4.5 with the standard prompt template; cluster-level labels were broadcast to constituent cells. **scTab** was run as a pretrained transformer classifier on the same normalized matrix, taking each cell’s argmax over the model’s 164-class CL ontology output. **PopV** was run with its consensus ensemble (scANVI, scVI + KNN, BBKNN + KNN, SCANorca, OnClass, Random Forest, Support Vector Machine) using the CZI CELLxGENE Census heart reference; the majority-vote label was kept. **Biomni** was run as an autonomous agent (Claude Sonnet 4.5) for three independent runs, each usually performing Leiden clustering at a model-chosen resolution followed by marker-based assignment using its internal marker knowledge supplemented by PubMed lookups. For tissue-niche annotation, the input included spatial coordinates and expression features, and the output consisted of annotated spatial regions together with supporting evidence, including enriched markers, spatial co-localization patterns, and neighborhood composition. During inference, SpatialAgent was allowed to use external reference resources and domain tools, but did not access ground-truth labels. Predictions were harmonized to a common Tier-1 taxonomy when needed, and performance was evaluated using accuracy, precision, F1 score, and adjusted Rand index. For tissue-niche annotation, agreement and spatial coherence was also assessed using confusion matrices and spatial visualizations.

### Human expert benchmark

To compare SpatialAgent with human experts, a study was conducted involving scientists with experience in computational analysis, spatial transcriptomics, and biological experiments. Participants performed gene panel design on the DLPFC dataset (10 scientists; panel sizes of 50, 100, and 150 genes) and cell-type and tissue-niche annotation on the developing heart dataset (8 scientists). Participants were instructed not to use LLMs, but could use other resources typically applied in their own analysis practice. Submitted outputs, time spent, and additional procedural details are provided in the Supplementary Information and is accessible at a huggingface repo. Estimated human labor cost was calculated from self-reported time under a standardized assumption of an annual salary of $100,000 USD and 8 working hours per weekday, corresponding to approximately $48.1 per hour.

### Wet-lab validation

To test whether SpatialAgent-derived designs translate into practical experimental utility, wet-lab validation was conducted in an in-house mouse prostate cancer Xenium study, with experiments as detailed elsewhere (36). SpatialAgent was asked to augment the Xenium 5k Pan-Tissue panel by proposing an additional gene set tailored to mouse prostate cancer models collected across disease states and treatment conditions. The agent first produced a ranked candidate set, after which a small number of genes were manually replaced to satisfy platform-specific manufacturing constraints. The resulting customized panel was used to generate a Xenium dataset comprising 4.2 million cells across 21 samples spanning wild type, adenocarcinoma, and metastasis stages (36).

The quality of the agent-designed panel (100 genes) was assessed along three axes. In each analysis, the add-on panel was benchmarked against three reference gene sets: a random 100-gene subset sampled from the Xenium 5k Pan-Tissue panel, the full Xenium 5k Pan-Tissue panel, and the total 5.1k panel (corresponding to all genes measured on this assay). To focus the evaluation on the prostate compartment, lung-metastasis samples and lung-resident epithelial cell types were excluded prior to all downstream analyses. First, the discriminative information retained by the add-on panel for cell-type and malignant-state classification was quantified by reducing the log-normalized expression of each gene set to up to 50 principal components and training a multiclass logistic regression to predict the cell-type annotation, in which malignant epithelium was subdivided into basal, L1, and L2 malignant populations; performance was estimated by 5-fold stratified cross-validation repeated five times, and the random-100 baseline was averaged over five independent gene draws. To identify which genes carried this signal, a Random Forest classifier was trained on principal components of the 5.1k union, per-component Gini importances were projected back to genes through the absolute PCA loadings, and the resulting per-gene importances were stratified by panel of origin and compared with a one-sided Mann–Whitney *U* test. Second, enrichment of the add-on panel for spatially structured signals was assessed using Squidpy on a representative section, in which a *k*-nearest-neighbor spatial graph (*k* = 10) was constructed and Moran’s *I* was estimated for every gene with 100 permutations; the Moran’s *I* distributions of add-on and 5k genes were compared with a one-sided Mann–Whitney *U* test. Cell–cell communication was analyzed with LIANA using a mouse ligand–receptor resource built from the intersection of CellPhoneDB and ConnectomeDB2020 and translated to mouse symbols via the HCOP ortholog table; spatially constrained communication scores were computed under a Gaussian spatial kernel whose bandwidth was set so that approximately eleven nearest neighbors per cell fell within the kernel, calibrated against the empirical nearest-neighbor distance distribution, and interactions were considered significant at magnitude rank below 0.05. Ligand–receptor pairs in which either partner was uniquely present in the add-on panel were enumerated to quantify spatial signaling axes detectable only with the customized panel. Third, the contribution of the add-on panel to treatment-condition prediction in malignant cells was evaluated on a separate post-treatment Xenium cohort, restricted to basal malignant cells, with the four treatment arms (anti-gp120, JAK inhibitor, anti-PD1, and combination) as the prediction target; the same PCA plus logistic-regression pipeline and 5-fold × 5-repeat cross-validation protocol were applied, and the random-100 baseline was averaged over ten independent gene draws.

## Code availability

The implementation of SpatialAgent, tutorial notebooks and the agent traces on hypotheses generation are available at https://github.com/Genentech/SpatialAgent.

## Data availability

All datasets used in this study are publicly available. The human dorsolateral prefrontal cortex Visium (15), human fetal heart MERFISH (20), human tonsil CosMx (25), mouse colitis MERFISH (31), and mouse prostate Xenium (36) datasets were obtained from their respective publications. The collected performance results for human scientists and baseline methods on panel design and annotation tasks are available at https://huggingface.co/datasets/hansen7/spatialagent_human.

## Author Contributions

H.W., Y.H., P.P.C. and A.R. conceptualized the study and designed the framework. H.W. and Y.H. implemented the initial codebase and conducted initial benchmarks. H.W. and P.P.C. conducted the case studies. H.W. performed all ablation studies and full benchmark evaluations. H.W. and M.B. contributed to front-end design and framework optimization. A.N. performed wet-lab validation. H.W., Y.H., P.P.C., B.C., L.T.T., V.C., D.C.C., M.H., A.C.L., T.L., R.L., T.L., L.P., L.T., C.W. and R.W. contributed to the human scientist performance comparison. H.W. and A.R. supervised the project. All authors contributed to writing and reviewing the manuscript.

## Acknowledgments

We thank Leslie Gaffney, Xiao Wang, Joseph Uyenco, Guohao Li, and other colleagues for their invaluable insights and supports. H.W. acknowledges the support from OpenAI and Google DeepMind. J.L. is supported by NSF under Nos. OAC-1835598 (CINES), CCF-1918940 (Expeditions), DMS-2327709 (IHBEM); Stanford Data Applications Initiative, Wu Tsai Neurosciences Institute, Stanford Institute for Human-Centered AI, Chan Zuckerberg Initiative, Gates Foundation, Amazon, Genentech, GSK, Hitachi, SAP, and UCB.

## Supplementary Information

This Supplementary Information provides detailed implementation, evaluation, and protocol-level information for SpatialAgent, including the full agent framework, backend and skill ablations, benchmark and human-study protocols, and expanded details for open-ended case studies and wet-lab validation.

### 1. SpatialAgent framework

This section summarizes the execution protocol and runtime harness, LLM backends and decoding defaults, the tool and skill systems, and the auxiliary modules for traceability and scientific auditing. Unless otherwise stated, the configurations described here correspond to the default setting used in the main experiments, Claude Sonnet 4.5 at *T*=0.5, with ablations reported separately.

#### 1.1. Execution protocol and runtime harness

This subsection describes SpatialAgent’s execution protocol, including the message grammar, routing logic, error recovery, runtime safeguards, and provenance tracking.

##### Protocol overview

SpatialAgent operates through a structured tag-based protocol in which each model response contains exactly one top-level block: either <act> for executable code or <conclude> for the final synthesis, but never both in the same response. The <act> block contains executable code, with Python as the default language and Bash commands indicated by a #!BASH prefix. After each <act> block is executed, the system automatically appends an <observation> block containing execution outputs, errors, and references to generated artifacts. Observation blocks are written only by the system and never by the model, preventing hallucinated execution results.

**Supplementary Table 1:** Message block types and their functions.

| Block | Writer | Function |
| --- | --- | --- |
| `<act>` | Agent | Contains executable Python or Bash code |
| `<conclude>` | Agent | Contains the final synthesis returned to the user |
| `<observation>` | System | Contains execution outputs generated after each `<act>` |

**An example <act> block with Python code**.

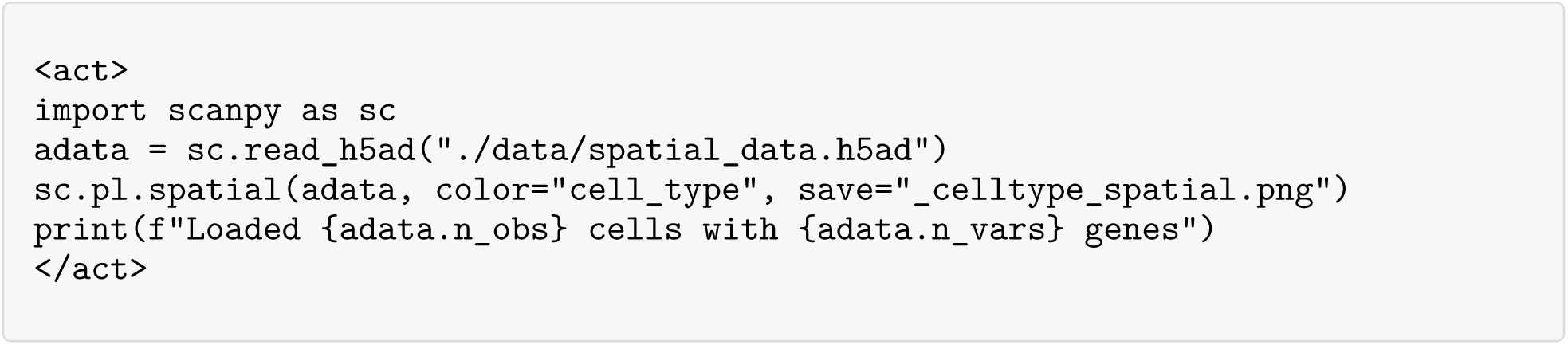

**An example <act> block with Bash commands**.

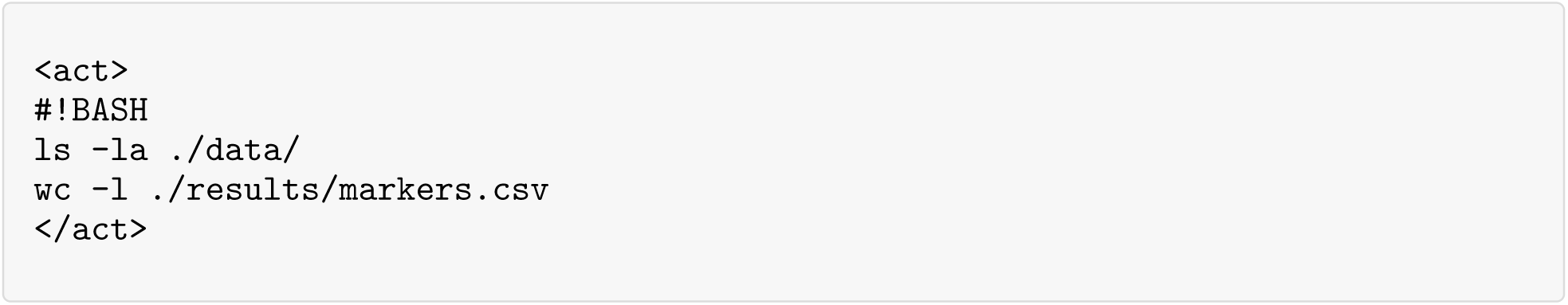

**An example <observation> block**.

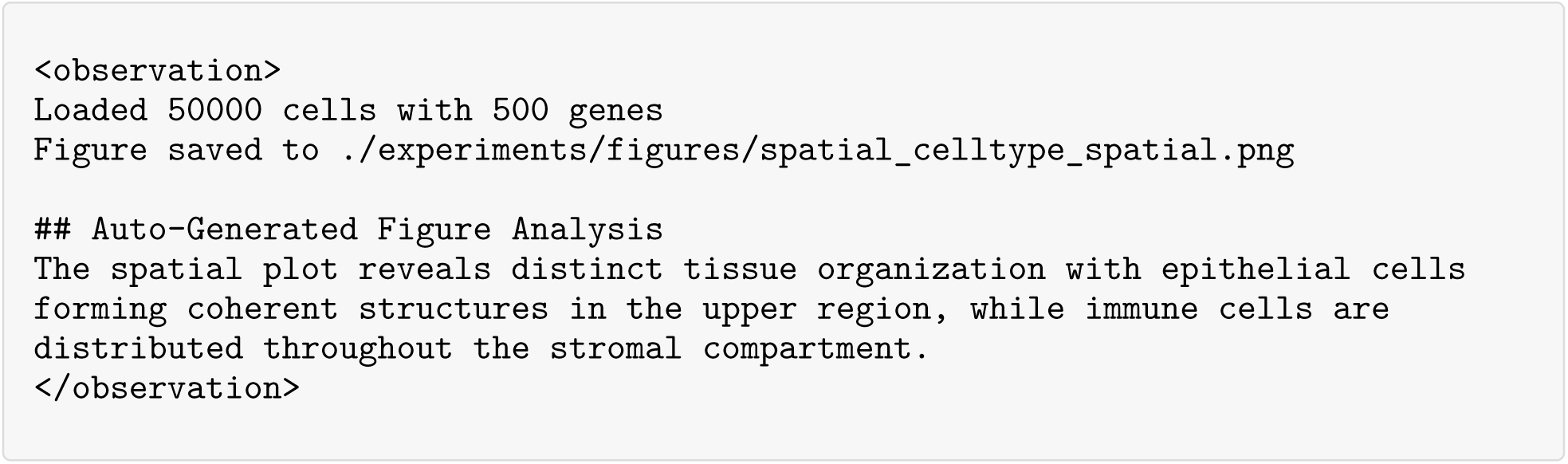

**An example <conclude> block**.

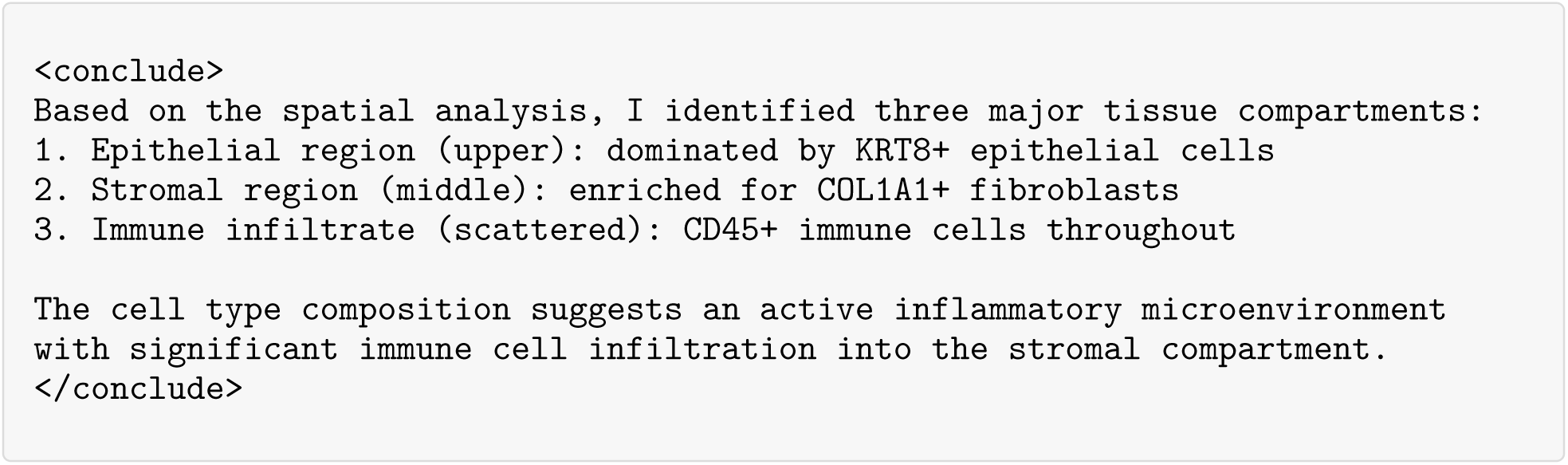

##### Routing and termination

A lightweight router examines each response using regular-expression matching and dispatches execution accordingly. Responses containing `<act>` are sent to the execution node, whereas responses containing `<conclude>` terminate the run. If neither tag is present, the system returns to the planning stage and prompts the model to continue. To improve robustness, the parser automatically closes unclosed tags before parsing. Execution terminates either when the agent emits a `<conclude>` block or when the recursion limit is reached (default: 50 action steps), which serves as a hard stop against runaway execution.

##### Parsing recovery

Malformed or incomplete outputs can arise from provider-side truncation, empty responses, or safety filtering. When required tags are missing, the system injects a corrective prompt requesting either a valid <act> block or a valid <conclude> block and then resumes the loop. If parsing fails twice consecutively, the session terminates to avoid infinite retries.

##### Runtime harness

All actions execute within a controlled environment with timeout protection and structured error handling. Code runs in a persistent Python session, allowing variables, loaded datasets, intermediate tables, and generated artifacts to be reused across steps without reinitialization. Shell execution is also supported when needed for preprocessing or external utilities. Common scientific libraries used in spatial biology, including NumPy, Pandas, Scanpy, Squidpy, Matplotlib, and Seaborn, are pre-imported. Each session writes outputs to a dedicated run directory with deterministic artifact naming. Individual `<act>` blocks are subject to a configurable timeout (default: 1,800 s), and overly long outputs are truncated at 15,000 characters to prevent context overflow. Execution failures are returned to the model inside the corresponding `<observation>` block, enabling error-aware recovery rather than abrupt termination.

**Supplementary Table 2:** Execution harness configuration parameters.

| Parameter | Default | Description |
| --- | --- | --- |
| act_timeout | 1,800 s | Maximum execution time per <act> block |
| recursion_limit | 50 | Maximum graph iterations before forced termination |
| Output truncation | 15,000 chars | Maximum length of execution output |

##### Provenance and logging

SpatialAgent maintains a detailed audit trail of each session through a structured observation log stored as observation_log.jsonl. Each executed step appends a JSON record containing the step index, timestamp, executed code, summarized outputs, and any automatically generated figure interpretations. This log serves as both a provenance record and an artifact index, enabling downstream report generation and claim verification to reference figures, tables, and intermediate results produced during the analysis.

**Supplementary Table 3:** Observation log entry schema.

| Field | Type | Description |
| --- | --- | --- |
| step | Integer | Step index counting executed <act> blocks |
| timestamp | String | ISO 8601 timestamp of execution |
| code_snippet | String | Executed code block |
| result_summary | String | Execution output, return values, or errors |
| figure_interpretations | String | Auto-generated interpretations of newly produced figures |

#### 1.2. Backend interface and deployment

This subsection describes SpatialAgent’s backend interface and deployment options, including provider adapters, backend normalization, tool-call standardization, stopping behavior, and support for locally served open-weight models.

##### Provider adapters

SpatialAgent supports both API-based models and locally hosted open-weight models through a unified backend interface. The make_llm factory function routes model requests to the appropriate provider based on model name patterns and environment configuration.

##### Backend normalization

The backend standardizes message formatting, tool-call parsing, and stopping conditions across providers. Environment variables control endpoint routing. In particular, `CUSTOM_MODEL_BASE_URL` enables custom OpenAI-compatible endpoints for local deployment, whereas provider-specific keys (*e.g.*, `ANTHROPIC_API_KEY`,) route requests to the corresponding cloud services.

##### Universal tool wrapper

Different LLMs produce tool calls with different syntax conventions. Claude commonly emits dict-style calls such as `tool({“param“: “value“})`, whereas Gemini and GPT models often emit keyword-argument calls such as `tool(param=“value”)`. To ensure consistent execution across models, SpatialAgent wraps all tools with a universal adapter that accepts both conventions and normalizes them to the dict format expected by LangChain’s `invoke()` method.

##### Stop sequences

All backends use stop sequences aligned to the Plan-Act-Conclude protocol, specifically </act> and </conclude>, to enforce single-block outputs per response. When one of these closing tags is generated, decoding terminates immediately. This prevents the model from emitting multiple action blocks in one response or hallucinating observation content that should instead be generated by the execution system.

**Supplementary Table 4:** Supported LLM providers and their configurations.

| Provider | Models | Configuration |
| --- | --- | --- |
| Anthropic | Claude Sonnet 4.5/4/3.7, Haiku 4.5, Opus 4.5 | Direct API via `langchain_anthropic` |
| OpenAI | GPT-4o, GPT-5 series | Direct API via `langchain_openai` |
| Azure OpenAI | GPT-4o, GPT-5 series, o3/o3-pro | Azure endpoints with region routing |
| Google | Gemini 2.5 Pro, Gemini 3 Pro | OpenAI-compatible endpoint |
| Local inference | Qwen3-VL-32B, other open-weight models | vLLM or MLX via LiteLLM proxy |

**Supplementary Table 5:** Stop sequence behavior across providers.

| Provider | Stop Parameter | Notes |
| --- | --- | --- |
| Anthropic | `stop_sequences` | Native support |
| OpenAI/Azure | `stop` | Native support; disabled for GPT-5 reasoning models |
| Gemini | `stop` | Via OpenAI-compatible interface |
| Local (vLLM) | `stop` | Via OpenAI-compatible interface |

##### Cost tracking

SpatialAgent tracks token usage and estimated cost through a callback mechanism that records input and output tokens for each LLM call and aggregates them over the session.

##### Local deployment of open-weight LLMs

For privacy-preserving operation or settings without external API access, SpatialAgent supports fully local deployment using open-weight models. We provide two deployment configurations: vLLM for NVIDIA GPUs and MLX for Apple Silicon. The vLLM configuration serves models such as Qwen3-VL-32B-Instruct and Ministral-3-14B-Instruct, with tensor parallelism for multi-GPU servers and a separate lightweight embedding server for tool and skill retrieval. The MLX configuration serves quantized variants of the same models on Apple Silicon. Both configurations use LiteLLM as a proxy layer exposing a unified OpenAI-compatible API, allowing SpatialAgent to switch between API and local backends by changing a single environment variable, without modifications to application code.

**Supplementary Table 6:** Local LLM deployment configurations.

| Platform | Inference Engine | Models | Quantization | Hardware |
| --- | --- | --- | --- | --- |
| NVIDIA GPU | vLLM | Qwen3-VL-32B-Instruct, Ministral-3-14B-Instruct | FP16 | 2–4× A100 (80GB) |
| Apple Silicon | MLX | Qwen3-VL-32B-Instruct-8bit, Ministral-3-14B-Instruct-8bit | 8-bit | M1/M2/M3 Max/Ultra |

#### 1.3. Tool system

This section describes SpatialAgent’s tool system, including the registry contract, structured outputs, error handling, and the CodeAct inspection mechanism for tool adaptation.

##### Tool registry contract

Each tool in SpatialAgent is registered with a standardized contract that enables consistent invocation across different LLM providers. The `Tool` dataclass encapsulates the tool’s identity, interface specification, and executable function.

##### Tool registry entry schema

Tools are defined using LangChain’s @tool decorator with typed annotations that specify parameter constraints, descriptions, and validation rules. The registry automatically extracts JSON Schema from these annotations.

##### Structured outputs and stable filenames

Tools that produce file outputs follow a deterministic naming convention to support artifact tracking and reproducibility. Output filenames incorporate the tool name, relevant parameters, and iteration round where applicable.

**Supplementary Table 7:** Tool output naming conventions.

| Tool Category | Filename Pattern | Example |
| --- | --- | --- |
| Database queries | {database}_{entity}_{round}.csv | pangdb_celltype_1.csv |
| Analysis results | {analysis}_{target}.csv | liana_interactions_macrophage.csv |
| Figures | {plot_type}_{content}.png | spatial_celltype_umap.png |
| Mappings | {method}_{source}_to_{target}.h5ad | tangram_scrna_to_spatial.h5ad |

##### Structured error handling

Tools return errors as structured messages that the LLM can interpret and act upon. Simple tools return error strings prefixed with ERROR: to enable pattern matching. Complex tools that return dictionaries include an error field alongside empty results.

##### CodeAct inspection

Beyond calling tools through their standard interfaces, SpatialAgent supports CodeAct-style inspection that allows the agent to retrieve and examine the entire source code of each tool. When the agent calls `inspect_tool_code`, the system uses Python’s `inspect` module to retrieve the source code and recursively collects all helper functions defined in the same module. This enables the agent to understand implementation details and create targeted modifications.

##### Tool curation principles

SpatialAgent’s tool library was designed following a principled approach that balances comprehensive analytical capability with the flexibility of code generation. Rather than wrapping every possible function, dedicated tools were created only when they provide substantial value beyond what the agent can accomplish through direct code execution.

##### When we create tools

We developed dedicated tools for operations that meet one or more of the following criteria:

- **Multi-step complex pipelines**: Operations requiring coordinated sequences of preprocessing, model fitting, and post-processing steps that benefit from encapsulation. For example, the Tangram mapping pipeline (preprocessing → training → projection) involves multiple interdependent steps with specific data format requirements at each stage.
- **External API calls and database queries**: Operations requiring authentication, rate limiting, error handling, and response parsing for external services. Database tools handle API-specific quirks, implement retry logic with exponential backoff, and heterogeneous response formats.
- **GPU-accelerated deep learning models**: Methods from tools such as scvi-tools, GraphST, and SpaGCN require specific model initialization, training loops, and inference procedures that benefit from standardized wrappers with sensible defaults.
- **Specialized format requirements**: Tools like CellPhoneDB require specific data formatting, metadata structures, and file organization that would be error-prone to recreate each time.
- **Domain expertise encoded in defaults**: Tools where parameter choices encode significant biological or computational knowledge, for example, QC thresholds, normalization strategies, or algorithm-specific hyperparameters validated through benchmarking.

##### What we intentionally exclude

We deliberately chose not to wrap simple, atomic operations as tools. Instead, SpatialAgent provides `execute_python` and `execute_bash` for direct code execution, which covers:

- **Basic preprocessing:** *e.g.*, `sc.pp.normalize_total()`, `sc.pp.filter_genes()`. *Rationale:* Single function calls with intuitive parameters.
- **Differential expression:** *e.g.*, `sc.tl.rank_genes_groups()`. *Rationale:* Well-documented, single-step analysis.
- **Simple statistics:** *e.g.*, NumPy/Pandas operations (mean, std, groupby). *Rationale:* Standard library functions.
- Basic visualization: *e.g.*, *sc.pl.umap()*, *sc.pl.violin()*. *Rationale:* Direct plotting calls.
- **Data manipulation:** *e.g.*, subsetting, merging, reshaping `AnnData`. *Rationale:* Straightforward `AnnData` operations.

This design philosophy ensures that SpatialAgent provides maximum value where tools matter most: complex, expert-level analyses requiring multi-step coordination. For simpler operations, we instead leverage the LLM’s code generation for flexibility and transparency.

##### Tool Categories

The 72 tools are organized into 13 functional categories spanning the complete spatial transcriptomics analysis workflow: database queries (9), literature research (7), core spatial analytics (8), spatial mapping (5), cell-cell communication via CellPhoneDB (5) or LIANA (5), spatial statistics (8), deconvolution (4), spatial domain detection (2), trajectory inference (5), multimodal integration (6), interpretation and code execution (6), and autonomous sub-agents (2).

##### Databases and reference query tools

Multiple databases were integrated to support retrieving reference datasets and conduct basic analysis, leveraging their structured metadata and large-scale annotations.

- **search_czi_datasets** searches the Chan-Zuckerberg Initiative (CZI) CELLxGENE Census for reference single-cell datasets matching a query containing tissue, condition, and organism information (40). The tool converts both the query and dataset metadata into dense vector embeddings using OpenAI’s text-embedding-3-small model, then ranks datasets by cosine similarity to return the top-k most relevant matches. Embedding vectors are cached with checksum validation to optimize repeated queries. CELLxGENE was accessed via its Python API at https://chanzuckerberg.github.io/cellxgene-census/ on November 26, 2024. The retrieved database comprises 135, 560, 133 total cell profiles from 1003 datasets, including 71,730,590 unique cell profiles, annotated across 780 distinct cell types, 63 tissues, from 22,009 human donors and 4,752 mouse models.
- **extract_czi_markers** processes CZI reference datasets to extract cell types and their marker genes. For each cell type in the dataset, the tool queries the CellGuide REST API to retrieve both computational and canonical marker genes, normalizing gene names by case convention (uppercase for human, title case for mouse). Results are saved as CSV files containing cell type identifiers, cell counts, and up to 100 marker genes per cell type.
- **download_czi_reference** downloads scRNA-seq reference data from CELLxGENE for use in Harmony-based label transfer. The tool filters out zero-expression genes, removes cells with unknown cell types, and excludes cell types with fewer than 10 cells to ensure reference quality. Data are converted to proper h5ad format with categorical annotations for efficient storage.
- **search_panglao** queries the PanglaoDB database for marker genes of specified cell types. PanglaoDB database is a web-based resource for exploring mouse and human single-cell RNA-seq data, integrating over 1,000 experiments with more than 4 million cell profiles (41). The tool uses semantic embedding-based matching (OpenAI’s text-embedding-3-small or Qwen3-Embedding-0.6B) to handle variations in cell type nomenclature. The database was downloaded in TSV format from https://panglaodb.se/markers.html on May 20, 2024 (last update March 27, 2020 (10:44:00 CET)). The database contains 8,286 associations, spanning 178 cell types, 4,679 gene symbols, and 29 tissues.
- **search_cellmarker2** queries the CellMarker 2.0 database, which provides manually curated markers for 2,578 cell types across 656 tissues in humans and mice, totaling 83,361 tissue-cell type-marker associations (42), across 26 marker types (*e.g.*, protein-coding genes, lncRNAs) derived from 24,591 studies. The tool performs hierarchical matching (species → tissue → cell type) using semantic embeddings, filtering results by organism and tissue context to return tissue-specific marker associations. The database was downloaded in csv format from http://www.bio-bigdata.center/CellMarker_download.html on Oct 20, 2024.
- **query_tissue_expression** queries the ARCHS4 database via the gget library to retrieve tissue-specific gene expression patterns. For a given gene symbol, the tool returns median TPM values across up to 50 tissues, enabling validation of marker gene tissue specificity.
- **query_celltype_genesets** retrieves cell type-specific gene sets from Enrichr using the ‘PanglaoDB Augmented 2021‘ library. Given a tissue type (brain, liver, lung, heart, kidney, pancreas, skin, or intestine), the tool performs seed gene-based enrichment analysis to return the top-k cell type signatures with associated p-values.
- **validate_genes_expression** validates whether a list of candidate genes are expressed in a target tissue by querying ARCHS4 expression data. The tool ranks the top 20 expressing tissues for each gene and uses string matching to verify target tissue presence, returning a summary of validated versus non-expressed genes with TPM values.
- **query_disease_genes** queries disease-associated genes from multiple sources including the GWAS Catalog REST API and OpenTargets GraphQL API. The tool supports multi-database querying with automatic fallback, implements exponential backoff retry for rate limiting (1, 2, 4 seconds), and aggregates genes across sources with confidence scoring to return ranked gene lists by association strength.

##### Literature research tools

- **query_pubmed** searches PubMed using the Biopython Entrez interface. The tool implements query fallback logic (retrying with simplified queries by removing the last word) and rate limiting (1 second between retries) to handle failed searches. Results include PubMed IDs, titles, journal names, and abstracts truncated to 1,500 characters.
- **query_arxiv** searches arXiv preprints using the official arxiv Python library. Results are sorted by relevance and include titles, summaries (truncated to 1,500 characters), arXiv IDs, and publication dates.
- **search_semantic_scholar** searches Semantic Scholar using their free API, returning paper metadata including titles, abstracts, author lists (first 3 authors plus “et al.”), publication year, citation counts, and journal names.
- **web_search** provides unified web search supporting multiple providers (Anthropic, OpenAI, Google) with automatic provider detection from model names. The tool returns structured results with citations and supports domain filtering for focused searches.
- **extract_url_content** extracts text content from webpages using BeautifulSoup4. The tool removes non-content elements (script, style, navigation, header, footer, aside, iframe) and prioritizes content from semantic HTML tags (main, article, body) with paragraph-based extraction.
- **extract_pdf_content** extracts text from PDF files using PyPDF2. The tool supports both direct PDF URLs and automatic PDF link detection from HTML pages, performing page-by-page text extraction with whitespace normalization.
- **fetch_supplementary_from_doi** retrieves supplementary materials for papers given their DOI. The tool resolves DOIs via CrossRef, implements exponential backoff for rate limiting (max 3 retries), and uses keyword matching to identify and download supplementary files (limited to 5 files per paper).

##### Core spatial analytics tools

- **preprocess_spatial_data** performs standardized preprocessing of spatial transcriptomics data using the Scanpy pipeline. The tool applies quality control filtering (min_genes=5, min_counts=10, min_cells=5), normalization (normalize_per_cell followed by log1p transformation), and dimensionality reduction (PCA → k-NN graph → UMAP embedding).
- **harmony_transfer_labels** transfers cell type labels from scRNA-seq reference data to spatial data via Harmony (43) integration. The tool performs proportional reference sampling to match spatial cell counts, computes 30-component PCA, applies Harmony batch correction, and trains an MLP classifier (hidden layers: 100, 50; max iterations: 500) to transfer labels.
- **run_utag_clustering** performs spatial domain identification using the UTAG algorithm, which integrates spatial neighborhood graphs with gene expression profiles. The tool supports per-sample clustering with configurable maximum neighbor distance, minimum cluster size (default: 100 cells), and multiple Leiden resolutions (default: 0.05, 0.1, 0.3). Small clusters are automatically removed and their cells reassigned to maintain a minimum niche threshold (default: 5 niches).
- **aggregate_gene_voting** aggregates marker genes by group using Scanpy’s rank_genes_groups with the Wilcoxon test, extracting the top 20 marker genes per group for downstream analysis.
- **infer_dynamics** performs differential gene expression analysis across experimental conditions using statistical testing to identify condition-specific gene expression patterns.
- **summarize_conditions** provide LLM-powered biological summarization of experimental conditions, identified cell types, and tissue region compositions respectively.
- **summarize_celltypes** provide LLM-powered biological summarization of experimental conditions, identified cell types, and tissue region compositions respectively.
- **summarize_tissue_regions** provide LLM-powered biological summarization of experimental conditions, identified cell types, and tissue region compositions respectively.

##### Tangram spatial mapping tools

- **tangram_preprocess** prepares single-cell and spatial data for mapping by identifying shared genes, filtering for informative features, and formatting data structures for the Tangram model.
- **tangram_map_cells** learns optimal cell-to-spot mapping using a conditional variational autoencoder (cVAE) that optimizes gene expression correlation between mapped cells and spatial spots.
- **tangram_project_annotations** transfers cell type labels from the reference to spatial coordinates using the learned mapping, computing posterior probabilities for each cell type at each spatial location.
- **tangram_project_genes** imputes unmeasured genes in spatial data by projecting gene expression from the single-cell reference using the learned cell-to-spot mapping.
- **tangram_evaluate** provides quantitative assessment of mapping quality by computing correlation metrics between projected and observed gene expression.

##### CellPhoneDB cell-cell communication (ccc) tools

- **cellphonedb_prepare** formats AnnData objects for CellPhoneDB analysis by extracting normalized expression matrices and cell type annotations in the required input format.
- **cellphonedb_analysis** performs statistical inference of significant ligand-receptor interactions using permutation testing to assess interaction specificity between cell type pairs.
- **cellphonedb_degs_analysis** extends interaction analysis to identify differentially expressed ligand-receptor pairs between experimental conditions.
- **cellphonedb_filter** filters interaction results by specificity thresholds and statistical significance to identify the most biologically relevant interactions.
- **cellphonedb_plot** generates publication-ready visualizations of interaction networks, including dot plots and heatmaps of significant interactions.

##### LIANA multi-method ccc tools

- **liana_inference** runs multiple ligand-receptor inference methods (CellPhoneDB, Connectome, NATMI, SingleCellSignalR, and others) and computes consensus rankings using robust rank aggregation.
- **liana_spatial** performs spatially-aware ligand-receptor analysis using Moran’s bivariate statistic to identify interactions that show spatial co-localization patterns.
- **liana_misty** applies MISTy (Multi-view Intercellular SpaTial modeling framework) for multi-view learning that integrates intrinsic, juxtacrine, and paracrine views of cell-cell communication.
- **liana_tensor** combines LIANA with Tensor-Cell2Cell for multi-sample interaction analysis. The tool runs per-sample rank aggregation, then applies non-negative tensor factorization (rank=3) to decompose condition-specific interaction patterns across samples. Organism is auto-detected from gene name formatting (mouse: title case; human: uppercase).
- **liana_plot** generates visualizations of interaction networks and patterns, including chord diagrams and ranked interaction plots.

##### Squidpy spatial statistics tools

- **squidpy_spatial_neighbors** constructs spatial neighborhood graphs using k-nearest neighbors in spatial coordinates or Delaunay triangulation, enabling downstream spatial statistics.
- **squidpy_nhood_enrichment** tests for significant cell type co-localization in spatial neighborhoods using permutation-based enrichment analysis.
- **squidpy_co_occurrence** analyzes spatial co-occurrence patterns at multiple distance scales, computing pairwise co-localization metrics between cell types.
- **squidpy_spatial_autocorr** computes spatial autocorrelation statistics (Moran’s I and Geary’s C) to identify spatially variable genes that show non-random spatial expression patterns.
- **squidpy_ripley** computes Ripley’s K function and the normalized L-function to characterize spatial point patterns and test against complete spatial randomness (CSR).
- **squidpy_centrality** calculates network centrality metrics (betweenness, degree, closeness) on the spatial neighborhood graph to identify spatially important cell populations.
- **squidpy_interaction_matrix** quantifies cell type interaction frequencies by computing the observed-to-expected ratio of cell type pairs in spatial neighborhoods.
- **squidpy_ligrec** performs spatially-constrained ligand-receptor analysis using the CellPhoneDB database with permutation testing restricted to spatial neighbors.

##### scvi-tools spatial deconvolution tools

- **destvi_deconvolution** performs multi-resolution deconvolution using DestVI (Destination Variational Inference), which provides uncertainty quantification for cell type proportions at each spatial location.
- **cell2location_mapping** estimates absolute cell abundances using a hierarchical Bayesian model that accounts for platform-specific technical effects and provides posterior distributions over cell counts.
- **stereoscope_deconvolution** applies transfer learning from single-cell reference data to spatial spots using a probabilistic model that learns cell type-specific expression profiles.
- **gimvi_imputation** performs cross-modality gene imputation using gimVI, enabling prediction of unmeasured genes in spatial data.

##### Spatial domain detection tools

- **spagcn_clustering** applies SpaGCN (Spatial Graph Convolutional Network), which integrates gene expression with histology image features using graph convolutional networks to identify spatially coherent domains.
- **graphst_clustering** uses GraphST, a self-supervised learning framework that combines graph neural networks with contrastive learning for spatial domain identification with automatic cluster detection.

##### Trajecthory inference tools

- **scvelo_velocity** estimates RNA velocity using scVelo’s dynamical systems modeling, computing the rate of change of gene expression from spliced and unspliced transcript ratios.
- **scvelo_velocity_embedding** visualizes the velocity field in low-dimensional embeddings (UMAP/t-SNE) using stream plots that show predicted future cell states.
- **cellrank_terminal_states** identifies terminal cell states using CellRank’s Markov chain analysis, computing stationary distributions to find absorbing states in the differentiation landscape.
- **cellrank_fate_probabilities** computes cell fate probabilities using absorption probability calculations, assigning each cell a probability of reaching each identified terminal state.
- **paga_trajectory** performs partition-based graph abstraction (PAGA) using Scanpy, constructing a coarse-grained graph of cell type relationships with pseudotime ordering.

##### Multimodal integration tools

- **totalvi_integration** performs joint modeling of RNA and protein (CITE-seq) data using totalVI, a variational autoencoder that learns a shared latent space across modalities.
- **multivi_integration** integrates multiple data views using multiVI, which handles missing modalities and batch effects in multi-modal single-cell data.
- **mofa_integration** applies Multi-Omics Factor Analysis (MOFA+) to discover shared and modality-specific sources of variation using non-negative matrix factorization.
- **scanpy_bbknn** performs batch-balanced k-nearest neighbor graph construction for batch effect correction while preserving biological variation.
- **scanpy_ingest** projects new data onto an existing reference embedding using k-NN mapping, enabling annotation transfer without full re-integration.
- **scanpy_score_genes** computes gene set scores by calculating mean expression of gene signatures, useful for pathway activity scoring.

##### Interpretation and support tools

- **annotate_cell_types** performs hierarchical two-level batch cell type annotation. The tool first clusters cells using multi-resolution Leiden clustering with automatic resolution selection, then applies a two-level annotation strategy: (1) broad category assignment to all clusters in a single LLM call for efficiency, and (2) refinement to specific subtypes within each category for accuracy. The tool uses tissue-type-aware ontology filtering and incorporates transferred cell type compositions as annotation hints.
- **annotate_tissue_niches** annotates spatial tissue niches with optional anatomical reference. The tool generates composite niche plots (all niches colored in one image per sample), performs per-sample batch annotation (one LLM call per sample), and applies multi-sample consensus merging (one final LLM call). Features include spatial coordinate normalization (0-100 scale) for anatomical reasoning and patient-perspective orientation handling for medical imaging.
- **interpret_figure** provides vision LLM-based interpretation of scientific figures. The tool resizes images to meet API limits (<4MB), auto-detects MIME types, and generates concise 3-5 line biological interpretations with binary observations and bullet-point findings.
- **execute_python** provides a stateful Python REPL with pre-imported libraries (numpy, pandas, scanpy, squidpy, matplotlib, seaborn) and injected tool functions. The tool supports persistent variable state across calls, automatic markdown code fence stripping, new image file tracking for auto-interpretation, and contextualized error reporting with line numbers.
- **execute_bash** executes shell commands with a 60-second timeout, providing access to system utilities with environment variables ($SAVE_PATH, $DATA_PATH) for file operations.
- **inspect_tool_code** retrieves the complete source code of any registered tool including all helper functions and dependencies. The tool uses AST-based recursive analysis to identify function calls, collects transitive dependencies via inspect.getsource(), and returns code in topological order. This enables the agent to understand tool implementations and create modified versions for non-standard use cases.

##### Sub-agent tools

- **report_subagent** generates publication-quality research reports through a multi-pass analysis of saved artifacts. The tool performs six passes: (1) inventory artifacts (observation logs, figures, CSVs, JSONs), (2) interpret all figures with vision LLM (max 15 figure files), (3) analyze CSV tables with statistics (ma× 10 table files), (4) extract JSON summaries (max 5 summary files), (5) format observation history (max 20 steps), and (6) generate comprehensive report. The output includes executive summary, introduction, methods, results with figure references, discussion, conclusions, limitations, future directions, and a figure gallery appendix. Reports are generated in both markdown and styled HTML formats.
- **verification_subagent** verifies analysis conclusions against saved evidence through a five-pass workflow: (1) inventory artifacts, (2) sample key figures for visual verification (5 stratified samples), (3) analyze CSVs for quantitative evidence (max 5), (4) extract observation history (last 10 steps), and (5) generate verification report. The report provides per-claim assessment (Supported, Partially, Not supported, Cannot verify), scores for query alignment, evidence support, completeness, and scientific rigor (each scored on a 10-point scale), and an overall verdict with recommendations (ACCEPT / REVISE / MAJOR REVISION NEEDED).

#### 1.4. Retrieval and skill system

This subsection describes how SpatialAgent selects a task-relevant working set of tools and reusable skills for each query.

##### Tool retrieval

Tools are embedded and retrieved by cosine similarity using Qwen3-Embedding-0.6B (768 dimensions) via the sentence-transformers library. When a user query arrives, it is embedded and compared against pre-computed tool embeddings using normalized dot product. The top-*k* most similar tools (default *k* = 20) are retrieved as candidates.

For semantically subtle queries where embedding similarity may be insufficient, we optionally apply an LLM-based tool selector. The LLM receives the full tool catalog and selects the most relevant subset based on the specific task requirements. This two-stage approach balances efficiency (fast local embeddings) with accuracy (nuanced LLM reasoning).

**Supplementary Table 8:**
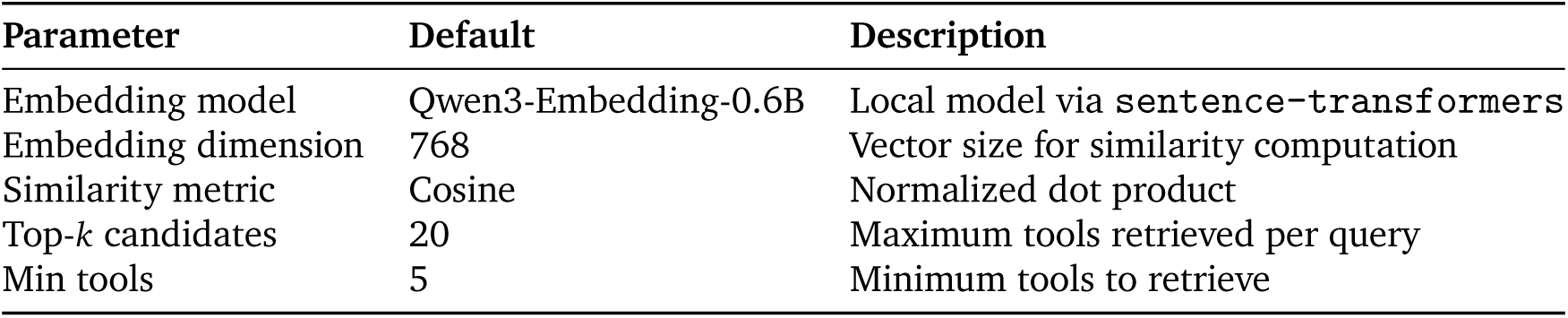
Tool retrieval configuration.

| Parameter | Default | Description |
| --- | --- | --- |
| Embedding model | Qwen3-Embedding-0.6B | Local model via `sentence-transformers` |
| Embedding dimension | 768 | Vector size for similarity computation |
| Similarity metric | Cosine | Normalized dot product |
| Top-$k$ candidates | 20 | Maximum tools retrieved per query |
| Min tools | 5 | Minimum tools to retrieve |

A set of core tools are always loaded regardless of the retrieval results, ensuring fundamental capabilities. These include `execute_python` and `execute_bash` for code execution, `inspect_tool_code` for source code inspection, `query_pubmed` for literature search, and `web_search` for online search.

##### Skill routing

We maintain a library of 17 reusable skills (workflow templates) stored as markdown files. Each skill encodes a multi-step standard operating procedure for common spatial transcriptomics tasks.

A lightweight LLM classifier decides whether a query warrants a skill-based workflow or can be executed *ad hoc*. The classifier receives the query along with descriptions of all available skills and outputs either a matching skill name or `NO_MATCH`. For queries under 50 characters or simple commands, the classifier typically returns `NO_MATCH` to allow flexible ad-hoc execution. When a skill is selected, tools referenced inside the skill are automatically extracted and loaded alongside retrieved tools. The extraction uses pattern matching to identify tool names in backticks (*e.g.*, `search_panglao`) and known tool prefixes.

##### Skill generation from memory

After successful novel analyses, the generate_skill_from_memory function can distill the execution trace into a reusable markdown SOP to expand the library. The function extracts all <act> blocks from the conversation history and uses an LLM to synthesize a structured workflow template capturing the essential steps, tools used, and decision points. This mechanism enables the skill library to grow organically from successful analyses, capturing institutional knowledge and best practices discovered during real usage.

**Supplementary Table 9:**
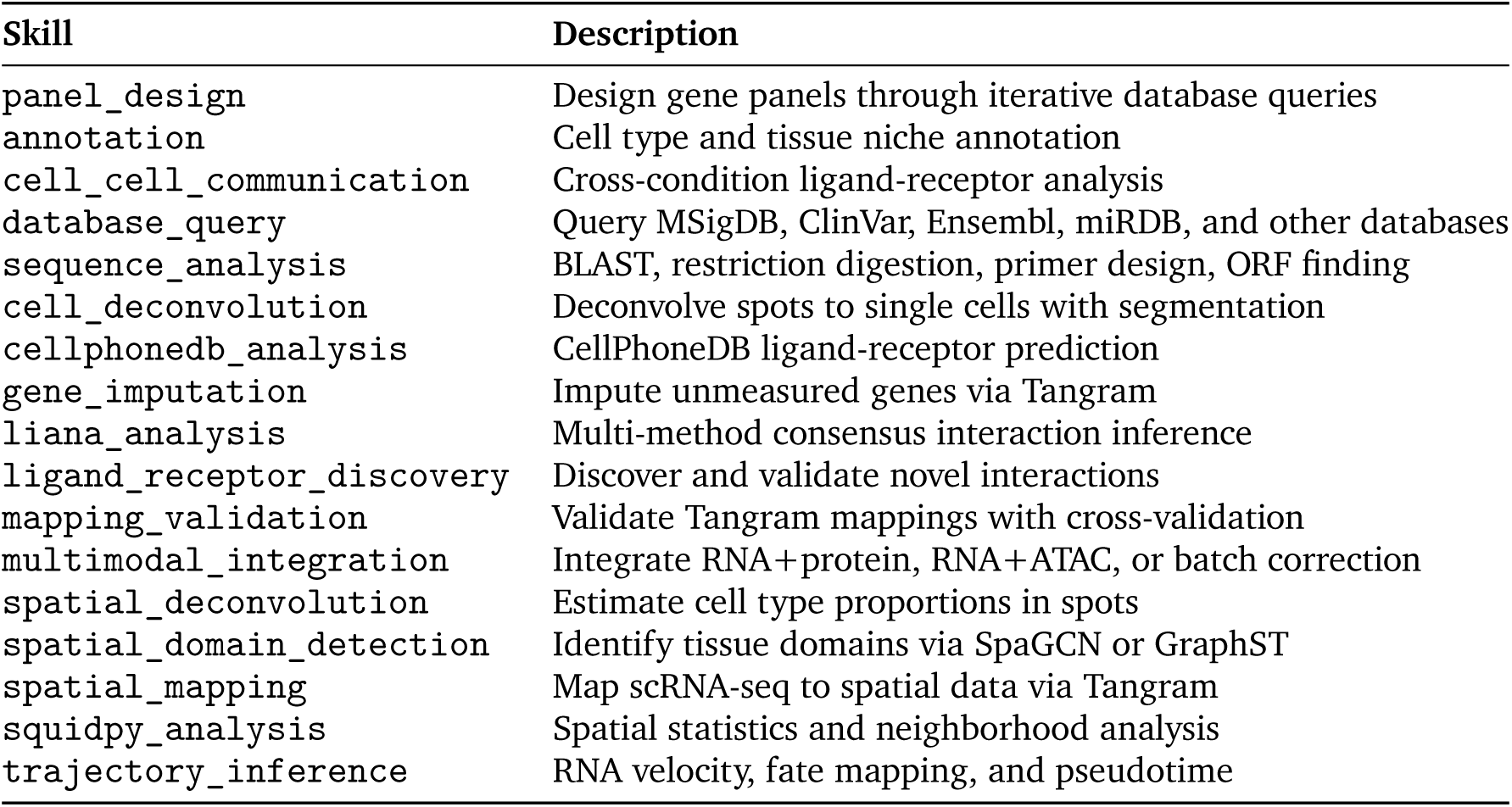
Skill library contents.

| Skill | Description |
| --- | --- |
| panel_design | Design gene panels through iterative database queries |
| annotation | Cell type and tissue niche annotation |
| cell_cell_communication | Cross-condition ligand-receptor analysis |
| database_query | Query MSigDB, ClinVar, Ensembl, miRDB, and other databases |
| sequence_analysis | BLAST, restriction digestion, primer design, ORF finding |
| cell_deconvolution | Deconvolve spots to single cells with segmentation |
| cellphonedb_analysis | CellPhoneDB ligand-receptor prediction |
| gene_imputation | Impute unmeasured genes via Tangram |
| liana_analysis | Multi-method consensus interaction inference |
| ligand_receptor_discovery | Discover and validate novel interactions |
| mapping_validation | Validate Tangram mappings with cross-validation |
| multimodal_integration | Integrate RNA+protein, RNA+ATAC, or batch correction |
| spatial_deconvolution | Estimate cell type proportions in spots |
| spatial_domain_detection | Identify tissue domains via SpaGCN or GraphST |
| spatial_mapping | Map scRNA-seq to spatial data via Tangram |
| squidpy_analysis | Spatial statistics and neighborhood analysis |
| trajectory_inference | RNA velocity, fate mapping, and pseudotime |

#### 1.5. Automated figure interpretation and reporting/verification sub-agents

This subsection describes SpatialAgent’s auxiliary modules for automated figure interpretation, conclusion verification, and publication-style report generation. Automated figure interpretation is implemented as a lightweight LLM call within the main execution loop, whereas reporting and verification are implemented as separate sub-agents with their own tools and multi-step procedures.

##### Automated figure interpretation

After each action, the runtime detects newly generated figures by scanning monitored directories before and after execution. The system maintains a list of existing images before code runs and identifies new files (PNG, JPG, JPEG) created during execution. For each new figure, it infers plot context from the executed code via regex pattern matching to extract plot type, variables being visualized, axis labels, and gene names. It then passes the image together with this context to a vision-capable LLM and appends a 3–5 line biological interpretation to the observation stream. This feature is enabled via the `auto_interpret_figures` parameter (default: `True`) and skips PDF files, which are less suitable for direct vision-based interpretation.

**Supplementary Table 10:** Auto-figure interpretation configuration.

| Parameter | Default | Description |
| --- | --- | --- |
| `auto_interpret_figures` | True | Enable automatic figure interpretation |
| Supported formats | PNG, JPG, JPEG | Image formats processed by vision LLM |
| Skipped formats | PDF | Files excluded from interpretation |
| Context extraction | Regex | Plot type, variables, titles from code |
| Output length | 3–5 lines | Concise biological interpretation |

##### Verification Sub-agent

The verification sub-agent performs a five-pass analysis to cross-check conclusions against saved evidence. It inventories all artifacts generated during the session, including observation logs, figures, CSV files, and JSON summaries; samples key figures for visual verification using a vision-capable LLM; reads CSV outputs to assess quantitative claims; reconstructs the analysis history from observations; and produces a structured verification report. The verifier assigns claim-level support labels to specific conclusions and returns prioritized recommendations together with an overall verdict of Accept, Revise, or Major Revision Needed.

**Supplementary Table 11:** Verification rubric dimensions.

| Dimension | Score Range | Assessment Criteria |
| --- | --- | --- |
| Query alignment | 0–10 | How well conclusions address the research question |
| Evidence support | 0–10 | Whether conclusions are supported by figures and data |
| Completeness | 0–10 | Whether sufficient analyses were performed |
| Scientific rigor | 0–10 | Alternative interpretations, confounders, limitations |

**Supplementary Table 12:** Claim-level support labels.

| Label | Meaning |
| --- | --- |
| ✓ Supported | Evidence clearly supports the claim |
| ! Partially supported | Some evidence supports, but gaps exist |
| × Not supported | Evidence contradicts or does not support claim |
| ? Cannot verify | Insufficient evidence to evaluate |

##### Report sub-agent

The report sub-agent performs a six-pass synthesis to generate a publication-style research report. It inventories available artifacts, interprets selected figures using a vision-capable LLM, analyzes CSV tables, reads JSON summaries, reconstructs the analysis narrative from logged observations, and synthesizes these materials into a structured markdown report. This separation between primary analysis and downstream reporting allows the final document to explicitly reference generated artifacts and intermediate results.

**Supplementary Table 13:** Report sub-agent pass structure.

| Pass | Action | Output |
| --- | --- | --- |
| 1 | Inventory artifacts | List of figures, CSVs, JSONs, observations |
| 2 | Interpret figures | 200–300 word analysis per figure (up to 15) |
| 3 | Analyze CSV tables | Statistics, distributions, key values |
| 4 | Read JSON summaries | Structured insights |
| 5 | Extract observation history | Analysis narrative and methods |
| 6 | Synthesize report | Publication-style markdown document |

##### Report structure and image handling

The generated report follows a standard scientific structure, including an executive summary, introduction, methods, results, discussion, conclusions, and a figure gallery. To satisfy API image-size constraints during report generation, oversized images are automatically resized before submission while preserving aspect ratio and visual fidelity.

#### 1.6. OpenWebUI integration and session persistence

This subsection describes the lightweight OpenWebUI integration layer used to expose SpatialAgent as an interactive chat agent, together with the mechanism for session persistence across turns.

##### Input handling

To provide an interactive user experience, we deploy a minimal OpenWebUI [pipe](https://docs.openwebui.com/features/pipelines/pipes) wrapper around SpatialAgent. This wrapper intercepts incoming chat requests and forwards them to the agent without modifying the user input.

##### Output rendering

The wrapper is responsible for converting agent outputs into a format suitable for OpenWebUI, which expects markdown. It parses structured agent tags, including `<act>`, <observation>, and <conclude>, and groups them into logical execution steps for display in the frontend. As each step is produced by the agent, the wrapper streams the formatted content to the OpenWebUI interface.

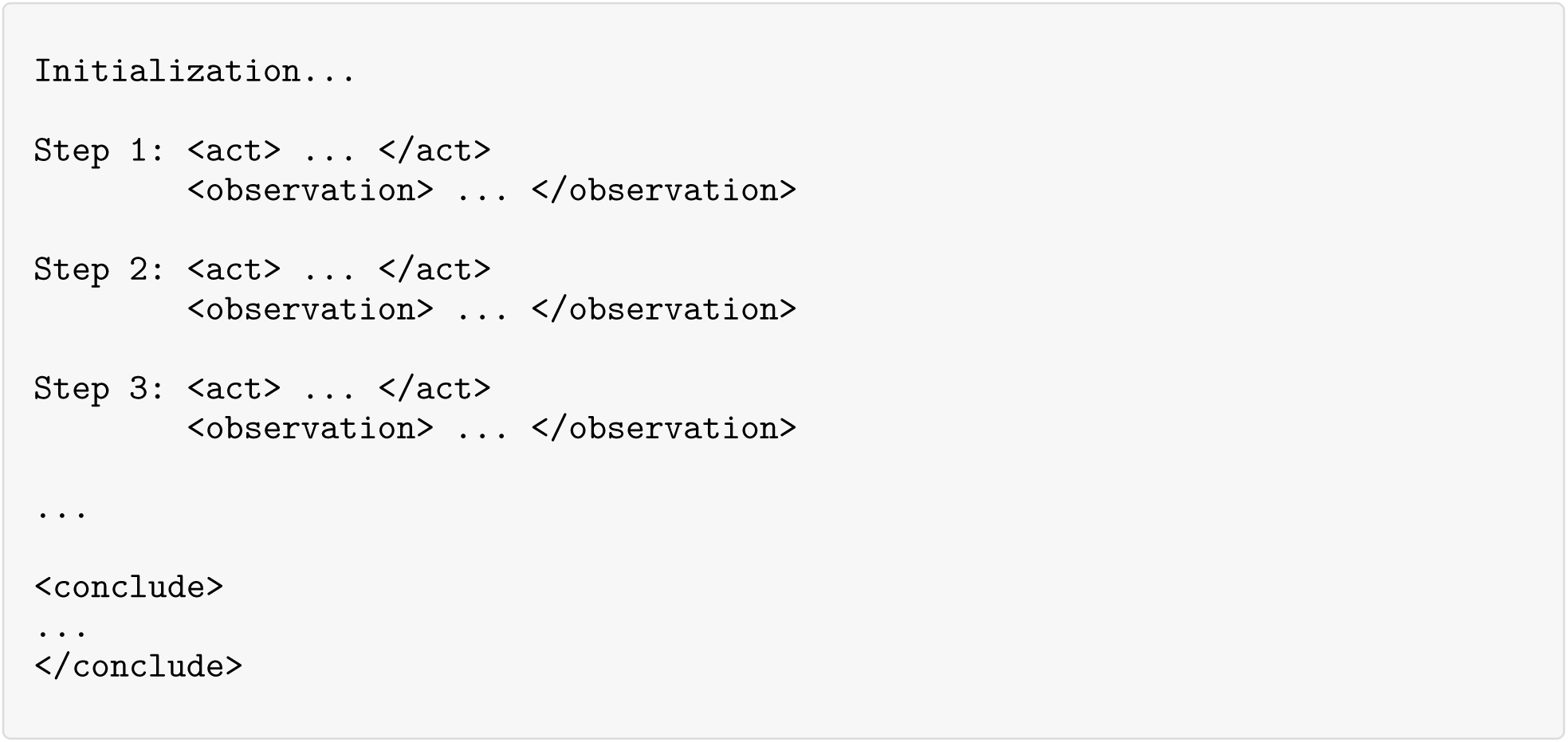

##### Session persistence and lifecycle

SpatialAgent uses LangGraph’s [SqliteSaver](https://reference.langchain.com/python/langgraph/checkpoints#langgraph.checkpoint.sqlite.SqliteSaver) to store execution history on disk. Each execution runs in an isolated container. OpenWebUI automatically passes request headers including `X-User-Email` and `X-OpenWebUI-Chat-Id`, which are used to create or resume a specific LangGraph thread. Thread state is persisted to S3 and restored before subsequent executions of the same thread.

To preserve execution context beyond the checkpointed graph state, we also snapshot the working filesystem before and after each execution, identify files created or modified by the agent, and persist these changes to S3 under the corresponding thread identifier. After each run, the execution container is torn down and recreated in preparation for the next interaction.

##### Continuing the conversation

When a user continues a conversation, both the thread checkpoint and the associated filesystem state are restored from S3 into a fresh execution container. The new user message is then processed using the restored state, with `X-User-Email` and `X-OpenWebUI-Chat-Id` used to identify the relevant thread.

### 2. LLM backends, decoding settings, and skill ablations

This section summarizes the ablations used to select the default language-model configuration for SpatialAgent. We evaluated both API-based and open-weight backends, compared decoding settings, and measured the effect of enabling the skill library across representative tasks. Unless otherwise noted, the main experiments used Claude Sonnet 4.5 at temperature *T* = 0.5, selected on the basis of these analyses (Extended Data Figs. 1–4).

This section evaluates how SpatialAgent depends on configuration choices that commonly vary across deployments. Because SpatialAgent executes in a closed loop (planning, tool calls, code execution, and constrained outputs), configuration changes can affect both the final answer and intermediate behaviors (*e.g.*, planning, tool selection, coding decisions, and conclusion). We therefore characterize sensitivity along four axes: LLM backend, sampling, task, and skills, while holding the rest of the system fixed.

For clarity, we organize the analysis in four stages. We first compare API-based and open-weight backends under matched protocols to establish which performance differences are primarily attributable to model capability rather than decoding. We then examine temperature and other decoding controls within each backend, before quantifying how much explicit skill scaffolds improve robustness. We conclude by synthesizing these results into the default configuration used in the main experiments.

#### Factors varied

Unless a factor is explicitly varied, we keep the execution environment, toolset, datasets, prompts, and evaluation code unchanged.

- **LLM backend.** We evaluate API-based models (Sonnet 4.5, Gemini 3 Pro, GPT-4o, GPT-5) and open-weight models served locally (Qwen3-VL-32B-Instruct, Ministral-3-14B-Instruct-2512).
- **Sampling.** For models that expose temperature, we sweep temperature. For GPT-5, temperature is not exposed and we instead sweep `reasoning_effort` (Low/Medium/High). For open-weight models, we additionally evaluate a small set of decoding presets that vary truncation controls (`top_p`, `top_k`); for Qwen3 we also test hybrid decoding that applies different presets to text-only versus image questions.
- **Task.** We consider (i) question-answering (QA) benchmarks (GPQA Diamond, HLE Biology split, LabBench) and (ii) autonomous workflows with objective evaluation metrics (gene panel design and cell-type annotation).
- **Skills.** For workflows, we compare runs with skills enabled versus disabled. Skills provide an ordered scaffold of tool primitives and validation steps intended to improve completion and constraint adherence.

#### Sampling protocol and nondeterminism

For API-based backends, sampling is controlled primarily via temperature (Sonnet 4.5, Gemini 3 Pro, GPT-4o) or via `reasoning_effort` for GPT-5. For open-weight backends served locally, we vary temperature and, where applicable, decoding truncation controls (`top_p`, `top_k`) using a small set of recommended presets; for Qwen3 we additionally evaluate a hybrid preset that applies different decoding parameters to text-only versus image questions. For hosted APIs, *T*=0 does not guarantee deterministic outputs, and we observe run-to-run variation consistent with provider-side nondeterminism. For local serving, outputs can also vary across runs due to stochastic decoding and parallel inference; accordingly, all results are summarized over repeated runs.

#### Tasks and readouts

We report task-level performance with system-level robustness measures.

- **QA benchmarks.** Accuracy on GPQA Diamond, HLE Biology, and LabBench. Where available, we also report cost per GPQA run and latency per GPQA question.
- **Gene panel design.** Accuracy, macro-F1, spatial reconstruction *R*^2^, and wall-clock time, evaluated under the sampling sweep, with and without skills.
- **Cell-type annotation.** Accuracy, class-balanced accuracy, macro-F1, ARI, recall, and coverage (number of annotated types), under the same sweep protocol, with and without skills.

For tasks with hard requirements, we additionally track **constraint adherence**. For panel design, we summarize the distribution of generated panel sizes under the requested gene-count constraint.

#### Completion and failure modes

For each configuration, we report completion rate and representative failure modes (*e.g.*, tool-call errors, formatting violations, missing artifacts, or constraint violations). Some failures arise from infrastructure or provider-side instability (*e.g.*, transient HTTP errors) rather than from sampling, and are reported separately from task metrics when identifiable. We also summarize qualitative differences in execution patterns across backends (*e.g.*, tool usage and coding style) when they are stable across sampling settings.

#### Repetition and aggregation

Each configuration is evaluated with three independent runs and reported as mean ± standard deviation. In plots, the outlined bar denotes the best setting within each subplot. For skills ablations, we provide two complementary summaries: **best vs. best** (each condition at its best sampling setting) and **mean vs. mean** (averaged over all sampled settings), separating peak performance from stability across sampling choices.

#### 2.1. API and open-weight backends

We first compare API-based and open-weight backends because backend choice sets the overall capability ceiling for the downstream workflows. To reduce confounding from decoding, we use temperature sweeps for backends that expose temperature and `reasoning_effort` sweeps (Low/Medium/High) for GPT-5. For each task, we summarize both (i) the best-performing setting within each backend; and (ii) the average performance across the sweep when it is informative. Unless otherwise stated, all results are reported as mean ± s.d. over three independent runs.

##### Cross-backend summary views

Sweep plots are valuable for diagnosing decoding sensitivity within a model, but they can obscure cross-backend gaps by spreading each backend across many settings. We therefore provide compact summaries that compare backends directly under matched sweeps. Specifically, we report:

- *best-vs-best* comparisons that report the best setting for each backend (optionally overlaid with the sweep mean to indicate stability);
- *quality–efficiency* trade-offs for tasks where runtime is recorded (QA latency/cost; panel-design wall time);
- *reliability* views that report completion rate and task-specific validity constraints (panel-size adherence; annotation coverage and taxonomy consistency).

These summaries complement the full sweep figures in Section 2.2.

##### 2.1.1. Backend Comparison on Paper QA Benchmarks

Across both GPQA Diamond and HLE Biology, backend choice dominates decoding choice: the ordering between backends is largely preserved when comparing each backend at its own best setting, and the gap between each backend’s best and sweep mean is generally smaller than the gap between different backends (Supplementary Fig. 1). Among API models, Gemini 3 Pro attains the highest accuracy on both benchmarks, while GPT-5 is consistently competitive but with a larger best–mean separation, reflecting a stronger dependence on the backend-specific compute knob (`reasoning_effort`). Sonnet 4.5 remains robust across its sweep, with a comparatively small drop from best to mean, which is desirable when the same backend must serve many prompts and tasks without per-task tuning. Open-weight models remain below the leading APIs under the same evaluation protocol, and their best–mean gaps can be more pronounced, consistent with decoding presets (`top_p`/`top_k`) affecting reproducibility in addition to mean accuracy.

**Supplementary Figure 1:**
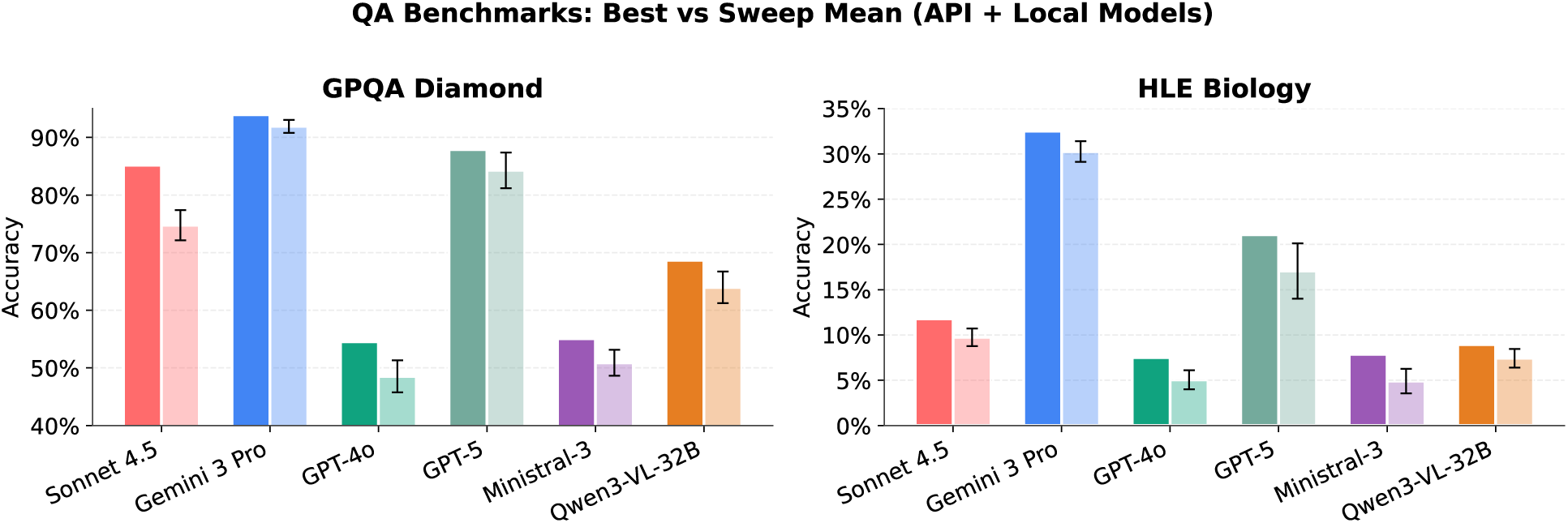
Backend comparison on QA benchmarks under matched sweeps. Best-vs-sweep-mean accuracy (y axis) for each backend (x axis) on GPQA Diamond and HLE Biology. Darker bar: best setting within its backend sweep (temperature, or reasoning_effort for GPT-5); lighter bar: sweep mean, highlighting both peak performance and sensitivity to decoding.

##### 2.1.2. Backend Comparison on Gene Panel Design Without Skills

**Supplementary Figure 2:**
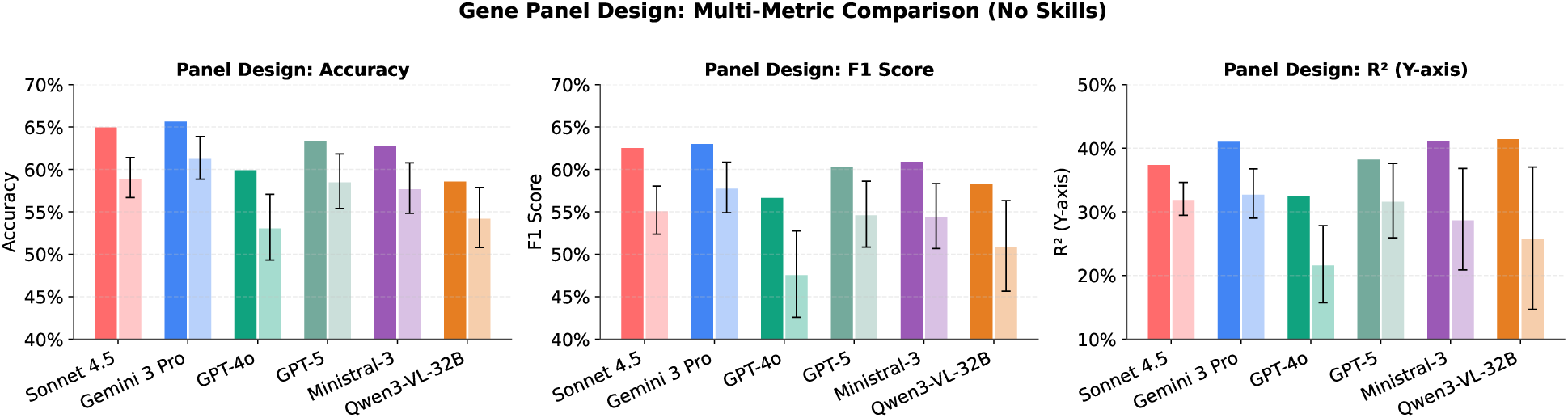
Backend comparison on gene panel design (no skills). Best-vs-sweep-mean performance metrics (y axis) for each backend (x axis) on the gene panel design without skills. Dark bars: best setting for each backend’s sweep; light bars: sweep mean.

Under no-skills execution, backend effects are amplified because errors in instruction following (*e.g.*, returning the wrong number of genes) propagate into downstream evaluation and runtime. Across metrics, the leading APIs maintain an advantage in both accuracy/F1 and spatial *R*^2^, while weaker backends exhibit larger dispersion between best and sweep mean (Extended Data Fig.3), indicating that decoding can modulate performance but cannot erase capability gaps. The spatial metric is more discriminative than accuracy/F1: backends that are relatively close on classification-oriented metrics can separate more clearly on spatial reconstruction, suggesting that fine-grained fidelity is more sensitive to backbone capability and intermediate decision quality.

In practice, the relevant operating point depends on the trade-off between throughput and peak quality. Reliability views further clarify that a substantial fraction of end-to-end error arises from validity failures rather than subtle metric differences. Stronger backends exhibit high panel-size exact-match rates and low deviations from the requested gene count, whereas weaker backends show markedly lower exact-match and larger deviations (Supplementary Fig. 3). These constraint violations can simultaneously depress downstream scores and distort runtime (*e.g.*, under-generation can shorten downstream steps), motivating explicit scaffolds (skills) in later sections.

**Supplementary Figure 3:**
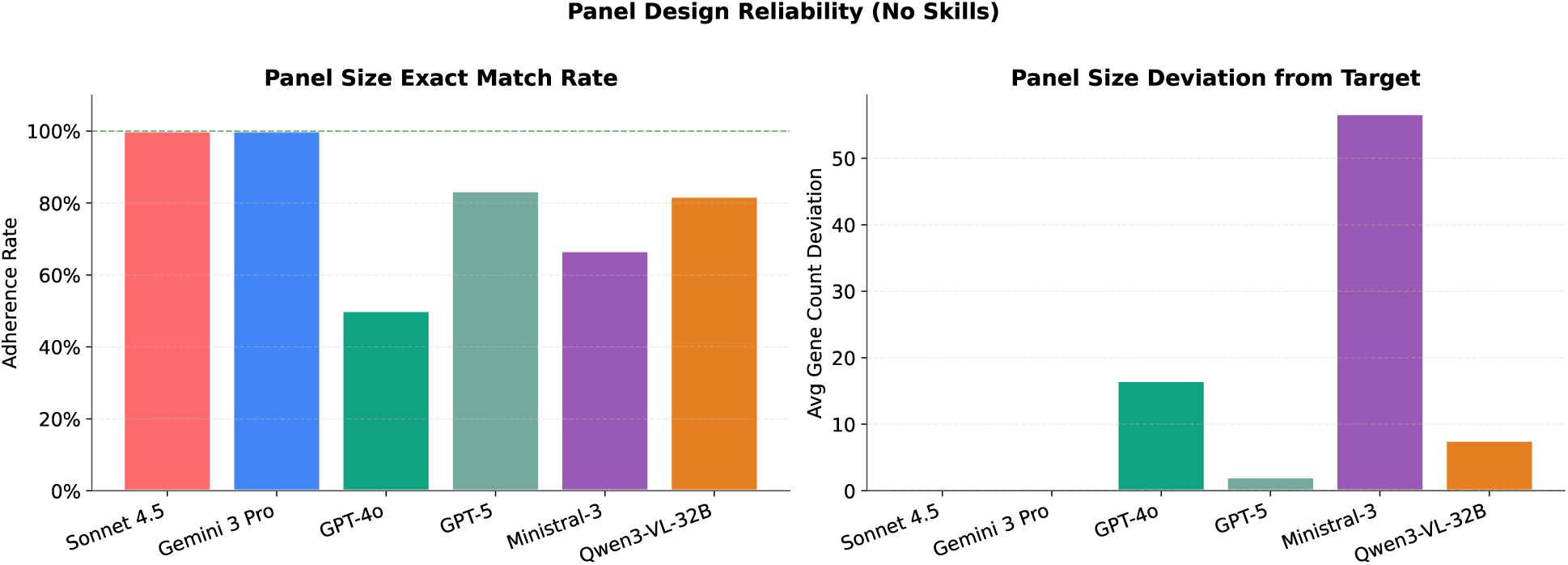
Reliability for gene panel design (no skills). Exact-match rate for the requested panel size (left, y axis) and mean absolute deviation from the target gene count (right, y axis) across backends (x axis) under no-skills execution.

##### 2.1.3. Backend Comparison on Cell-Type Annotation Without Skills

**Supplementary Figure 4:**
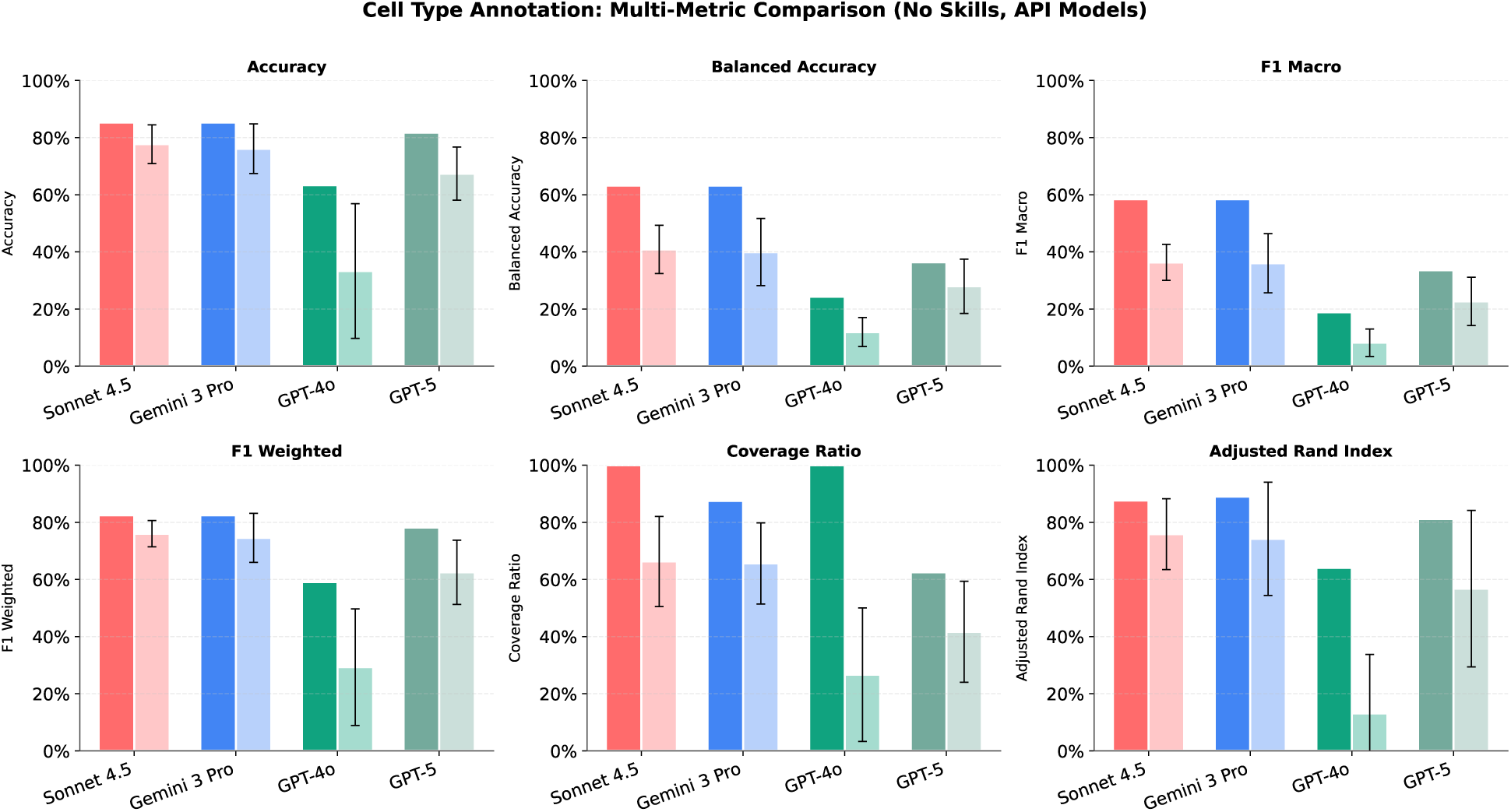
Backend comparison on cell-type annotation (no skills, API backends). Best-vs-sweep-mean performance metrics (y axis) for each API backend (x axis) on cell-type annotation without skills. Dark bars: best setting within each backend’s sweep. Light bars: sweep mean.

For annotation, backend differences are most visible in class-balanced and structure-aware metrics, which penalize terminological drift and inconsistent granularity more strongly than overall accuracy. Across backends, macro-F1 and ARI show larger best–mean gaps than accuracy (Supplementary Fig. 4), consistent with decoding and backend behavior affecting rare types and taxonomy consistency. GPT-5 and Gemini 3 Pro achieve the strongest agreement metrics under their best settings, while Sonnet 4.5 remains competitive with comparatively stable sweep means, supporting use cases where consistent behavior across many runs is preferred over marginal peak gains. GPT-4o remains substantially weaker across metrics, and its coverage-related readouts should be interpreted cautiously: higher apparent coverage can reflect fragmented naming or mixed-granularity outputs rather than improved biological resolution. Together, these results reinforce that backend selection for annotation should be guided by stability on class-balanced and agreement metrics, not only top-line accuracy.

##### 2.1.4. Summary and Practical Implications

Across tasks, backend choice influences peak performance, stability across decoding settings, and validity in closed-loop execution. Two practical lessons emerge from the matched-sweep comparisons. First, for strong backends, performance is often relatively flat across a broad range of decoding settings, so selecting an operating point should prioritize operational considerations (throughput, cost, failure rates, and constraint adherence) rather than chasing small peak gains. Second, for weaker backends, a non-trivial portion of end-to-end error arises from validity failures (constraint violations in panel design; taxonomy drift in annotation), which skills and explicit normalization can mitigate more effectively than additional decoding tuning.

This backend comparison therefore provides the reference frame for the remainder of the section: we next ask how much decoding changes behavior within a fixed backend, and then whether skills can compensate for backend-specific failure modes.

#### 2.2. Temperature and decoding settings

We next hold the backend fixed and isolate sensitivity to temperature and other decoding controls within each model. This analysis asks whether operating-point selection meaningfully changes behavior once a backend has already been chosen. For each LLM backend, we adjust sampling parameters while keeping the task, prompts, tools, evaluations, and execution environment fixed. Results are reported as mean ± s.d. over three independent runs.

##### 2.2.1. QA Benchmarks

We begin with QA benchmarks that do not require tool execution, so the sweep probes decoding effects without confounding from tool selection or multi-step execution. We evaluate GPQA Diamond, HLE Biology, and LabBench using each benchmark’s required answer format and scorer, and we additionally track cost and latency where available. We first summarize each dataset briefly below, with both API-based LLM results across sampling sweeps (Extended Data Fig.1) and open-weight LLM results under the evaluated decoding presets (Tables 14 and 15).

###### GPQA Diamond

GPQA Diamond is a multiple-choice benchmark spanning physics, chemistry, and biology. We use the evaluator-required “final answer” format (“ANSWER: X”) and randomly permute answer choices per question to control for position bias; scoring is based on the extracted final choice.

###### LabBench

LabBench evaluates life-science QA across diverse formats (*e.g.*, tables, figures, protocols) under a unified harness. We use the authors’ prompt template and answer tags to ensure consistent parsing across backends and sampling settings, and we retain the benchmark’s standard retry behavior for transient failures.

###### HLE Biology

The Biology/Medicine subset of Humanity’s Last Exam (HLE) includes both text-only and image-based questions, with a mix of multiple-choice and exact-match formats. We follow the official response structure and use an extraction step for answer normalization on exact-match items, then evaluate text-only and multimodal questions under the same sweep.

**Supplementary Table 14:** Sampling configurations and performance on GPQA Diamond. We report mean accuracy (%) ± SD over three independent runs and average end-to-end latency (sec) per question. “Config” lists (temperature, top_p, top_k) (top_k = −1 disables top-*k* truncation; “–” indicates the parameter is not exposed by the API). For Sonnet 4.5, the second accuracies correspond to Extended Thinking mode (*), where temperature is not user-configurable. GPT-5 was evaluated using the default Medium reasoning effort (**). “reported” rows list results from official sources. For Qwen3 and Ministral3, we **bold** the highest-mean configuration and any others whose mean±SD overlaps with it.

| Name | Config: temperature, top_p, top_k | Accuracy (%) | Latency (sec) |
| --- | --- | --- | --- |
| Qwen3-VL-32B-Instruct | vllm_default: 1.0, 1.0, -1 | **52.0 ± 2.6** | 69.0 |
|  | vllm_greedy: 0, 1.0, -1 | **63.1 ± 4.1** | 97.2 |
| | qwen_vl: 0.7, 0.8, 20 | 61.9 ± 1.3 | 80.6 |
|  | qwen_text: 1.0, 1.0, 40 | **65.5 ± 3.1** | 69.7 |
|  | *reported* | N/A (68.9 for full set) | N/A |
| Ministral-3-14B-Instruct | vllm_default: 1.0, 1.0, -1 | **50.7 ± 4.7** | 30.8 |
| | vllm_greedy: 0, 1.0, -1 | 51.2 ± 0.6 | 36.7 |
| | production: 0.05, 1.0, -1 | 49.5 ± 1.3 | 36.9 |
|  | creative: 0.15, 1.0, -1 | **52.0 ± 2.6** | 31.7 |
|  | *reported* | N/A (71.2 for -Reasoning) | N/A |
| Sonnet 4.5 | best of ours: 0, 0.99, - | 78.0 ± 6.2; **80.1 ± 1.7\*** | 17.4; 53.3* |
|  | *reported* | 83.4 | N/A |
| Gemini 3 Pro | **best of ours:** 0.3, 0.95, 64 | 92.8 ± 1.3 | 79.3 |
|  | *reported*<sup>4</sup> | 91.9 | N/A |
| GPT-5 (Medium)** | **best of ours:** 1.0, 1.0, - | 85.7 ± 3.4 | 33.7 |
|  | *reported*<sup>5</sup> | 85.7 | N/A |
| GPT-4o | **best of ours:** 0.5, -, - | 52.0 ± 2.5 | 4.7 |
|  | *reported*<sup>6,7</sup> | ~ 49 (53.6 for full set) | N/A |

**Supplementary Table 15:** Serving configurations and performance on HLE Biology. We report mean accuracy (%) ± SD over three runs for overall, text-only, and vision subsets, plus average latency (sec/question). The decoding presets are defined above; “qwen_hybrid” uses “qwen_text” for text-only questions (222) and “qwen_vl” for vision questions (58). “best of ours” is the best reproducible setting in our setup (no official reported results are available for this split).

| Name | Config | Accuracy (%): overall; text; vision | Latency (sec) |
| --- | --- | --- | --- |
| Qwen3-VL-32B-Instruct | vllm_default | $7.7 \pm 0.9$; $5.9 \pm 1.2$; $14.9 \pm 1.0$ | 38.0 |
| | vllm_greedy | $6.4 \pm 1.2$; $4.4 \pm 1.4$; $14.4 \pm 1.0$ | 186.3 |
| | qwen_vl | $7.1 \pm 0.4$; $5.6 \pm 0.9$; $13.2 \pm 2.0$ | 38.9 |
| | qwen_text | $7.6 \pm 0.8$; $5.9 \pm 0.8$; $14.4 \pm 1.0$ | 46.7 |
| | qwen_hybrid* | $8.2 \pm 1.2$; $7.1 \pm 0.5$; $12.6 \pm 5.3$ | 38.8 |
| Ministral-3-14B-Instruct | vllm_default | $5.7 \pm 0.6$; $5.0 \pm 0.8$; $8.6 \pm 2.9$ | 5.8 |
| | vllm_greedy | $7.2 \pm 1.4$; $6.6 \pm 1.3$; $9.8 \pm 2.0$ | 17.0 |
| | production | $6.8 \pm 1.3$; $5.7 \pm 2.1$; $10.9 \pm 2.6$ | 21.5 |
| | creative | $6.6 \pm 1.2$; $5.1 \pm 1.0$; $12.1 \pm 1.8$ | 18.2 |
| Sonnet 4.5 | best of ours: 0.6 | $10.6 \pm 0.9$ | – |
| Gemini 3 Pro | best of ours: 0.2 | $31.4 \pm 1.3$ | – |
| GPT-5 (Medium) | best of ours: 1.0 | $18.1 \pm 1.8$ | – |
| GPT-4o | best of ours: 1.0 | $5.7 \pm 0.9$ | – |

###### Key observations and deployment trade-offs

On QA, *backend choice dominates sampling choice* for the API models (Extended Data Fig.1): the relative ranking is largely preserved across the sweep, while temperature mainly induces second-order shifts. Gemini 3 Pro is consistently strong and temperature-robust on GPQA, HLE Biology, and LabBench, with only small fluctuations around its optimum; in practice, many temperatures lie on an accuracy plateau and can be selected based on downstream considerations (*e.g.*, output diversity) rather than accuracy alone. Sonnet 4.5 exhibits a similarly broad plateau once *T* is modest, indicating low sensitivity to decoding noise in this setting. GPT-4o shows a clearer temperature dependence, including a mild mid-range optimum on GPQA, but remains substantially below the strongest backends across benchmarks, so tuning temperature does not compensate for backend capability. GPT-5 follows a different control axis: increasing `reasoning_effort` yields monotonic accuracy improvements across QA tasks, consistent with additional sampling-time compute improving deliberation, but the gains are accompanied by substantial increases in cost and latency, creating a practical operating point at Medium for most runs.

Open-weight models show a complementary pattern (Supplementary Tables 14 and 15): absolute accuracy is lower than the leading APIs under the same evaluation protocol, and decoding presets that adjust `top_p`/`top_k` can affect both reproducibility and runtime. On GPQA, Qwen3-VL-32B-Instruct is consistently the strongest open-weight checkpoint across the evaluated presets, while Ministral-3-14B is lower; however, multiple Qwen3 presets can attain comparable mean accuracy with markedly different variance, indicating that “best” decoding should be defined by both mean performance and stability. On HLE Biology, accuracies are low for open-weight models, but Qwen3 benefits from modality-aware decoding: the hybrid preset improves overall performance by applying text-oriented parameters to text-only questions and vision-oriented parameters to image questions, supporting the use of simple routing when a benchmark mixes modalities. Across both benchmarks, greedy decoding is generally a poor trade-off for open-weight serving, frequently increasing latency without improving accuracy, consistent with occasional overly long generations dominating wall-clock time.

Finally, run-to-run variation at low sampling settings reinforces that *T*=0 should not be treated as strictly deterministic for hosted APIs, and that reproducibility should be quantified directly. For subsequent experiments, we therefore (i) use mid-range temperatures for temperature-exposed APIs and GPT-5 Medium `reasoning_effort` as the default cost–accuracy compromise, and (ii) fix open-weight decoding to the most reproducible presets (Qwen3 “vLLM default”; Ministral “production”) unless otherwise stated.

##### 2.2.2. Gene Panel Design Without Skills

We next evaluate sampling sensitivity on the gene panel design task with the skill library disabled, so that any differences arise from the LLM backend and decoding alone in an end-to-end tool-using workflow. For API backends, we sweep sampling (temperature, or `reasoning_effort` for GPT-5) and report accuracy, (macro-)F1, spatial reconstruction *R*^2^, and wall-clock time (Supplementary Figs. 5–6).

**Supplementary Figure 5:**
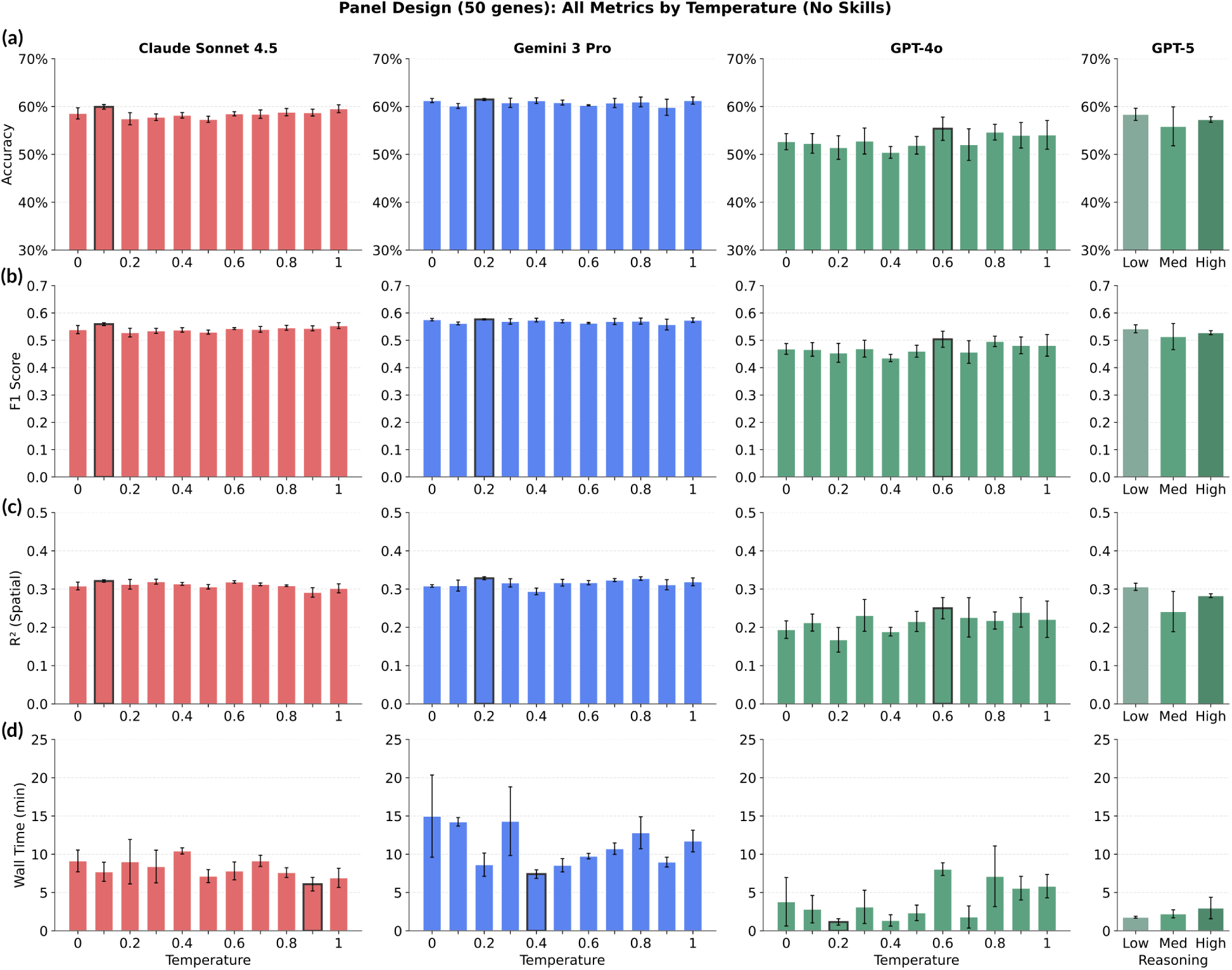
Sampling sensitivity for 50-gene panel design (no skills). Mean performance (y axis, a-c), or wall time (y axis, d) across temperature or `reasoning_effort` (GPT-5) sweeps (x axis) for each backend (columns). Error bars: s.d. over three independent runs; outlined bar: best setting in each subplot.

**Supplementary Figure 6:**
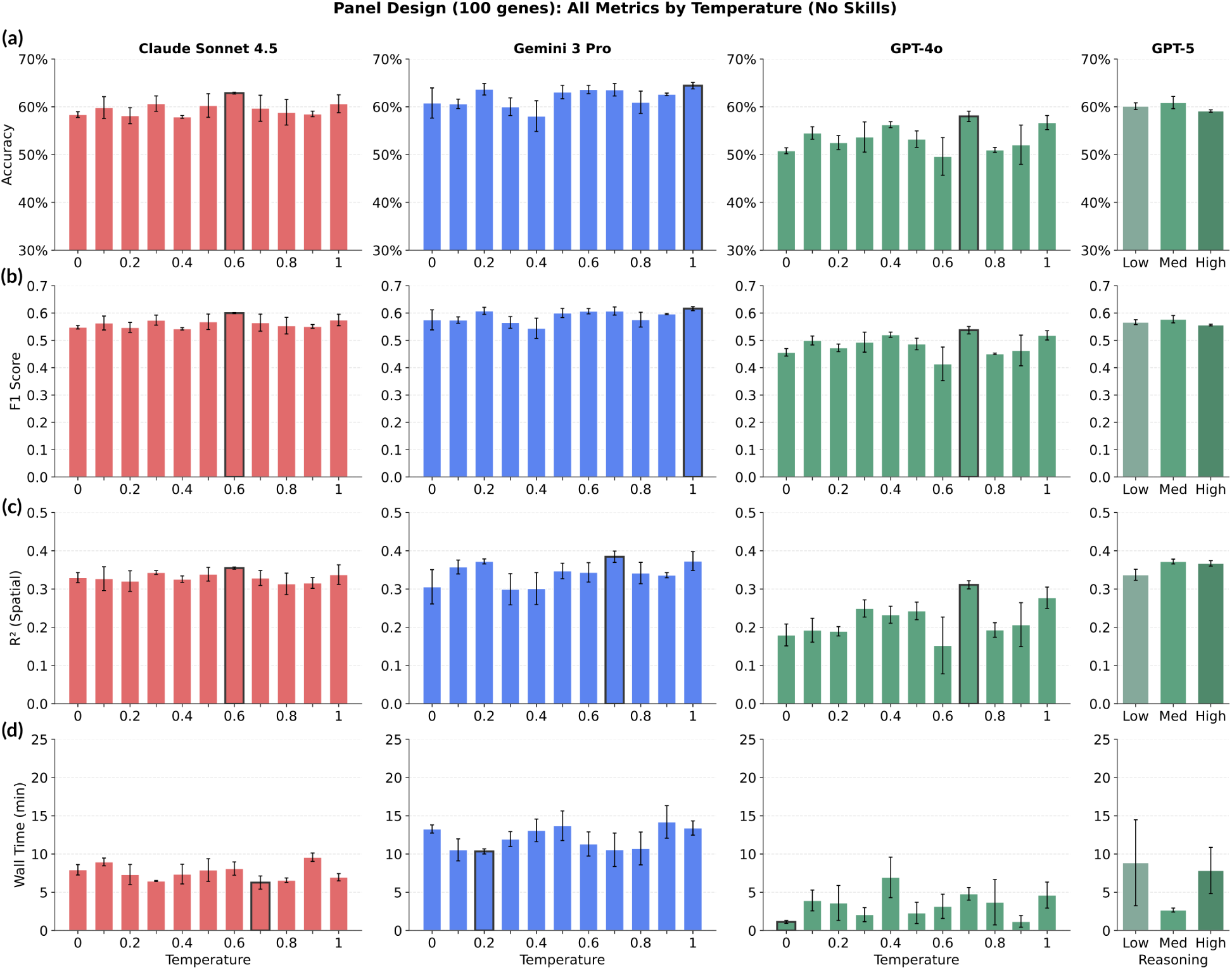
Sampling sensitivity for 100-gene panel design (no skills). Same as Fig. 5 for *N*=100 genes.

Across both *N*=50 and *N*=100 (Supplementary Figs. 5–6), accuracy and F1 are comparatively insensitive to temperature for the stronger API backends, with broad plateaus and largely preserved model ranking. In contrast, spatial *R*^2^ shows larger relative fluctuations and, for weaker backends, clearer sensitivity to decoding, suggesting that stochasticity affects fine-grained spatial fidelity more than coarse label agreement in this workflow. GPT-5 exhibits the expected cost–quality behavior: higher `reasoning_effort` can improve task metrics but should be interpreted jointly with runtime and constraint adherence.

**Supplementary Table 16:** Panel-size constraint adherence under sampling sweeps (no skills). Min–max (mean) number of genes produced per run for 50/100 gene panels across temperature sweeps.

|  | 50 Gene | 100 Gene |
| --- | --- | --- |
| Sonnet 4.5 | 50–50 (50.00) | 100–100 (100.00) |
| Gemini 3 Pro | 49–50 (49.97) | 100–100 (100.00) |
| GPT-4o | 4–50 (14.52) | 2–100 (14.74) |
| GPT-5 (low) | 48–50 (49.33) | 83–100 (89.33) |
| GPT-5 (medium) | 50–50 (50.00) | 83–96 (91.00) |
| GPT-5 (high) | 39–50 (44.33) | 54–73 (62.67) |

Runtime and downstream scores should be interpreted alongside the hard gene-count constraint (Table 16), since under-generation can both reduce wall time and limit the evaluator’s achievable score. Sonnet 4.5 and Gemini 3 Pro satisfy the requested panel size tightly across the sweep, whereas GPT-4o exhibits severe under-generation in no-skills mode. GPT-5 also under-generates in some settings, and higher `reasoning_effort` does not consistently improve constraint satisfaction, highlighting that more internal deliberation does not necessarily translate to better compliance in tool-using loops. Finally, some end-to-end failures arise from provider-side instability (*e.g.*, transient HTTP errors) rather than sampling; we therefore report completion separately from task metrics when identifiable.

##### 2.2.3. Cell-Type Annotation Without Skills

We evaluate sampling sensitivity for cell-type annotation with the skill library disabled, which exposes how decoding choices affect label selection and taxonomy consistency in an end-to-end workflow. We sweep temperature for Sonnet 4.5, Gemini 3 Pro, and GPT-4o, and sweep `reasoning_effort` (Low/Medium/High) for GPT-5. We report accuracy-oriented metrics (accuracy, balanced accuracy, macro-F1), structure-aware agreement (ARI), macro-recall, and coverage (number of annotated types) (Supplementary Fig. 7).

**Supplementary Figure 7:**
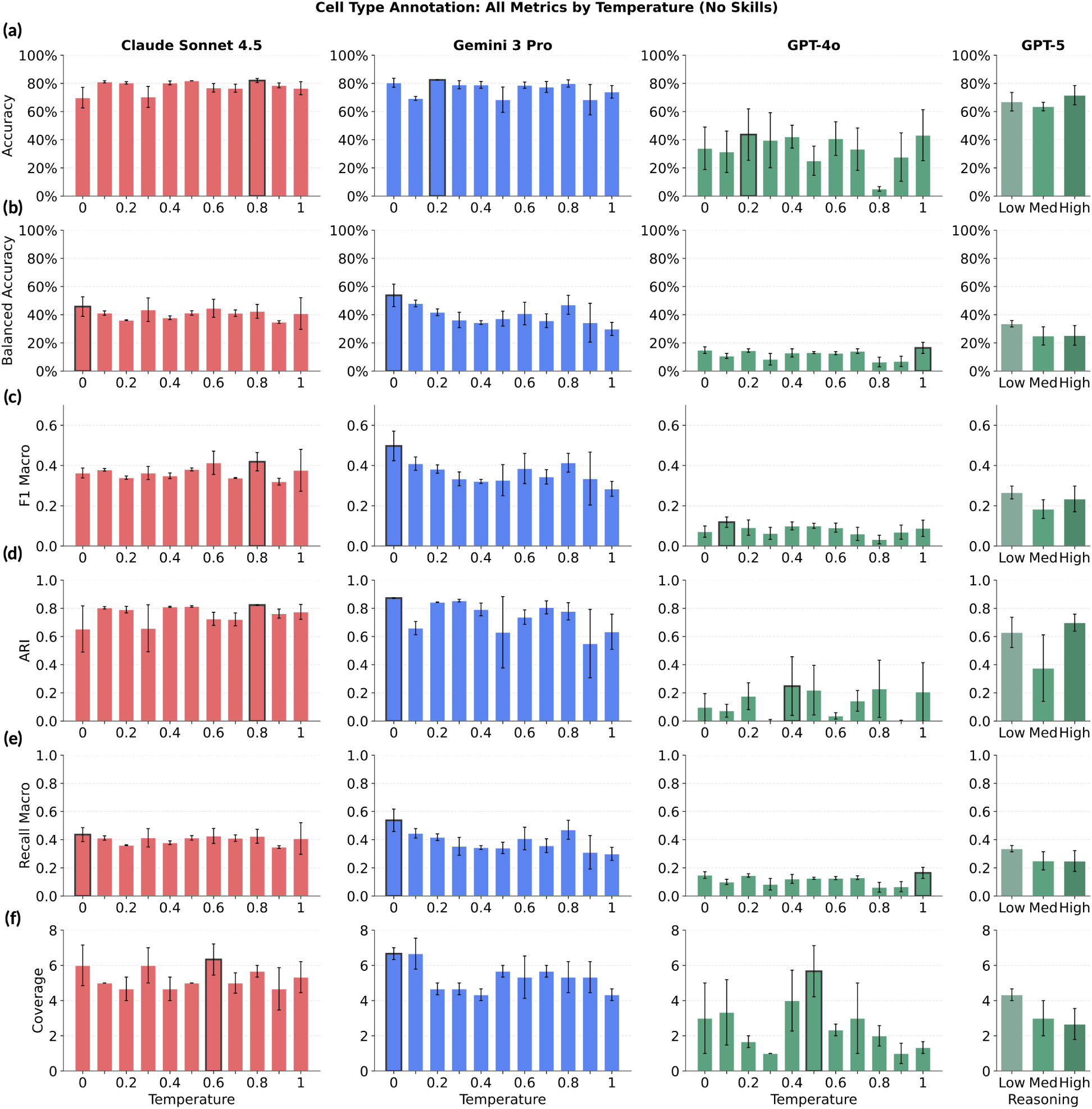
Sampling sensitivity for cell-type annotation (no skills). Mean performance metrics (y axis) for each backend (columns) across temperature sweeps (x axis) or reasoning effort (GPT-5, x axis) on cell type annotation. Error bars: SD over three independent runs. Outlined bar: best setting within each subplot.

Mean accuracy is often comparatively stable across sampling settings for the stronger backends, whereas balanced accuracy, macro-F1, and agreement metrics show larger shifts and increased variance at higher temperatures (Supplementary Fig. 7). This pattern is consistent with “terminological drift”: higher stochasticity increases the chance of producing over-specific labels, synonyms, or mixed-granularity outputs that deviate from the evaluation taxonomy. Such deviations can leave overall accuracy relatively unchanged (especially when dominant classes are easy), while reducing class-balanced metrics and cluster agreement due to inconsistent mapping across rare cell types.

Coverage further clarifies this effect. Higher temperatures can increase the number of distinct types produced, but this does not necessarily reflect better biological resolution; it can also arise from fragmented or inconsistent naming that lowers ARI and macro-F1. In practice, low-to-moderate temperatures provide the best reproducibility for Sonnet 4.5 and Gemini 3 Pro, yielding tighter error bars across metrics. GPT-4o remains substantially weaker and high-variance across the sweep, indicating that decoding changes do not reliably recover competitive annotation quality for this backend.

For GPT-5, increasing `reasoning_effort` changes the trade-off differently than changing temperature for other models: agreement metrics can improve at intermediate settings, but variance can also increase, suggesting that additional deliberation does not uniformly stabilize taxonomy alignment in the absence of explicit normalization.

Finally, completion differs substantially across backends in this no-skills setting; for large-scale annotation, this operational reliability can dominate small metric differences and motivates the use of skill templates and explicit normalization steps in subsequent sections.

##### 2.2.4. Summary

Across QA and both no-skills workflows, sampling primarily modulates variance, constraint adherence, and operational robustness, while mean task metrics for strong backends are often relatively flat across a broad range. Low-to-moderate temperatures generally provide the best reproducibility for Sonnet 4.5 and Gemini 3 Pro, GPT-5 performs well at `reasoning_effort=Medium`, and open-weight models benefit from fixed decoding presets chosen for stability rather than task-specific peak scores.

In closed-loop workflows, small differences in mean score can be outweighed by completion, constraint satisfaction, and taxonomy consistency. We therefore report mean ± s.d. over repeated runs throughout, interpret sampling sweeps jointly with robustness measures, and use these trends together with the backend and skill ablations to motivate the default configuration in Section 2.4.

#### 2.3. Skill ablations

We next isolate the contribution of *skills*—explicit workflow templates that decompose the task into ordered tool calls with intermediate checks and constrained outputs. Here, we ask whether workflow structure can recover reliability that decoding alone does not provide. Skills do not change the underlying tools or datasets; instead, they act as a control layer that (i) enforces task-specific structure (*e.g.*, required fields, valid label space), (ii) inserts lightweight validation and repair steps, and (iii) reduces failure modes from missed tool calls, formatting drift, and premature termination. To avoid over-interpreting a single decoding choice, we summarize *mean performance* across the sampling sweep within each backend (or the fixed decoding preset for open-weight models), comparing runs *without* skills to runs *with* skills under the same evaluation.

##### 2.3.1. Gene panel design

On panel design, skills yield backend-dependent gains (Fig. 8). For the strongest API backends, the mean metrics are already stable under no-skills execution, so skills provide only modest additional improvements. In contrast, the open-weight backends benefit more consistently: Qwen3-VL-32B shows clear mean gains across accuracy, F1, and spatial *R*^2^, indicating that an explicit scaffold helps it maintain the intended multi-step structure (produce a valid panel, propagate it through evaluation, and avoid partial outputs). Ministral shows smaller but generally positive shifts, consistent with the same mechanism but with a weaker backbone. Notably, skills do not uniformly improve every backend: for GPT-4o, the mean spatial *R*^2^ can decrease under skills, suggesting that the additional structure cannot compensate for core limitations in this workflow and may shift the model toward safer but less informative generations (*e.g.*, more conservative panels that satisfy constraints but provide weaker spatial fidelity). Overall, panel design highlights a key role of skills as a *stabilizer* for weaker or locally served backends, while the best-performing APIs are primarily limited by backbone capability rather than by missing workflow structure.

**Supplementary Figure 8:**
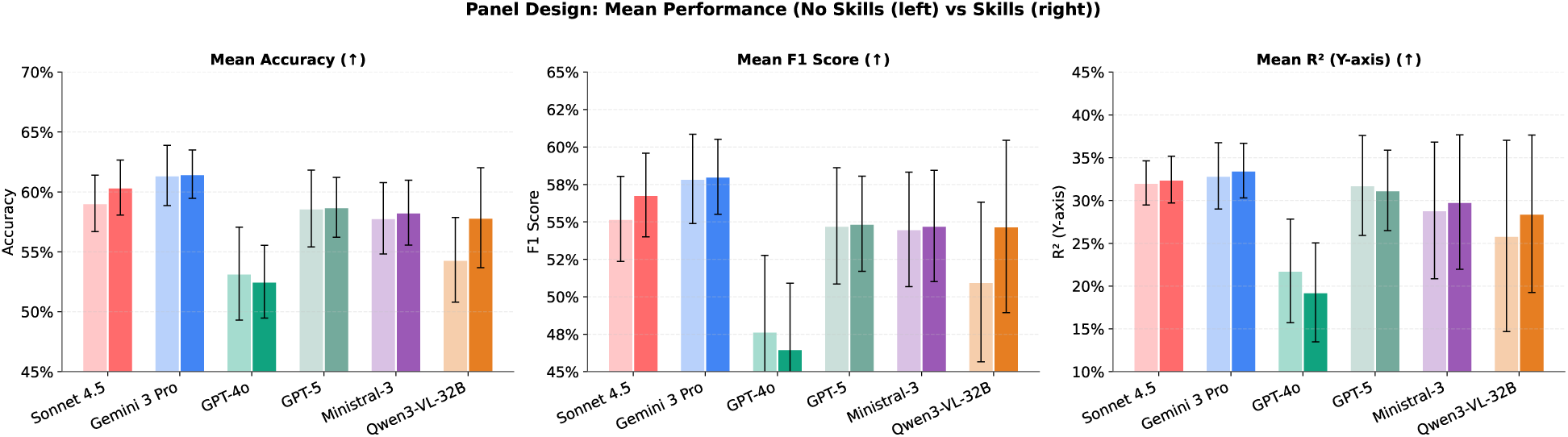
Skill ablation on gene panel design. Mean performance (y axis) on a panel design task across sampling settings for each backend (x axis) in no-skills (left bar) versus skills (right bar) settings. Error bars: standard deviation across three independent runs.

**Supplementary Figure 9:**
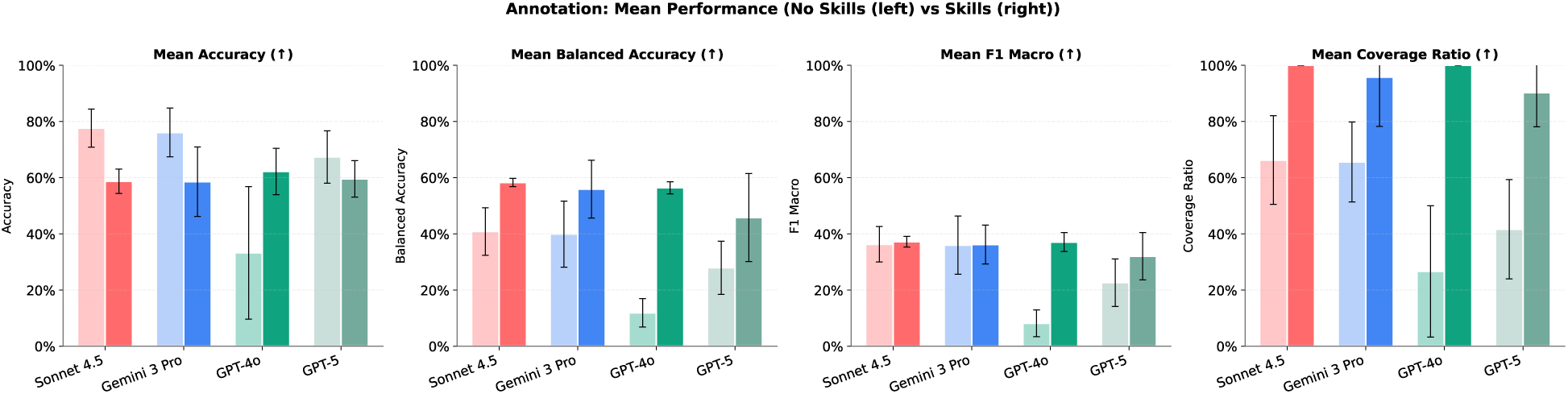
Skill ablation on cell-type annotation. Mean performance (y axis) on a cell annotation task across sampling settings for each backend (x axis) in no-skills (left bar) versus skills (right bar) settings. Error bars: standard deviation across three independent runs

##### 2.3.2. Cell-type annotation

For annotation, skills have a qualitatively different effect than in panel design: they primarily improve *taxonomy consistency and coverage*, which is more visible in balanced accuracy, macro-F1, and coverage than in overall accuracy (Supplementary Fig. 9). In the no-skills setting, backends can achieve deceptively high overall accuracy by collapsing onto frequent classes, while under-performing on rare types (low balanced accuracy) and leaving many types unassigned (low coverage). Skills counteract this failure mode by enforcing a structured labeling procedure and normalization steps that reduce terminological drift and encourage complete, taxonomy-aligned outputs. This shifts performance toward better class balance and broader coverage across backends.

The effect is most pronounced for weaker backends: GPT-4o shows large gains across accuracy, balanced accuracy, macro-F1, and coverage once skills are enabled, consistent with skills preventing common failure modes such as incomplete label sets, inconsistent granularity, or non-taxonomic responses. For stronger backends (Sonnet 4.5, Gemini 3 Pro, GPT-5), skills substantially increase coverage and improve balanced accuracy, while overall accuracy can decrease because the model no longer defaults to majority-class predictions and instead attempts a more fine-grained, taxonomically complete labeling. In practice, we treat this as a desirable trade-off for annotation-style deployments, where the goal is not only to maximize top-line accuracy, but also to maintain consistent coverage and reliable mapping across the full label space.

Across both tasks, skills act less like an “accuracy booster” and more like a *robustness layer*: they reduce workflow brittleness, improve constraint satisfaction and coverage, and shift performance toward metrics that reward faithful, complete structure. This benefit is largest for open-weight and weaker API backends, motivating the use of skills in the remaining experiments where end-to-end reliability is critical.

Taken together with the preceding subsections, these results suggest a simple decision rule: select a capable backend first, place it in a stable decoding regime second, and use skills to improve reliability and structural correctness, especially for weaker or locally served models.

#### 2.4. Default model selection: Claude Sonnet 4.5, temperature *T*=0.5

The analyses of Sections 2.1–2.3 converged on Claude Sonnet 4.5 at *T*=0.5 as the default configuration used in the main experiments. This choice balances strong task performance with stable closed-loop execution, good constraint adherence, and relatively low sensitivity to moderate changes in decoding.

Three considerations motivate this default. First, Sonnet 4.5 exhibits a broad performance plateau on QA benchmarks (Extended Data Fig.1), so moderate temperatures do not materially reduce accuracy relative to the best observed setting. Second, on the no-skills agentic workflows, Sonnet 4.5 shows stable end-to-end behavior, including strong adherence to hard constraints in panel design (Table 16) and comparatively low variance in annotation metrics at low-to-moderate temperatures (Supplementary Fig. 7). Third, once skills are enabled, stronger backends are primarily distinguished by backbone capability and robustness rather than by aggressive decoding, making *T*=0.5 a pragmatic operating point that preserves flexibility for intermediate decisions without over-specializing to near-greedy decoding.

Unless otherwise stated, we therefore use:

- **Claude Sonnet 4.5 (default).** *T*=0.5 for the main agentic experiments.
- **Gemini 3 Pro.** A low-to-moderate temperature within its stable regime when used as an API comparator.
- **GPT-5.** `reasoning_effort=Medium` as the default trade-off between accuracy and efficiency.
- **Open-weight backends.** Fixed decoding presets selected for reproducibility and stable downstream evaluation.

### 3. Benchmark and human-study protocols

This section consolidates the benchmark definitions, human-study design, evaluation criteria, and efficiency measurements used throughout the downstream analyses.

#### 3.1. Overview of benchmark tasks

We evaluate three categories of tasks: question-answering benchmarks (GPQA Diamond, HLE Biology, and LabBench), constrained gene-panel design, and spatial annotation workflows. For tool-using workflows, inputs are standardized before evaluation using the preprocessing conventions described in Section 1, including quality-control filtering, normalization, log transformation, and dimensionality reduction as implemented in `preprocess_spatial_data`. Unless a benchmark explicitly varies a factor, prompts, datasets, tools, and evaluation code are held fixed across runs. For structured outputs, predictions are harmonized to author-provided taxonomies or target formats before scoring so that performance differences reflect task behavior rather than formatting mismatches.

#### 3.2. Gene panel design

Automated gene-panel design is evaluated as a constrained generation task in which the agent must propose fixed-size panels and support downstream spatial reconstruction and discrimination. In the ablations reported here, we evaluate 50-gene and 100-gene settings and report accuracy, macro-F1, spatial reconstruction *R*^2^, panel-size adherence, completion rate, and wall-clock time. Because the requested panel size is a hard constraint, exact gene-count compliance is tracked separately from downstream performance. The human study additionally includes 150-gene panel design, allowing direct comparison between model-generated and human-designed panels at multiple target budgets.

#### 3.3. Cell-type and tissue-niche annotation

Cell-type annotation is evaluated as an end-to-end workflow that maps spatial profiles or clusters to a common cell-type taxonomy. We benchmark SpatialAgent against CellTypist and GPTCellType, harmonizing all predictions to author-provided ground truth before scoring. Reported readouts include accuracy, balanced accuracy, macro-F1, ARI, recall, coverage, and completion, which together separate overall performance from taxonomy consistency and rare-class behavior.

Tissue-niche annotation is evaluated under a common Tier-1 spatial taxonomy aligned with author-provided labels. Because this task depends on spatial organization as well as gene expression, we assess both quantitative agreement with ground truth and qualitative anatomical coherence in the resulting niche maps. Human and model outputs are visualized in the same spatial frame to facilitate side-by-side comparison of niche assignment patterns and recurrent failure modes.

#### 3.4. Human scientist comparison protocol

The study design is described in the main **Methods**. Briefly, experts with PhD or MD/PhD backgrounds in relevant fields completed two stages: (1) designing 50-, 100-, or 150-gene panels for the DLPFC tissue; and (2) rating gene panels generated by SpatialAgent, GPT-4o, other scientists, and hybrid methods using a 5-point rubric covering accuracy, completeness, reasoning, and conciseness. Researchers employed several distinct strategies when selecting genes for spatial transcriptomics panels (Extended Data Table. 1). Most researchers relied on either the provided reference dataset or literature sources, with only a few integrating multiple data sources. Their selection methods typically fell into three categories: (1) algorithmic approaches (Persist algorithm, iterative greedy methods, or highly variable gene selection), (2) differential expression analysis to identify cell type-specific markers, or (3) knowledge-driven selection of known markers from literature. The more comprehensive approaches, such as those from scientist 7, combined multiple strategies by cross-referencing markers from different sources and including genes related to specific biological processes or pathways of interest. Some researchers focused exclusively on cell type identification, while others extended their panels to capture pathway activities or disease-relevant genes.

For annotation tasks, researchers employed consistent approaches for cell and niche annotation in spatial transcriptomics data (Supplementary Table. 2). For cell annotation, most relied on computational clustering (primarily Leiden) followed by either manual annotation with marker genes or automated label transfer methods like CellTypist (22) or TACCO (44). Many provided hierarchical (multi-tier) annotations to capture both major cell types and subtypes. Only one researcher used a unique co-expression pattern approach instead of clustering. For niche annotation, UTAG spatial clustering was the predominant method, and differences emerged primarily in how researchers assigned biological meaning to these spatial clusters using spatial position, anatomical images, and prior anatomical knowledge.

Detailed instructions, allowed time, and study logistics are described in the main **Methods**. In brief, participants operated under a shared task definition and common benchmark materials, which makes differences in submitted outputs interpretable as differences in strategy rather than access to different resources. For gene panel design, participants submitted fixed-size candidate panels and subsequently rated anonymized outputs from humans, SpatialAgent, GPT-4o, and hybrid designs. For annotation, submitted cell-type and tissue-niche labels were harmonized to a common Tier-1 taxonomy before comparison to ground truth.

Besides the standard quantitative measures, we also asked the human scientists to rate anonymized gene panels from other scientists, SpatialAgent, and hybrid designs for accuracy, completeness, and reasoning, using a 1–5 scale (1 = Poor, 5 = Excellent) (Supplementary Fig.10). We additionally visualized the human annotation outputs in a common coordinate system and compared both cell-type and tissue-niche assignments to the ground truth, enabling direct inspection of agreement and boundary-case disagreements.

#### 3.5. Evaluation, runtime, and cost

We track both task quality and operational efficiency. For QA benchmarks, we report accuracy together with cost per GPQA run and latency per GPQA question when provider APIs expose the relevant accounting signals. For gene panel design, we report accuracy, macro-F1, spatial reconstruction *R*^2^, panel-size adherence, completion, and wall-clock time. For cell-type and tissue-niche annotation, we report agreement and coverage metrics after harmonizing outputs to the target taxonomies so that scoring reflects biological alignment rather than label-format variation.

SpatialAgent also tracks token usage and estimated cost through a callback mechanism that records input and output tokens for each LLM call and aggregates them over the session. Throughout the supplement, we interpret runtime and cost jointly with completion and constraint adherence, because failures such as under-generation, taxonomy drift, or provider-side instability can change both quality metrics and observed execution time.

We visualized the human annotations (Supplementary Fig. 11-16) in the same UMAP coordinates, where each scientist’s annotations represent their individual interpretations. For tissue niche annotations, we map them onto the spatial coordinates of each MERFISH spot. We further compared each scientist’s cell type annotations (Supplementary Fig.17–19) and tissue niche annotations (Supplementary Fig.19–20) to the ground truth.

**Supplementary Figure 10:**
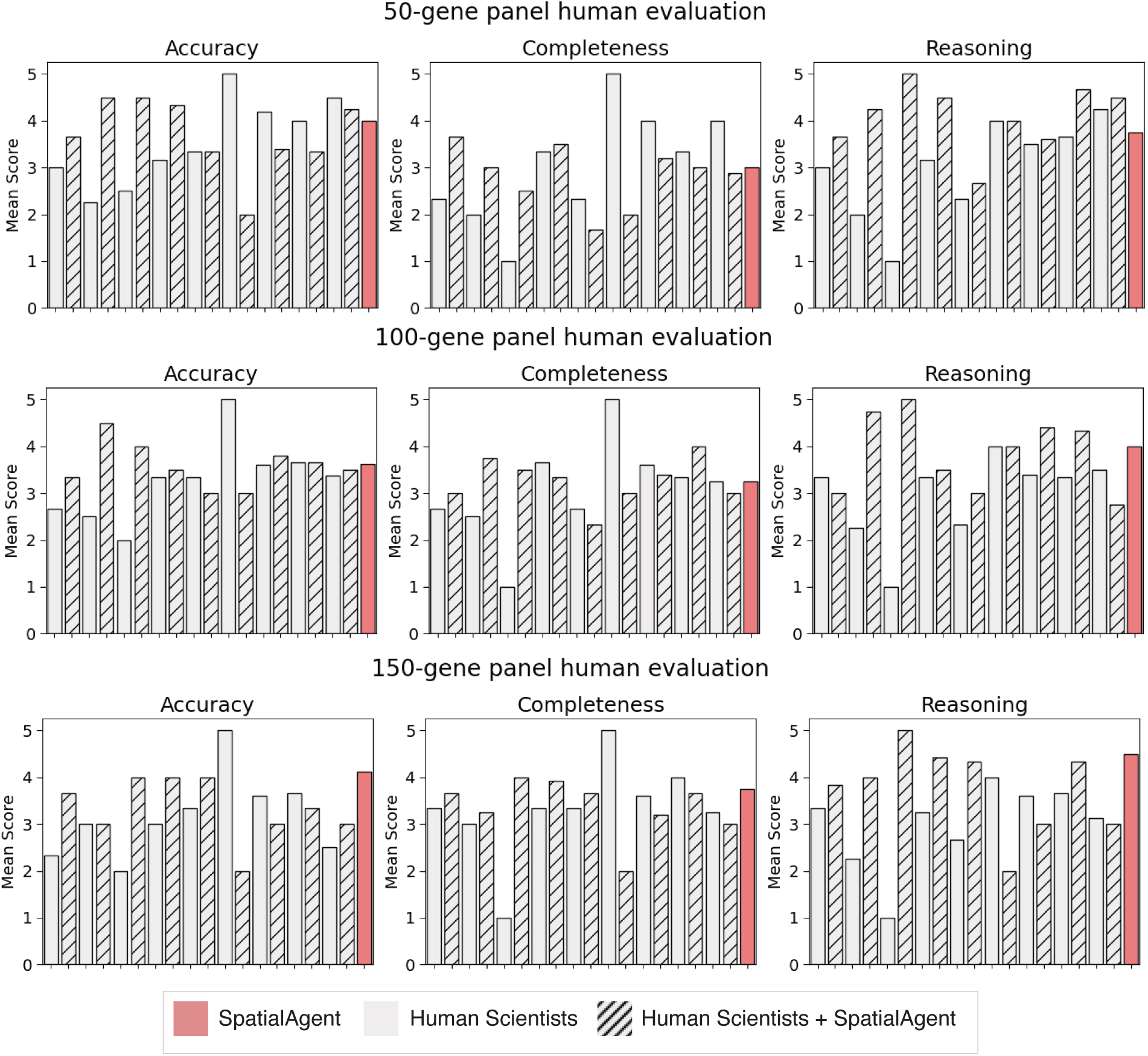
Human evaluation of panel design

**Supplementary Figure 11:**
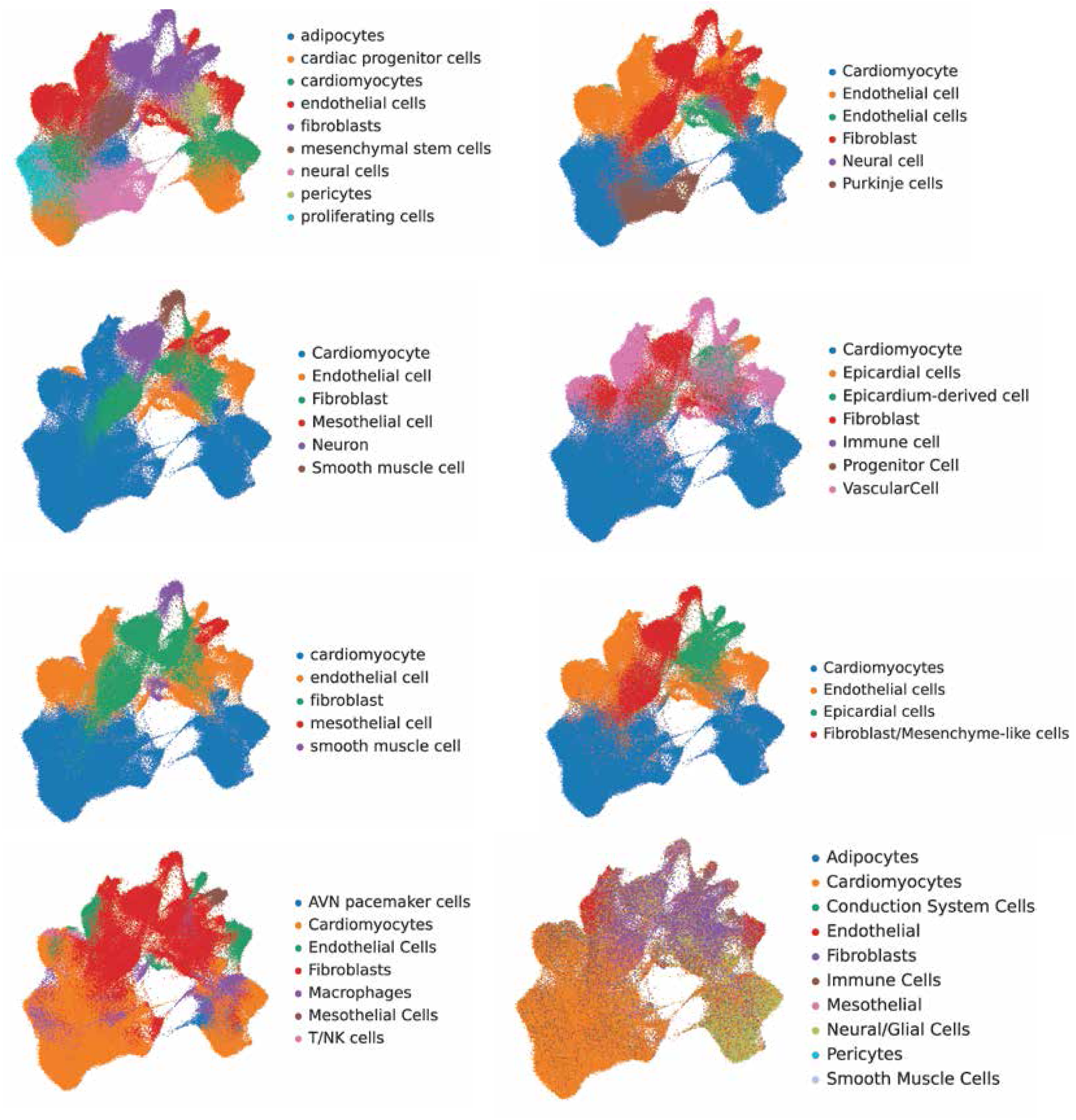
Original Tier 1 cell type annotations.

**Supplementary Figure 12:**
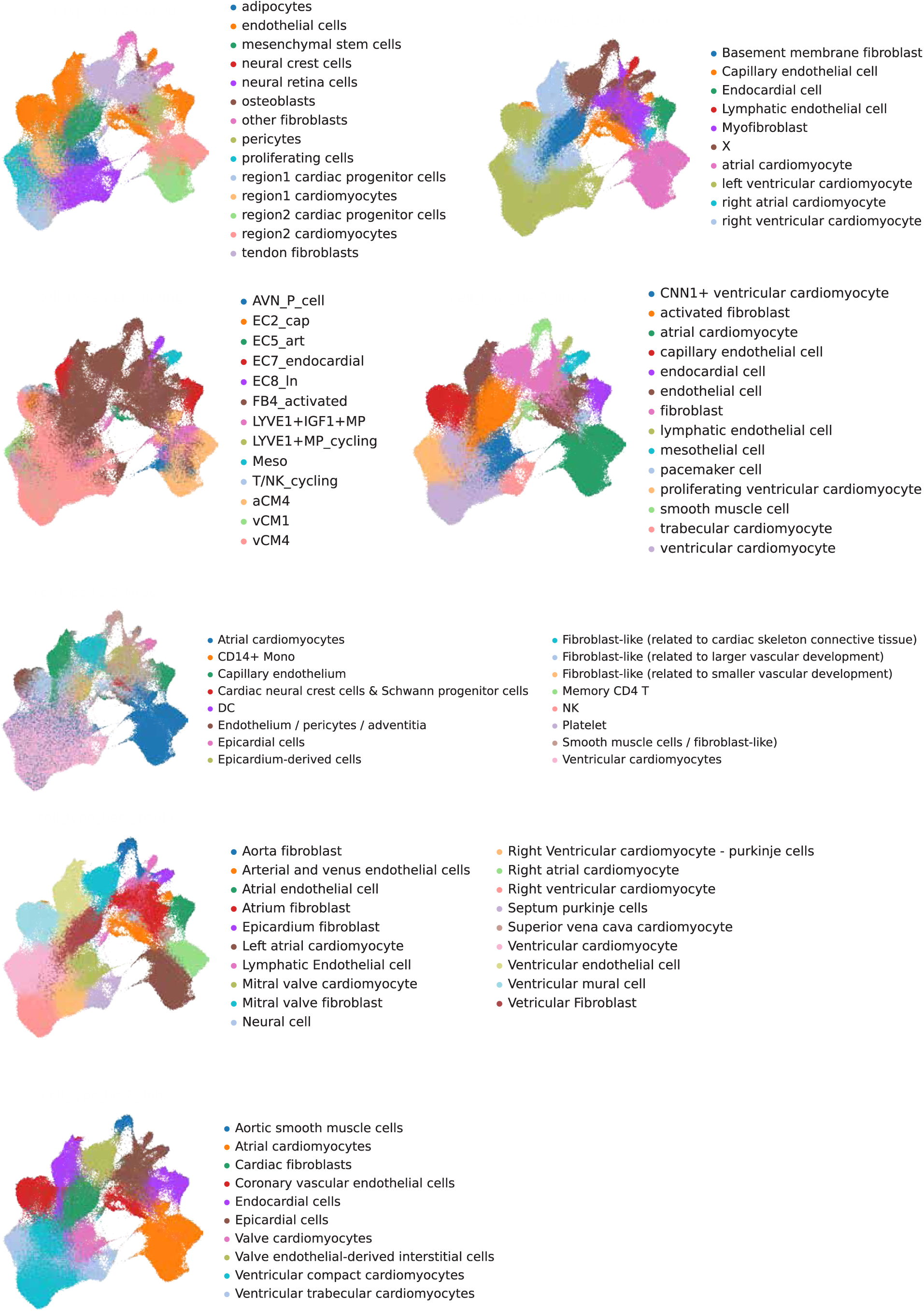
Original Tier 2 cell type annotations.

**Supplementary Figure 13:**
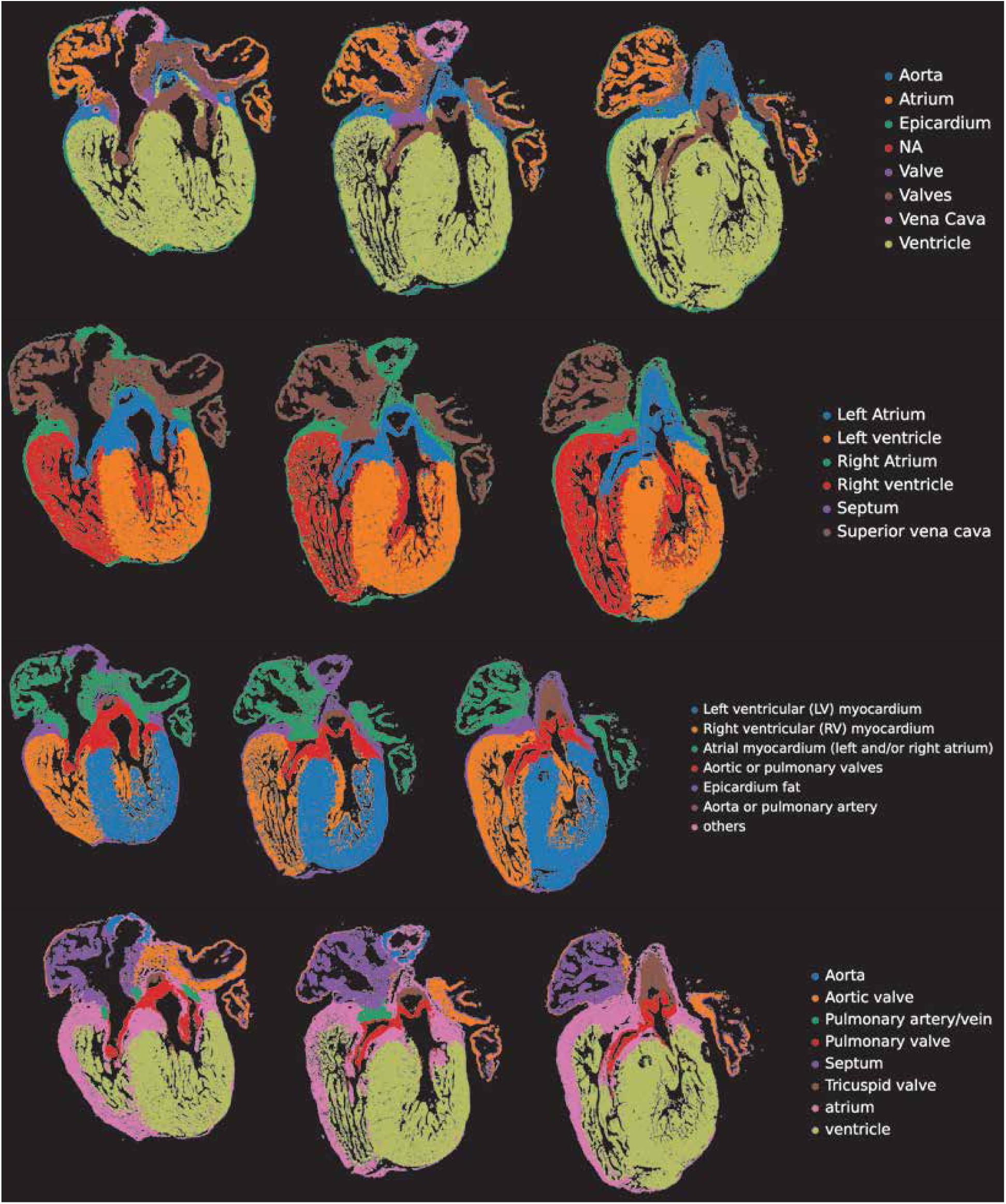
Original Tier 1 tissue niche annotations (1/2).

**Supplementary Figure 14:**
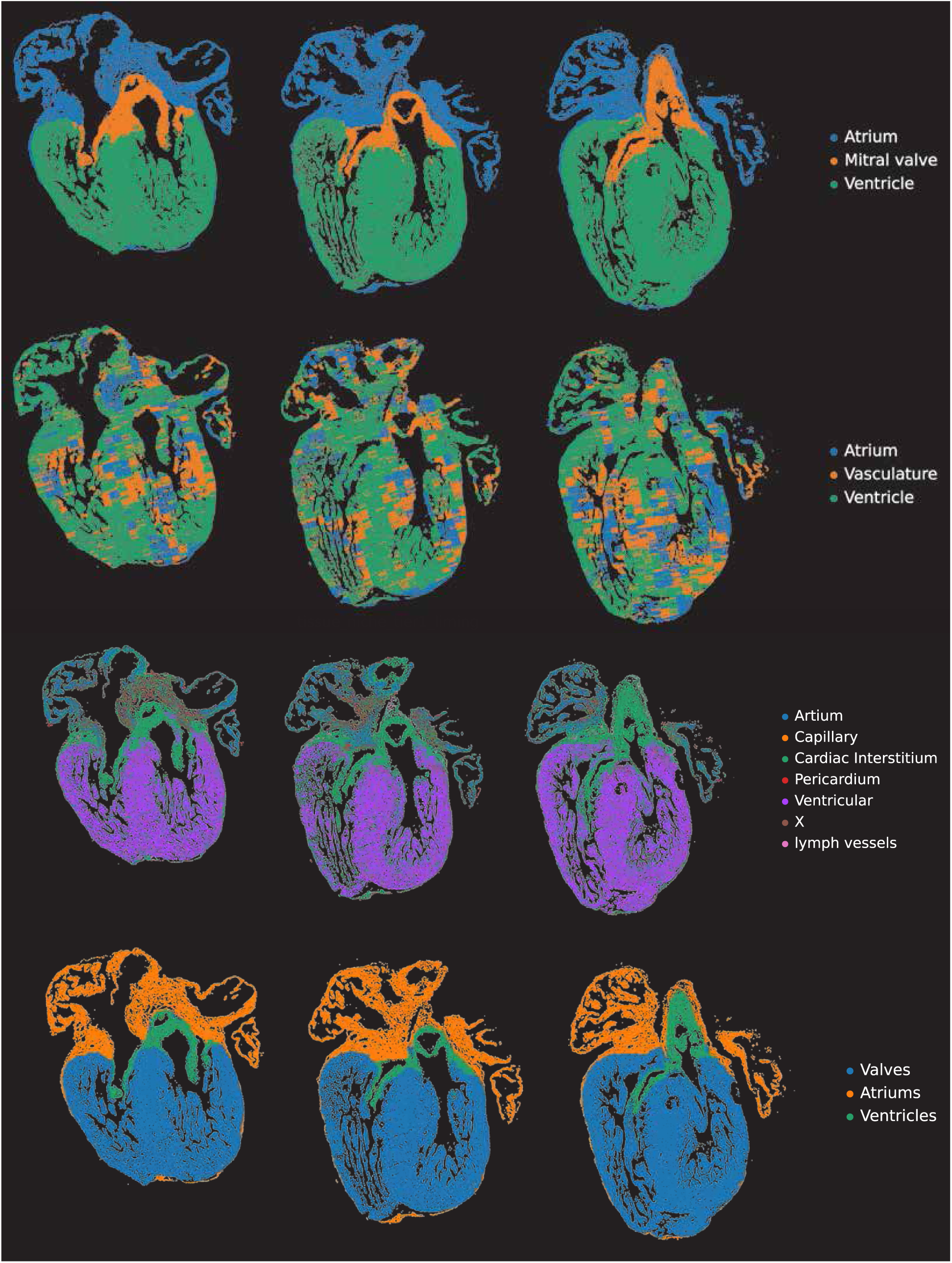
Original Tier 1 tissue niche annotations (2/2).

**Supplementary Figure 15:**
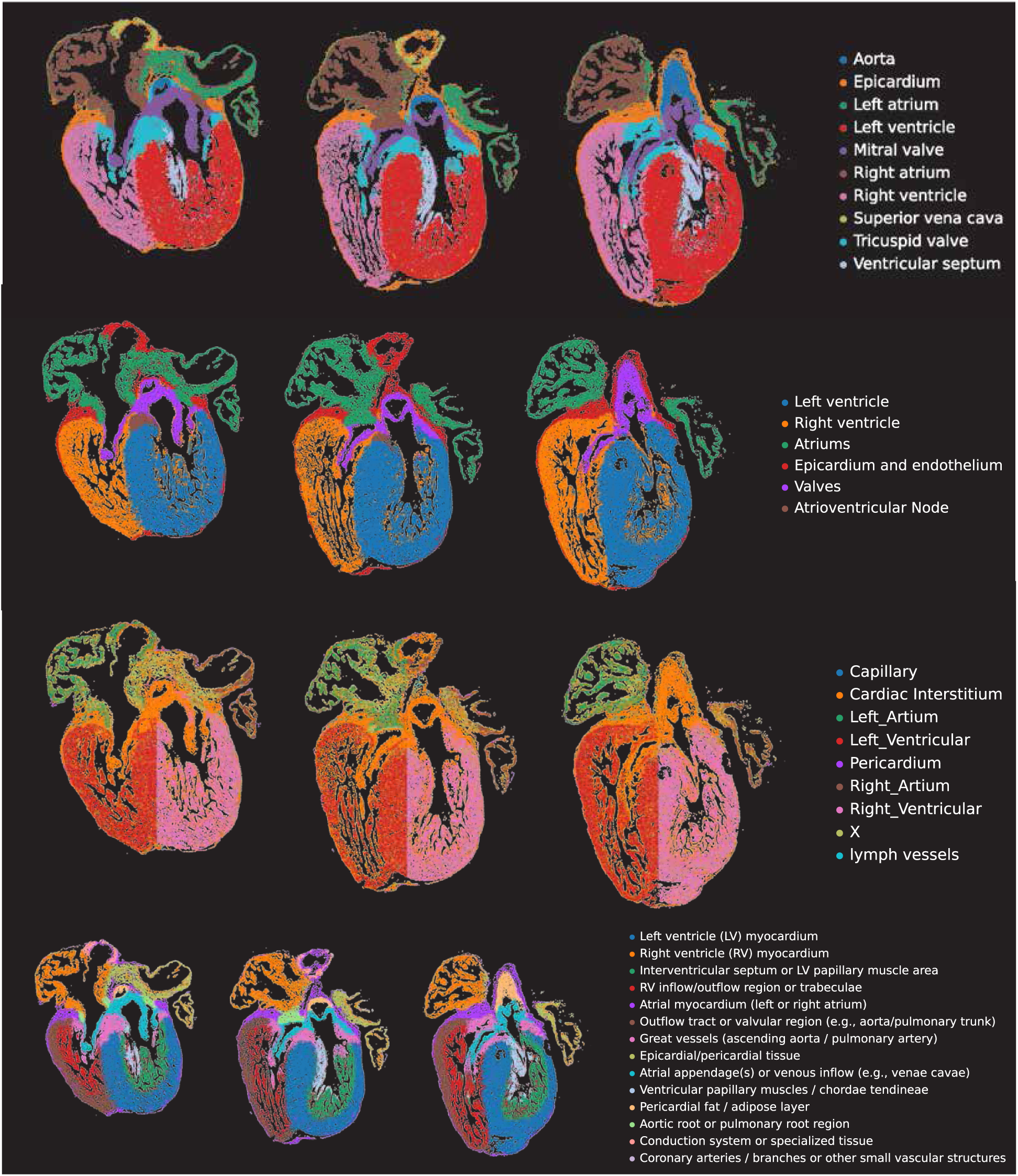
Original Tier 2 tissue niche annotations (1/2).

**Supplementary Figure 16:**
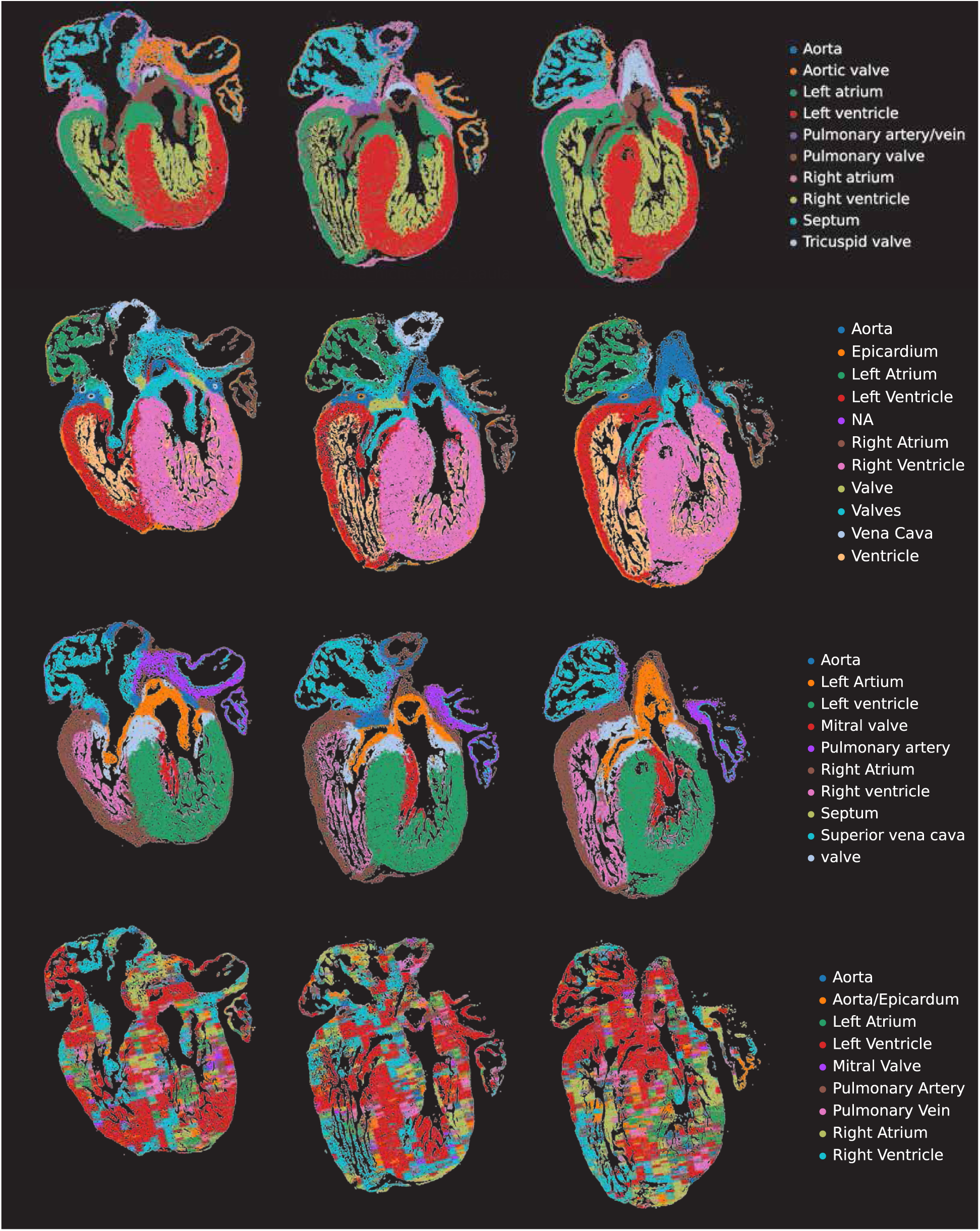
Original Tier 2 tissue niche annotations (2/2).

**Supplementary Figure 17:**
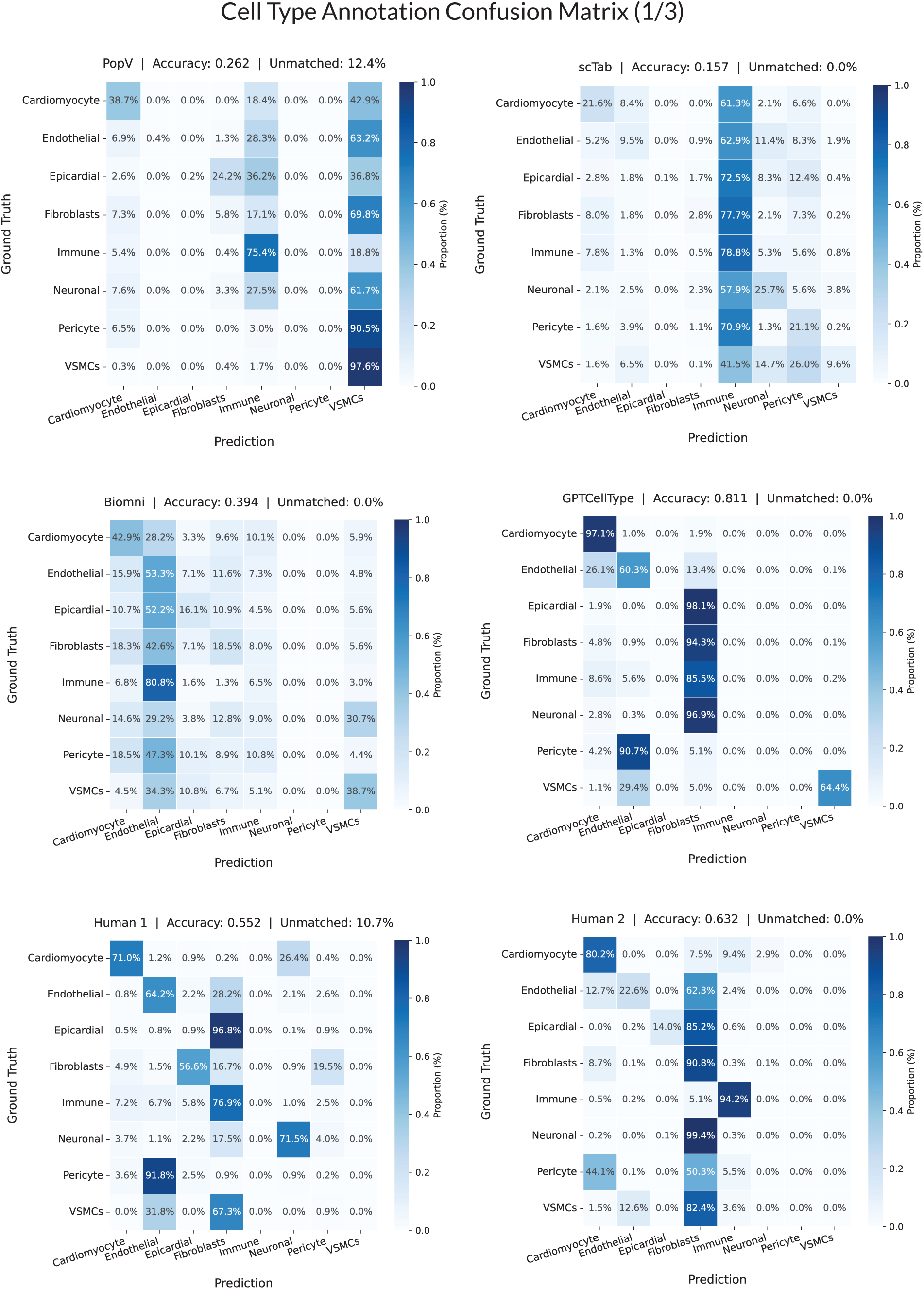
Confusion matrix of cell and tissue annotation (1/4).

**Supplementary Figure 18:**
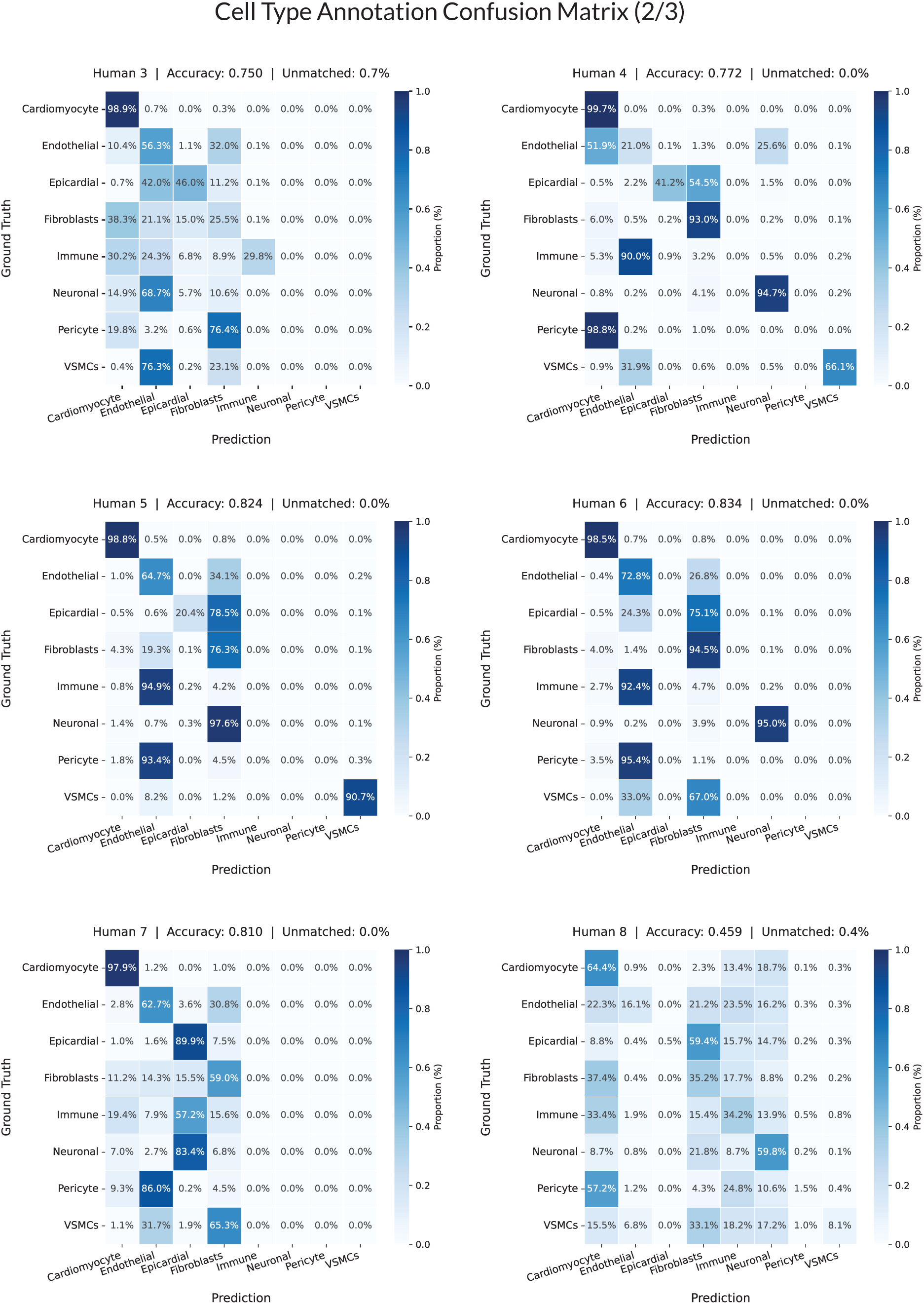
Confusion matrix of cell and tissue annotation (2/4).

**Supplementary Figure 19:**
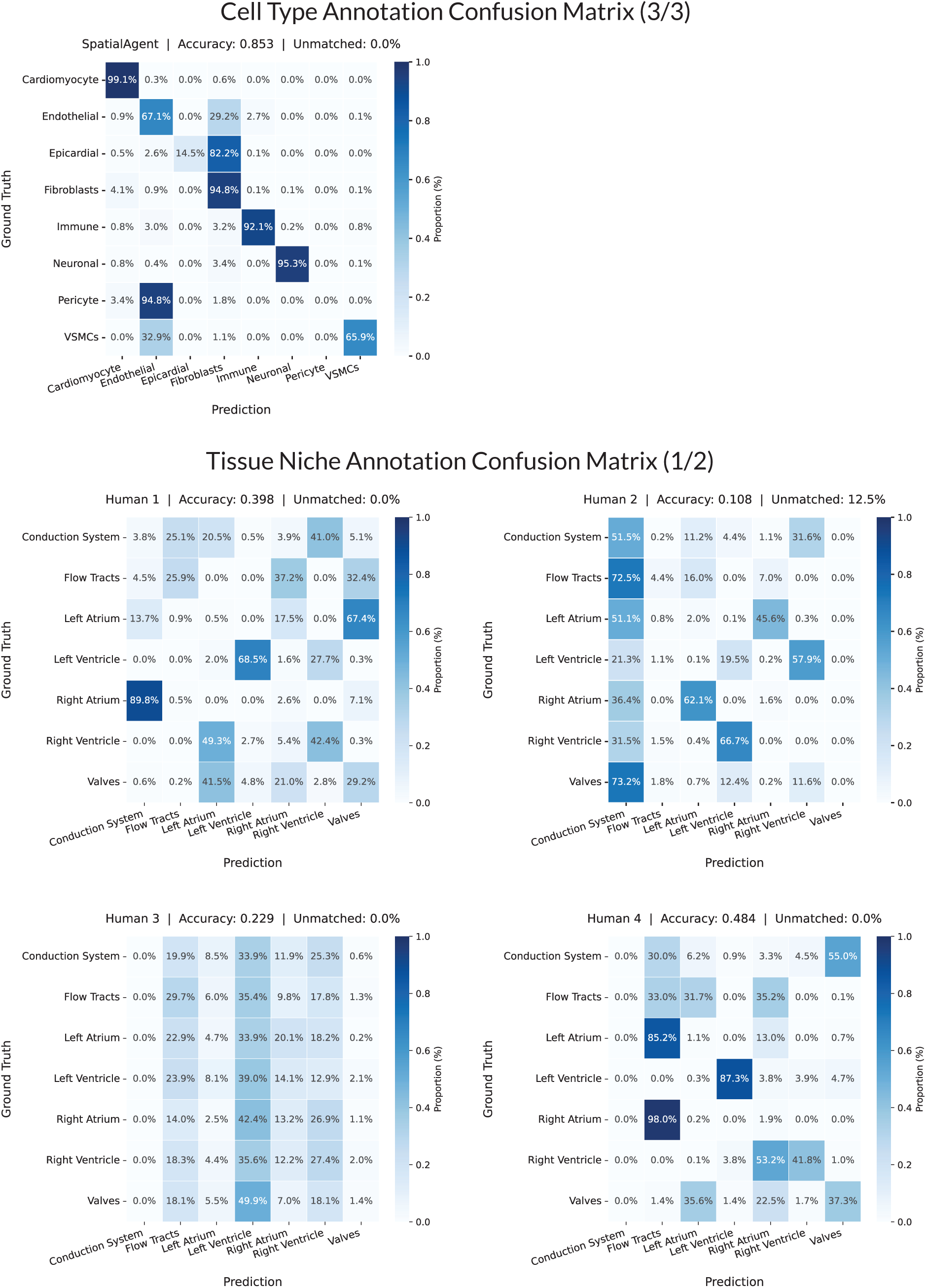
Confusion matrix of cell and tissue annotation (3/4).

**Supplementary Figure 20:**
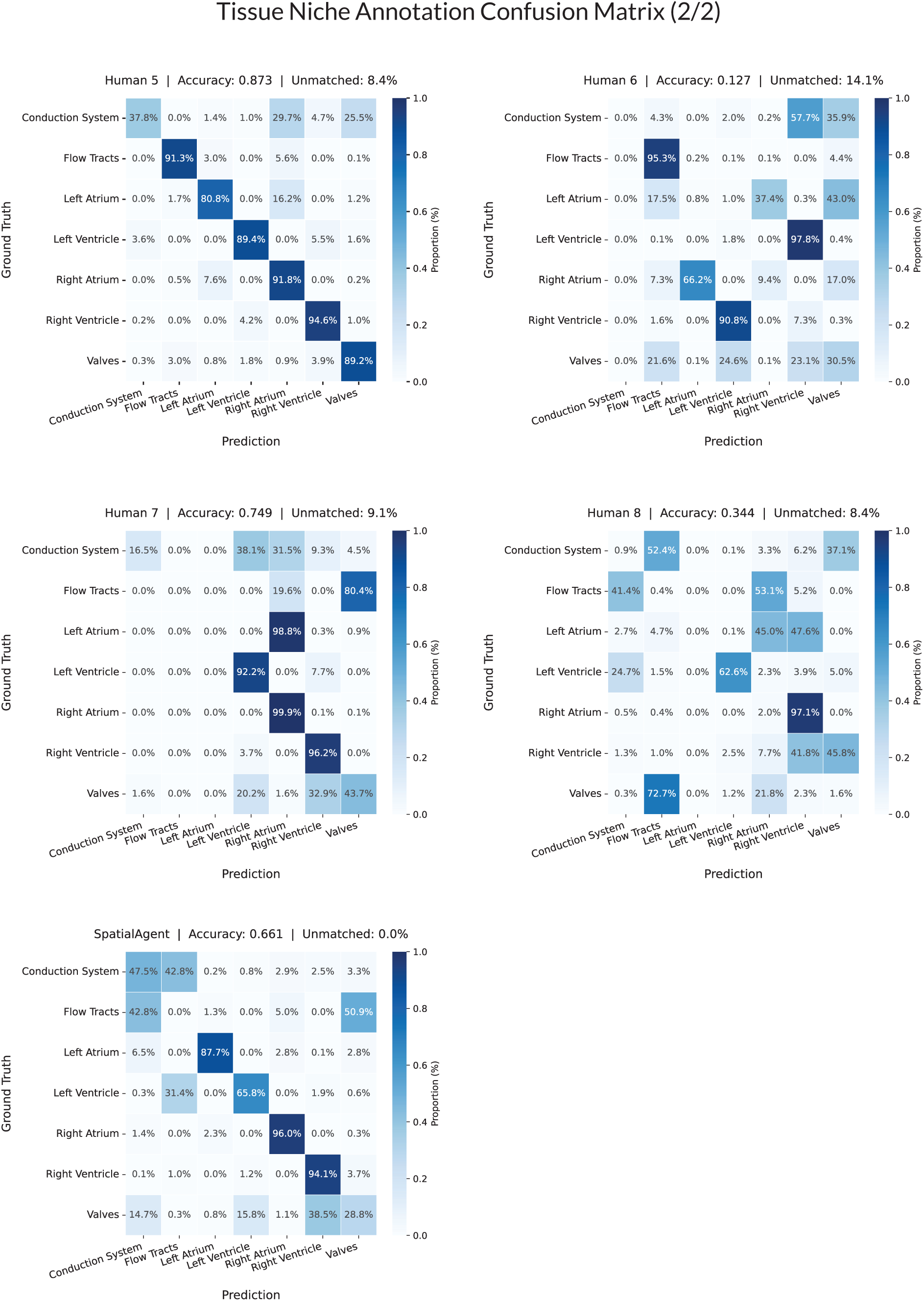
Confusion matrix of cell and tissue annotation (4/4).

### 4. Expanded case studies and wet-lab details

This section provides additional details on the analyses and wet-lab validation corresponding to the case studies discussed in the main text. Agent traces for all the case studies are available at https://github.com/Genentech/SpatialAgent/tree/case/notebooks.

#### 4.1. Neighborhood trajectory inference framework used in MERFISH mouse colitis dataset

All outputs shown are artifacts produced by SpatialAgent. In the MERFISH mouse colitis case study, SpatialAgent constructed a neighborhood trajectory analysis framework by anchoring spatial neighborhood dynamics on disease stage as ground truth. The agent first summarized each neighborhood by its normalized cell-type composition and quantified normalized neighborhood abundance across four disease stages: Healthy, DSS3, DSS9, and DSS21. It then calculated neighborhood abundance fold-changes between stages and inferred transition paths using three complementary criteria: temporal progression, cell-type composition similarity, and biological function. This framework identified seven neighborhood trajectories across the disease course, including mucosal trajectories (MU3→MU4→MU6→MU7→MU9 and MU1→MU2→MU8→MU10), a follicular trajectory (FOL2→FOL1→FOL3), a smooth muscle trajectory (SM1→SM2→SM3), a mesenchymal trajectory (ME1→ME3→ME4), and additional transitions linking MU5→MU8→LUM and FOL1→MU11.

**Supplementary Figure 21:**
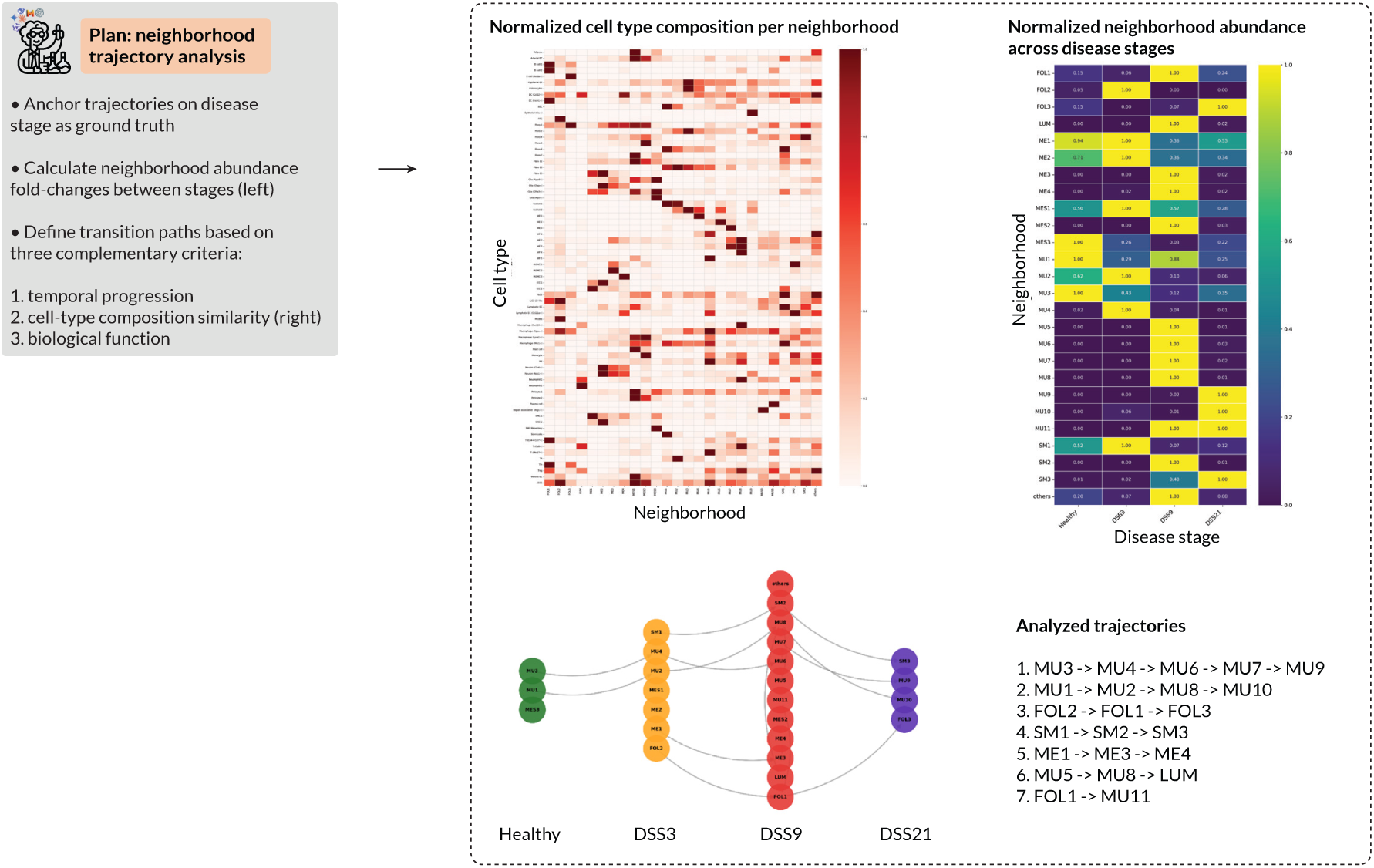
Neighborhood trajectory inference framework for the MERFISH mouse colitis case study. SpatialAgent autonomously planned a neighborhood trajectory analysis by anchoring trajectories on disease stage as ground truth. The framework integrates normalized cell-type composition per spatial neighborhood, normalized neighborhood abundance across disease stages, and abundance fold-changes between stages. Transition paths were defined using temporal progression, cell-type composition similarity, and biological function. This analysis identified seven trajectories spanning mucosal, follicular, smooth muscle, mesenchymal, luminal-associated, and follicle-to-mucosal neighborhood transitions across the disease course.

## References

[1] Hanchen Wang, Tianfan Fu, Yuanqi Du, et al. Scientific discovery in the age of artificial intelligence. Nature, 620(7972):47–60, 2023.

[2] Tom Brown et al. Language models are few-shot learners. In Advances in Neural Information Processing Systems, 2020.

[3] Shanghua Gao, Ada Fang, Yepeng Huang, et al. Empowering biomedical discovery with ai agents. Cell, 187(22):6125–6151, 2024.

[4] Giovanni Palla, David Fischer, Aviv Regev, et al. Spatial components of molecular tissue biology. Nature Biotechnology, 40(3):308–318, 2022.

[5] Juraj Gottweis, Wei-Hung Weng, Alexander Daryin, et al. Towards an ai co-scientist. arXiv:2502.18864, 2025.

[6] Quintina Campbell, Sam Cox, Jorge Medina, et al. Mdcrow: Automating molecular dynamics workflows with large language models. arXiv:2502.09565, 2025.

[7] Gao Shanghao, et al. Txagent: An ai agent for therapeutic reasoning across a universe of tools. arXiv:2503.10970, 2025.

[8] Yusuf Roohani, Andrew Lee, Qian Huang, Jian Vora, Zachary Steinhart, Kexin Huang, Alexander Marson, Percy Liang, and Jure Leskovec. Biodiscoveryagent: An ai agent for designing genetic perturbation experiments. arXiv preprint arXiv:2405.17631, 2024.

[9] Kexin Huang, Serena Zhang, Hanchen Wang, Yuanhao Qu, Yingzhou Lu, Ryan Li, Yusuf Roohani, Lin Qiu, Shiyi Cao, Gavin Li, et al. Autonomous biomedical research with an artificial intelligence agent. Science, page eadz4351, 2026.

[10] Zuwan Lin, Wenbo Wang, Arnau Marin-Llobet, Qiang Li, Samuel D Pollock, Xin Sui, Almir Aljovic, Jaeyong Lee, Jongmin Baek, Ningyue Liang, et al. Spatial transcriptomics ai agent charts hpsc-pancreas maturation in vivo. bioRxiv, pages 2025–04, 2025.

[11] Kyle Swanson, Wesley Wu, Nash L Bulaong, John E Pak, and James Zou. The virtual lab of ai agents designs new sars-cov-2 nanobodies. Nature, 646(8085):716–723, 2025.

[12] Weike Zhao, Chaoyi Wu, Yanjie Fan, Xiaoman Zhang, Pengcheng Qiu, Yuze Sun, Xiao Zhou, Yanfeng Wang, Xin Sun, Ya Zhang, et al. An agentic system for rare disease diagnosis with traceable reasoning. arXiv preprint arXiv:2506.20430, 2025.

[13] Zhizheng Wang, Qiao Jin, Chih-Hsuan Wei, Shubo Tian, Po-Ting Lai, Qingqing Zhu, Chi-Ping Day, Christina Ross, Robert Leaman, and Zhiyong Lu. Geneagent: self-verification language agent for gene-set analysis using domain databases. Nature Methods, 22(8):1677–1685, 2025.

[14] Ludovico Mitchener, Angela Yiu, Benjamin Chang, Mathieu Bourdenx, Tyler Nadolski, Arvis Sulovari, Eric C Landsness, Daniel L Barabasi, Siddharth Narayanan, Nicky Evans, et al. Kosmos: An ai scientist for autonomous discovery. arXiv preprint arXiv:2511.02824, 2025.

[15] Kristen R Maynard, Leonardo Collado-Torres, Lukas M Weber, et al. Transcriptome-scale spatial gene expression in the human dorsolateral prefrontal cortex. Nature neuroscience, 24(3):425–436, 2021.

[16] Tim Stuart, Andrew Butler, Paul Hoffman, et al. Comprehensive integration of single-cell data. cell, 177(7):1888–1902, 2019.

[17] Alsu Missarova, Jaison Jain, Andrew Butler, et al. genebasis: an iterative approach for unsupervised selection of targeted gene panels from scrna-seq. Genome biology, 22:1–22, 2021.

[18] Ian Covert, Rohan Gala, Tim Wang, et al. Predictive and robust gene selection for spatial transcriptomics. Nature Communications, 14(1):2091, 2023.

[19] Louis Kuemmerle, Malte Luecken, Alexandra Firsova, et al. Probe set selection for targeted spatial transcriptomics. Nature methods, pages 1–11, 2024.

[20] Elie N Farah, Robert K Hu, Colin Kern, et al. Spatially organized cellular communities form the developing human heart. Nature, 627(8005):854–864, 2024.

[21] Wenpin Hou and Zhicheng Ji. Assessing gpt-4 for cell type annotation in single-cell rna-seq analysis. Nature Methods, pages 1–4, 2024.

[22] C Domínguez Conde, Chao Xu, Louie B Jarvis, et al. Cross-tissue immune cell analysis reveals tissue-specific features in humans. Science, 376(6594):eabl5197, 2022.

[23] Can Ergen, Galen Xing, Chenling Xu, Martin Kim, Michael Jayasuriya, Erin McGeever, Angela Oliveira Pisco, Aaron Streets, and Nir Yosef. Consensus prediction of cell type labels in single-cell data with popv. Nature Genetics, 56(12):2731–2738, 2024.

[24] Felix Fischer, David S Fischer, Roman Mukhin, Andrey Isaev, Evan Biederstedt, Alexandra-Chloé Villani, and Fabian J Theis. sctab: Scaling cross-tissue single-cell annotation models. Nature Communications, 15(1):6611, 2024.

[25] Bob Chen, Conrad Foo, Alma Andersson, Brandon D Kayser, Yiming Yang, Alsu Missarova, Priyanka Kulkarni, Jan-Christian Huetter, Sharmila Chatterjee, Youngmi Kim, et al. Spatial decoding of tertiary lymphoid structure maturation in non-small cell lung cancer using deep neural networks. bioRxiv, pages 2026–01, 2026.

[26] Anna Pascual-Reguant, Ralf Köhler, Ronja Mothes, Sandy Bauherr, Daniela C Hernández, Ralf Uecker, Karolin Holzwarth, Katja Kotsch, Maximilian Seidl, Lars Philipsen, et al. Multiplexed histology analyses for the phenotypic and spatial characterization of human innate lymphoid cells. Nature communications, 12(1):1737, 2021.

[27] F Alexander Wolf, Fiona K Hamey, Mireya Plass, Jordi Solana, Joakim S Dahlin, Berthold Göttgens, Nikolaus Rajewsky, Lukas Simon, and Fabian J Theis. Paga: graph abstraction reconciles clustering with trajectory inference through a topology preserving map of single cells. Genome biology, 20(1):59, 2019.

[28] Florian J Weisel, Griselda V Zuccarino-Catania, Maria Chikina, and Mark J Shlomchik. A temporal switch in the germinal center determines differential output of memory b and plasma cells. Immunity, 44(1):116–130, 2016.

[29] David Tarlinton and Kim Good-Jacobson. Diversity among memory b cells: origin, consequences, and utility. Science, 341(6151):1205–1211, 2013.

[30] Tommaso Biancalani, Gabriele Scalia, Lorenzo Buffoni, Raghav Avasthi, Ziqing Lu, Aman Sanger, Neriman Tokcan, Charles R Vanderburg, Åsa Segerstolpe, Meng Zhang, et al. Deep learning and alignment of spatially resolved single-cell transcriptomes with tangram. Nature methods, 18(11):1352–1362, 2021.

[31] Paolo Cadinu, Kisha N Sivanathan, Aditya Misra, et al. Charting the cellular biogeography in colitis reveals fibroblast trajectories and coordinated spatial remodeling. Cell, 187(8):2010–2028, 2024.

[32] Eóin N McNamee and Jesús Rivera-Nieves. Ectopic tertiary lymphoid tissue in inflammatory bowel disease: protective or provocateur? Frontiers in immunology, 7:308, 2016.

[33] Lianyu Zhao, Song Jin, Shengyao Wang, Zhe Zhang, Xuan Wang, Zhanwei Chen, Xiaohui Wang, Shengyun Huang, Dongsheng Zhang, and Haiwei Wu. Tertiary lymphoid structures in diseases: immune mechanisms and therapeutic advances. Signal transduction and targeted therapy, 9(1):225, 2024.

[34] Eóin N McNamee, Joanne C Masterson, Paul Jedlicka, Colm B Collins, Ifor R Williams, and Jesús Rivera-Nieves. Ectopic lymphoid tissue alters the chemokine gradient, increases lymphocyte retention and exacerbates murine ileitis. Gut, 62(1):53–62, 2013.

[35] Yuki Sato, Karina Silina, Maries Van Den Broek, Kiyoshi Hirahara, and Motoko Yanagita. The roles of tertiary lymphoid structures in chronic diseases. Nature Reviews Nephrology, 19(8):525–537, 2023.

[36] Abbas Nazir, Hanchen Wang, Ziyu Lu, et al. Spatiotemporal profiling reveals the role of inflammatory niche in driving prostate cancer. bioRxiv, 2026.

[37] Chris Lu, Cong Lu, Robert Tjarko Lange, Yutaro Yamada, Shengran Hu, Jakob Foerster, David Ha, and Jeff Clune. Towards end-to-end automation of ai research. Nature, 651(8107):914–919, 2026.

[38] Martiño Ríos-García, Nawaf Alampara, Chandan Gupta, Indrajeet Mandal, Sajid Mannan, Ali Asghar Aghajani, NM Krishnan, and Kevin Maik Jablonka. Ai scientists produce results without reasoning scientifically. arXiv preprint arXiv:2604.18805, 2026.

[39] Ziming Luo, Atoosa Kasirzadeh, and Nihar B Shah. The more you automate, the less you see: Hidden pitfalls of ai scientist systems. arXiv preprint arXiv:2509.08713, 2025.

[40] CZI Single-Cell Biology Program, Shibla Abdulla, Brian Aevermann, et al. Cz cell× gene discover: A single-cell data platform for scalable exploration, analysis and modeling of aggregated data. BioRxiv, pages 2023–10, 2023.

[41] Oscar Franzén, Li-Ming Gan, and Johan Björkegren. Panglaodb: a web server for exploration of mouse and human single-cell rna sequencing data. Database, 2019:baz046, 2019.

[42] Congxue Hu, Tengyue Li, Yingqi Xu, et al. Cellmarker 2.0: an updated database of manually curated cell markers in human/mouse and web tools based on scrna-seq data. Nucleic acids research, 51(D1):D870–D876, 2023.

[43] Ilya Korsunsky, Nghia Millard, Jean Fan, Kamil Slowikowski, Fan Zhang, Kevin Wei, Yuriy Baglaenko, Michael Brenner, Po-ru Loh, and Soumya Raychaudhuri. Fast, sensitive and accurate integration of single-cell data with harmony. Nature methods, 16(12):1289–1296, 2019.

[44] Simon Mages, Noa Moriel, Inbal Avraham-Davidi, Evan Murray, Jan Watter, Fei Chen, Orit Rozenblatt-Rosen, Johanna Klughammer, Aviv Regev, and Mor Nitzan. Tacco unifies annotation transfer and decomposition of cell identities for single-cell and spatial omics. Nature Biotechnology, 41(10):1465–1473, 2023.

